# Phylogenetic Estimation of branch-specific Shifts in the Tempo of Origination

**DOI:** 10.1101/2025.05.28.656596

**Authors:** Bjørn T. Kopperud, Sebastian Höhna

## Abstract

Studying rates of species diversification is one of the key themes in macroevolution. In particular, we are interested in if some clades in a phylogeny diversify more rapidly/slowly than others due to branch-specific diversification rates. A common approach in neontological studies is to use a phylogenetic birth-death process to model species diversification. Specifically, the birth-death-shift process is used to model branch-specific shifts in the tempo of diversification. Here, we present `Pesto`, a new method and software for estimating branch-specific diversification rates that does not rely on Markov chain Monte Carlo simulations for fitting the model and instead deterministically computes the posterior mean branch-specific diversification rates using only two traversals of the tree. This method is blazingly fast: the birth-death-shift model can be fitted to large phylogenies (*>* 20k taxa) in minutes on a personal computer while also providing branch-specific inference of diversification rate shift events. Thus, we can robustly infer branch-specific diversification rates and the number of diversification rate shift events for large-scale phylogenies as well as exploring the characteristic of the birth-death-shift model through complex and large-scale simulations. Here, we first describe the method and the software implementation `Pesto` and explore its behavior using trees simulated under the birth-death-shift model. Then, we explore the behavior of inferring significant branch-specific diversification rate shifts using both Bayes factors and effect sizes. We find few to none false positive inferences of diversification rate shift events but many false negatives (reduced power). The most difficult parameter to estimate is the rate at which diversification rate shifts occur (shift rate). Despite this, branch-specific diversification rate estimates are precise and nearly unbiased.

## Introduction

Estimating rates of diversification is one of the key interests in macroevolutionary biology and paleontology. Diversification rates are fundamental to our understanding of how historical biodiversity has changed. Recent developments in the field have focused on birth-death models that describe the tempo of lineage diversification (Ricklefs, 2007; Morlon, 2014). This process is highly variable; diversification rates may vary over time with episodes of mass extinctions (Raup and Sepkoski, 1982) or elevated speciation rates (Sepkoski, 1998) and vary among branches with clades in high or low turnover (Sanderson and Donoghue, 1994, 1996; Jetz et al., 2012). Moreover, we can speculate that diversification rate variation over time is associated with climatic or environmental factors (Hua and Wiens, 2013; Condamine et al., 2018; Palazzesi et al., 2022) whereas diversification rate variation among branches is associated with, amongst others, key innovations or changes in habitat (Hodges and Arnold, 1995; Allen and Gillooly, 2006; Weir and Schluter, 2007; Silvestro et al., 2014). Thus, we are ultimately interested in linking diversification rates to, for example, underlying ecological and environmental factors (Condamine et al., 2018). However, as a first step, we should infer diversification rates independently to learn if there was any diversification rate variation for the specific study group (Budd and Mann, 2024). Here, we will not consider branch-specific shifts in diversification rates in relation to the evolution of novel characters or changes in ecosystems, however we are nonetheless interested in identifying the branch-specific diversification rate shifts themselves. In particular, we are interested in where on the phylogeny the diversification rates shifted and the size of the shifts, in order to test hypotheses about the generation of biodiversity.

In recent years, several statistical approaches were developed to estimate branch-specific diversification rates from phylogenies (`BAMM`, Rabosky 2014; `ClaDS`, Maliet et al. 2019; `LSBDS`, Höhna et al. 2019; `MTBD`, Barido-Sottani et al. 2020; `MiSSE`, Vasconcelos et al. 2022; `RPANDA` Mazet et al. 2023; see Martínez-Gómez et al. 2024 for a review). One key limitation of existing methods is that they are prohibitively slow. Existing methods rely on simulation techniques (i.e., Markov-chain Monte Carlo simulations together with data augmentation or stochastic mapping) with long run times (i.e., often days to weeks). To complicate matters, these highly complex simulation algorithms are slow to converge and can often fail (Martínez-Gómez et al., 2024). Thus, current methods are (i) impractical for phylogenies that contain many tips (i.e., *>* 10, 000) which should be of main interest to study diversification rate variation (Rabosky et al., 2018), and (ii) too computationally demanding to explore model assumptions and behavior via large-scale simulation studies. We solve the computational bottleneck by developing a novel approach that deterministically computes branch-specific diversification rates on a given phylogeny. Thus, our approach does not suffer from convergence issues. Our approach uses the same underlying birth-death-shift process as developed by Barido-Sottani et al. (2020) and Höhna et al. (2019) and inherits its biological assumptions and statistical properties. Therefore, our new approach and the implementation in `RevBayes` (Höhna et al., 2016, 2019) give identical estimates of the posterior mean branch-specific diversification rates, if the same parameters (i.e., diversification rate categories and shift rate) and model assumptions are chosen. However, our new approach is orders of magnitude faster.

An outstanding question is whether we can reliably infer the number and location of diversification rate shifts, or if we can only reliably estimate branch-specific diversification rates (Shi and Rabosky, 2015; Moore et al., 2016; Mitchell and Rabosky, 2017; Höhna et al., 2019). A change in diversification rates could either be explained by several small or a few large shifts. Therefore, the number of estimated diversification rate shifts depends strongly on the prior assumption on the shift rate (Moore et al., 2016; Höhna et al., 2019). Nevertheless, hypothesis testing using Bayes factors can be used to assess the support for alternative hypotheses despite sensitivity to the prior (Kass and Raftery, 1995). In this study we develop an efficient approach to infer the number and location of diversification rate shifts on phylogenies using Bayes factors (see also Shi and Rabosky, 2015).

Our approach for inferring branch-specific diversification rates and branch-specific rate shift events is inspired by Bayesian inference of ancestral discrete characters (Yang et al., 1995; Nielsen, 2002). However, we discovered that the probabilistic description of the birth-death-shift model is incompatible with established theory on Bayesian belief networks (Pearl, 1988). Specifically, we are not able to represent branch-specific diversification rates as random variables that are separable from the phylogeny (see Fig. S1). Thus, if we assume the phylogeny as “observed” data, then the branch-specific diversification rates are “unobserved” data instead of parameters of the model. This is an issue, as the prior probability and the likelihood of branch-specific diversification rates are conflated (see also May and Rothfels, 2023). We suspect that the inseparability of prior and likelihood is also relevant for previous approaches that used Bayesian inference to fit identical (Höhna et al., 2019; Barido-Sottani et al., 2020) or very similar stochastic processes (Rabosky, 2014; Maliet et al., 2019; Quintero et al., 2024). Nevertheless, we conceptualize our approach using ideas and terminology from Bayesian inference theory (for more discussion see Supplementary Material Section S1).

We provide a `Julia` implementation of our new algorithm called *Phylogenetic Estimation of Shifts in the Tempo of Origination (*`Pesto`). The algorithm requires only two tree traversals: one postorder (from the tips to the root) and one preorder (from the root to the tips) tree traversal. The first pass over the tree is used to compute the likelihood scores and the second pass is used to compute and map the posterior mean branch-specific diversification rates. We show the time complexity of the algorithm for various tree sizes, demonstrating that `Pesto` infers robust branch-specific diversification rates and diversification rate shifts for phylogenies with *>* 10, 000 taxa in a few minutes on a standard personal computer. We validate `Pesto` by showing that branch-specific diversification rate estimates are identical to estimates obtained using `LSBDS` (Höhna et al., 2019), up to some Monte Carlo error, when the identical model is used. Furthermore, we explore the behavior of `Pesto` using a simulation study and show that parameter estimation is robust given sufficiently large phylogenies. For example, estimates of the shift rate are less reliable when there are few shifts, i.e., if shifts are rare and/or if the phylogeny is small. Branch-specific diversification rates are robustly inferred even for small trees (minimum 100 taxa). Finally, we present a simulation study to explore the ability to estimate the number and location of diversification rate shifts. `Pesto` is conservative in the inference of diversification rate shift events with virtually no false positives, but has low power (i.e., more false negative shift event inferences).

## Materials and methods

### The Underlying Theory of Phylogenetic Estimation of Shifts in the Tempo of Origination (PESTO)

Our new method *Phylogenetic Estimation of Shifts in the Tempo of Origination (*`Pesto`) relies on and extends existing methods for estimating branch-specific diversification rates. The underlying stochastic process in `Pesto` is equivalent to the *birth-death-shift* process (Höhna et al., 2019). The existing implementations of the birth-death-shift process and similar processes use computationally expensive techniques such as Markov chain Monte Carlo simulation with data augmentation (Rabosky, 2014; Maliet et al., 2019; Barido-Sottani et al., 2020) or stochastic mapping (Höhna et al., 2019) to infer branch-specific diversification rates. The fundamental novelty of `Pesto` is our deterministic algorithm for calculating the posterior mean branch-specific diversification rates and the number of diversification rate shifts, conditional on the specific choice of diversification rate categories and shift rate. Unlike with simulation approaches, the calculations are performed only once and there is no randomness or inherent uncertainty in the inference procedure. Hence, we can run analyses in `Pesto` using trees that have tens of thousands of tips in a matter of minutes on a personal computer. We describe the details of the method in the following sections.

### The birth-death-shift model

`Pesto` uses the same underlying birth-death-shift process as described by Höhna et al. (2019), which we recapitulate here. The birth-death-shift process has a speciation rate *λ*, an extinction rate *µ*, as well as a shift rate *η*. Suppose one begins a tree simulation with a starting (root) node that represents the common ancestor of the clade, and that the root node has two descendant branches of unknown lengths. After a waiting time modeled as an exponentially distributed random variable (with rate *λ* + *µ* + *η*), there are three possible events. First, there can be a speciation event, with probability *λ/*(*λ* + *µ* + *η*), where two new branches are created. Second, there can be an extinction event, with probability *µ/*(*λ* + *µ* + *η*), where the branch terminates. Third, there can be a rate shift event, with probability *η/*(*λ* + *µ* + *η*). At a rate-shift event, a new set of diversification rates (i.e. the pair *λ*^*′*^, *µ*^*′*^) is chosen randomly, which we will explain in the following section. Note that we assume that the new diversification rates are independent of the old diversification rates (unlike Maliet et al., 2019; Quintero et al., 2024). After some time *t* has passed, the tree has either gone entirely extinct or it has survived. If the process survived, then each species is sampled with probability *ρ*. Finally, extinction events and unsampled species are pruned, resulting in a so-called “reconstructed tree”, meaning that the tree appears as if it was reconstructed from molecular or morphological data at the present.

### Base distribution of diversification rates

When a diversification rate shift event occurs in the birth-death-shift process, new diversification rates are drawn randomly from a base distribution. For example, `BAMM` uses an exponential base distribution (Rabosky, 2014) and Höhna et al. (2019) proposed to use instead a log-normal distribution. However, any non-negative base distribution could in principle be used. Thus, in addition to the exponential and the log-normal base distributions, we also explored the use of a log-uniform and a gamma base distribution. We parameterized the base distributions in terms of their means and variances where applicable. Specifically, we specified the exponential distribution with a mean of 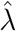, the log-normal distribution with a median of 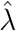 and standard deviation of *H* ≈ 0.587 (meaning that the 2.5%–97.5% quantile interval of the base distribution spans one order of magnitude), and the log-uniform with a mean of 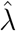 and a span of one order of magnitude. For the gamma distribution, we set the scale and shape parameters such that the mean was 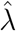, and the variance was equal to that of the log-normal distribution. This results in four base distributions whose location is controlled by the 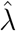-parameter, and the variance is either fixed or also determined by the 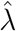 parameter. We will discuss specific choices of 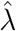 and the variance later in section *Estimating the parameters of the birth-death-shift model*.

### Discretization of diversification rates

Simulating a phylogeny under the birth-death-shift process with continuous base distributions is relatively straight forward. Moreover, modeling the base distributions as being fully continuous might be the preferable approach for inference. However, it is not trivial to calculate the probability of observing the phylogeny when using continuous base distributions. For the probability calculation to be computationally tractable, we follow the birth-death-shift implementation of the `LSBDS` model (Höhna et al., 2019) where an approximation approach is used.

Specifically, the continuous base distributions are discretized by splitting them into *n* bins with equal probability mass (i.e., the bins represent quantiles). This results in a vector of diversification rate *classes*, i.e.,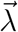 for speciation rates, and 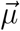 for extinction rates. Next, we select all pairwise combinations of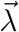 and 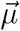 to form the vectors ***λ*** = [*λ*_1_, *λ*_2_, …, *λ*_*K*_] and ***µ*** = [*µ*_1_, *µ*_2_, …, *µ*_*K*_], which represent the *n*^2^ = *K* diversification rate *categories* (see Fig. 1). Choosing a higher number of rate classes will result in a better approximation of the base distributions (Höhna et al., 2019). In the limit of infinitely many rate classes, the birth-death-shift process with discrete and continuous base distributions for the diversification rates are equivalent. Therefore, we expect that inferences made using the discrete base distribution should converge and stabilize as more bins are used, however at the cost of increased computational time.

**Fig. 1:**
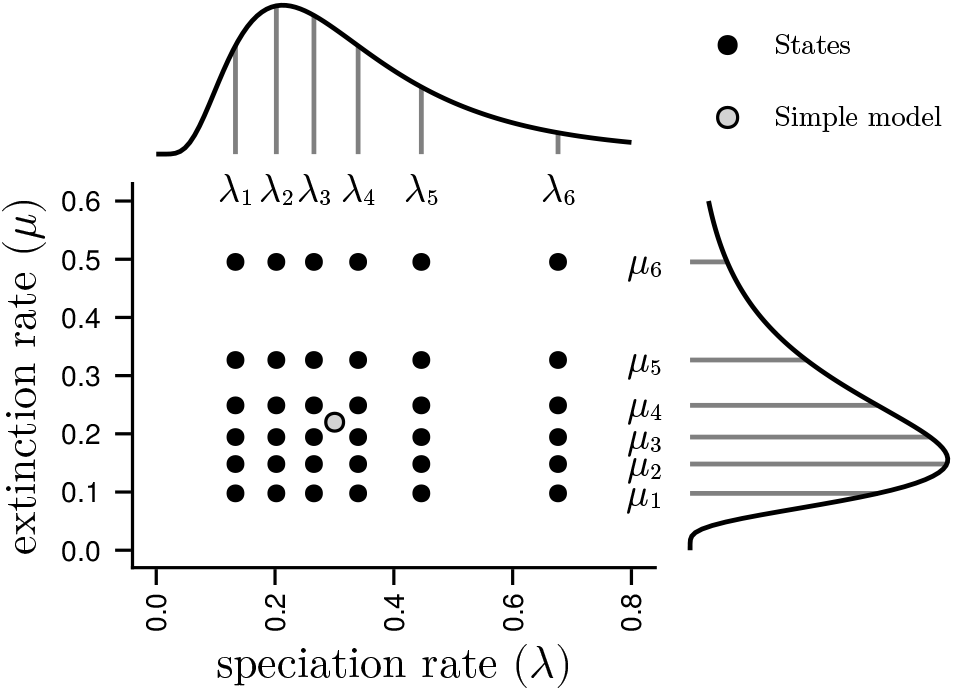
Schematic of diversification rate categories. First, Pesto estimates the speciation rate 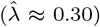 and the extinction rate 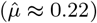 under the constant-rate birth-death process (gray dot). These constant-rate estimates are used as the mean of base distributions. Second, we select the number of quantiles (*n* = 6) and the standard deviation *H* = 0.587 chosen such that the 2.5%–97.5% quantile interval spans one order of magnitude. Third, we consider all pairwise combinations of the quantiles (black dots, *K* = *n*^2^ = 36). The log-normal distribution with *n* = 6 quantiles and a variance of *H* is the default option, however other distributions and other number of quantiles and variances can be specified. The rate category values can also be specified manually. Each black dot is a rate category in the birth-death-shift model, and together they represent the range of rate heterogeneity allowed under the model. Along each branch and across time, we calculate the probability that the process was in each rate category using Eq. 7.

### Computing the likelihood of the phylogeny

In this section we will explain how we compute the probability density of observing the phylogeny, as well as how we set up the likelihood function. While there are no known analytical solutions for the probabilities and probability densities, it is possible to find solutions using numerical methods. The idea is to express the change in probability (density) by considering which events (i.e., speciation, extinction or rate shift events) are possible in a small time interval Δ*t*, and taking the limit as Δ*t* → 0, in other words using differential equations (see Maddison et al., 2007; FitzJohn, 2012, for more details on how the differential equations are derived). As the data, we assume a rooted, bifurcating phylogenetic tree with branch lengths in units of time. We also assume that the tree is known without error. Note that the likelihood function (Eq. 4) is analogous to the likelihood function of a state-dependent birth-death process without observed states (Maddison et al., 2007; FitzJohn, 2012; Beaulieu and O’Meara, 2016; Freyman and Höhna, 2018) and we repeat it here as we require the likelihood function and the probability density computations to derive our algorithm to deterministically compute the posterior mean branch-specific diversification rates.

Conceptually, our algorithm contains the following steps. First, we calculate the extinction probabilities (**E**(*t*)), which are invariant to the topology and divergence times. Second, we calculate the probability density of observing the tree (**D**(*t*)). Since the probability density depends on the topology and divergence times, it needs to be computed for each branch in a postorder tree traversal. Third, we use the probability density of observing the tree to set up the likelihood function. Note that we write scalars in plain typeface and vectors and matrices in bold. We treat vectors as matrices with a single column and transposed vectors as matrices with a single row. See table S1 for an overview of the notation and symbols that are used.

### Computing extinction probabilities

In order to calculate the extinction probabilities, we use the following system of differential equations (Maddison et al., 2007):

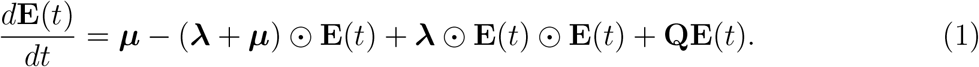

More precisely, *E*_*j*_(*t*) represents the probability that a lineage that was in rate category *j* at time *t* went extinct before the present or was not sampled in the phylogeny. Note that we use the symbol ⊙ to represent element-wise multiplication. The matrix **Q** has entries −*η* on the diagonal, and entries *η/*(*K* − 1) elsewhere. For example in the case of a three-category model, **Q** would be

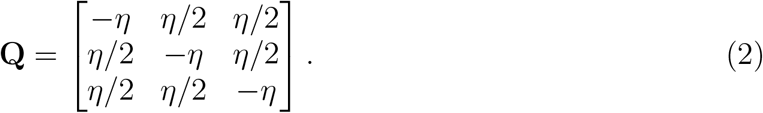

We allow for the possibility that the taxa are incompletely sampled, but we assume an equal probability of taxon sampling (FitzJohn, 2010; Höhna et al., 2011). Therefore, the initial values of the extinction probabilities are all set to *E*_*j*_(*t* = 0) = 1 − *ρ*, where *ρ* is the known/fixed taxon sampling fraction.

### Computing the probability density of the observed branches

The probability of observing the branch (or subtree) at time *t* is likewise represented as a set of differential equations (Maddison et al., 2007; FitzJohn, 2012):

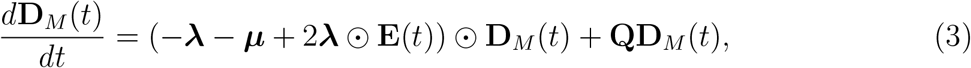

where *M* is the branch index. Specifically, if *M* is a terminal branch, then *D*_*M,j*_(*t*) is the probability of observing the branch if the process was in category *j* at time *t*. If *M* is an internal branch, then *D*_*M,j*_(*t*) represents the probability density of observing the subtree descended from branch *M*. For initial values, we assign *D*_*M,j*_(*t* = 0) = *ρ* for all categories *j* if *M* is a terminal branch. For internal branches, the initial (youngest) value of a branch *M* is assigned to **D**_*M*_ (*t*) := **D**_*A*_(*t*) ⊙ **D**_*B*_(*t*) ⊙ ***λ***, where the branches *A* and *B* are the descendants of branch *M*. We iterate and solve **D**_*M*_ (*t*) for all branches *M* in a postorder traversal of the tree (Fig. 2**a–c**) until the root node (**D**_*R*_(*t*)) is reached.

**Fig. 2:**
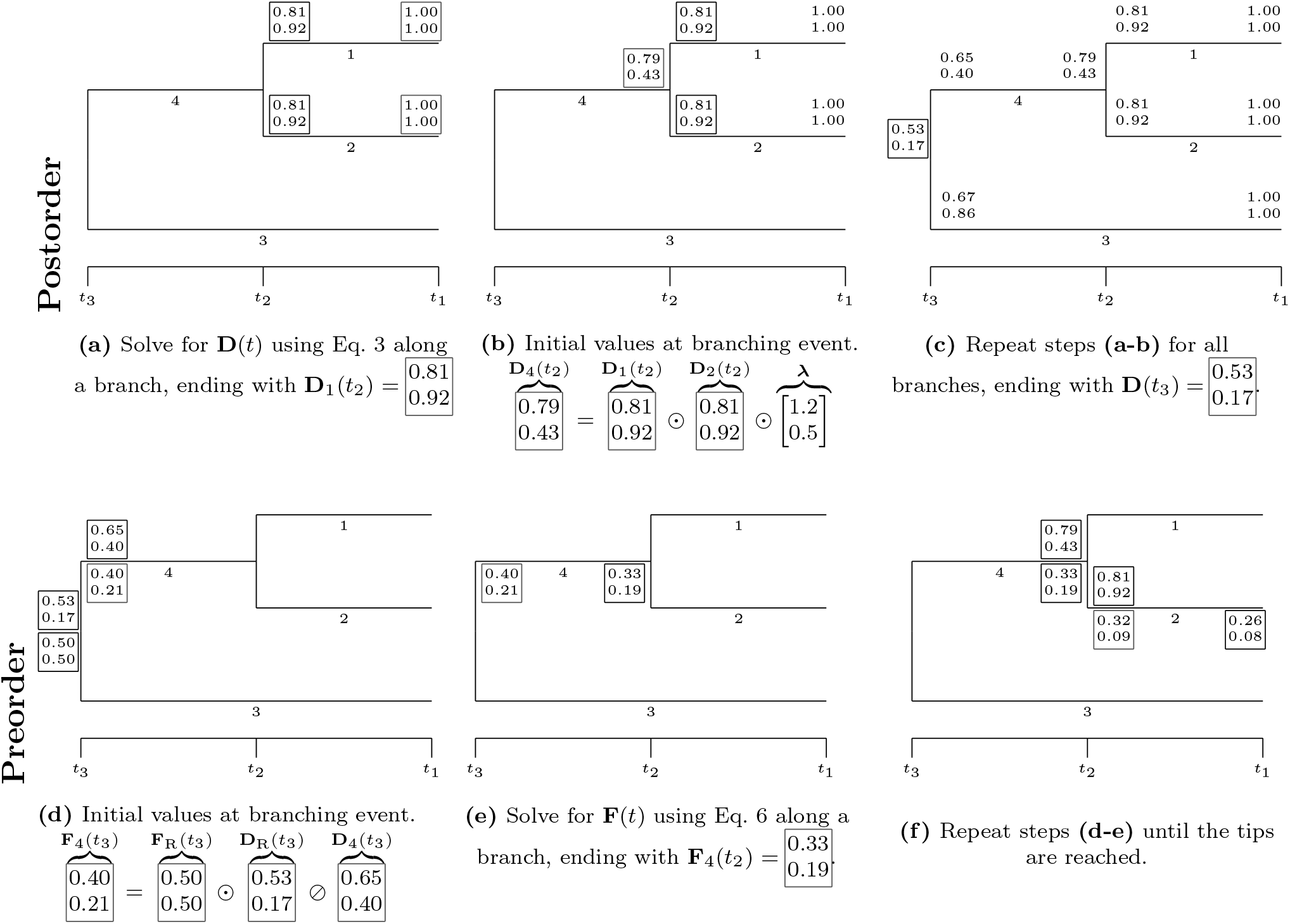
The algorithm for the two tree traversals on a miniature tree. The postorder pass **(a–c)** computes the probabilities given the descendants (**D**(*t*), written above the branches). The preorder pass **(d–f)** computes the probabilities given the tree up to but not including the descendants (**F**(*t*), written below the branches). The white boxes on the trees correspond to the white boxes in the caption. We set **D**(*t*_1_) = [1.0, 1.0] at the tips, and **F**_*R*_(*t*_3_) = [0.5, 0.5] at the root. The densities **D**(*t*) and **F**(*t*) are used to calculate the probability that the process was in a particular rate category at a particular time, see Eq. 7. In this example, we used divergence times *t*_1_ = 0, *t*_2_ = 0.1, *t*_3_ = 0.2, speciation rates ***λ*** = [1.2, 0.5], extinction rates ***µ*** = [1.0, 0.3], shift rate *η* = 0.01, extant sampling fraction *ρ* = 1, we did not condition on survival, and we did not re-scale the initial values.

### Computing the probability of observing the tree (likelihood)

The likelihood function for the birth-death-shift process is defined as the weighted average of the probabilities at the root node (Eq. 3) over all diversification rate categories. When calculating the likelihood of observing the tree, we condition on (i) that the two branches subtending from the root survived until the present, and (ii) that there was a speciation event at the root:

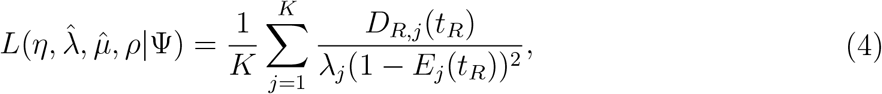

where *t*_*R*_ is the age of the root, Ψ represents the phylogeny (i.e., topology and branch lengths), and we follow Pagel (1994) in using a flat prior probability (1*/K*) of being in any of the diversification rate categories at the root (see also Goldberg and Igićc, 2008; FitzJohn et al., 2009; FitzJohn, 2012, for alternative root treatments).

### Estimating the parameters of the birth-death-shift model

Höhna et al. (2019) argued for estimating the shift rate η jointly with the parameters of the base distributions (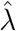 and 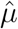) using Bayesian inference while Rabosky et al. (2014) advocates using an empirical Bayes approach to estimate the prior means of the base distributions under a constant rate birth-death process and the shift rate *η* is replaced with a geometric distribution directly on the number of shift events (Mitchell and Rabosky, 2017). We discovered, however, that estimating the base distribution parameters and the shift rate jointly is more difficult than previously thought. The joint likelihood surface includes many local optima and a hill-climber algorithm can easily get stuck in these local optima (Fig. S6). As a compromise, we settled for a two-step approach (Fig. 1). This involves first finding the maximum-likelihood estimates for 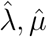 using the simpler lineage-homogeneous constant-rate birth-death model (as done in Rabosky et al., 2014). Second, we infer η by maximizing the likelihood of the birth-death-shift model while keeping the other parameters fixed

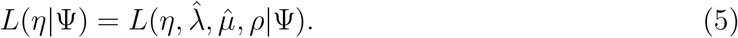

This is also called an estimated likelihood function (Pawitan, 2001, p. 274), with which we did not encounter any local optima issues. We note, however, that the estimated base distribution parameters and shift rate using the two-step approach are not necessarily the same as the true maximum likelihood estimates, and the optimal approach for selecting or estimating the diversification rate base distribution and shift rate requires further research.

### The posterior distribution of rate shift histories

It is not possible to directly observe the true rate shift history on a phylogeny except in the case of simulated phylogenies. When we wish to make inferences about the birth-death-shift process that led to an “observed” phylogeny, we consider the diversification rate shift history to be an unobserved random variable. As we can not observe it directly, we calculate the likelihood by considering all possible diversification rate shift histories that could have led to the phylogeny in question (Eq. 4). In other words, when calculating the likelihood, we integrate across all possible realizations of the diversification rate shift history. By doing so, however, we are omitting several aspects of interest. For example, we are interested in the branch-specific diversification rates and how many diversification rate shift events occurred, across branches and across time. Therefore, we wish to investigate the posterior distribution of diversification rate shift histories, i.e., the probability distribution of the diversification rate shift histories conditioned on the data. Höhna et al. (2019) showed how one could approximate the posterior distribution of the diversification rate shift histories using stochastic mappings similar to stochastic character mapping for discrete character evolution (Huelsenbeck et al., 2003). By simulating many samples of the posterior distribution of diversification rate shift histories, one can compute summaries such as the posterior mean branch-specific diversification rate, the posterior mean number of diversification rate shift events, or the posterior probability that there was at least one diversification rate shift. In order for this sampling approach to yield a good approximation of the posterior distribution, one must i) simulate with small step sizes in time, and ii) repeat the stochastic mapping thousands or millions of times. Accordingly, one of the main disadvantages of the stochastic mapping approach is its slow run time and computational burden, although it is more efficient than data augmentation (Höhna et al., 2019).

In the next sections, we will outline a deterministic algorithm that gives equivalent summaries of the posterior distribution. Instead of using stochastic mappings, this algorithm is entirely non-random (i.e., it is deterministic), and thus will yield the same results every time the algorithm is executed. Notably, we will compute i) the posterior probability that the process was in a particular diversification rate category, ii) the posterior mean branch-specific diversification rates, iii) the posterior mean number of branch-specific rate shift events, iv) the posterior probability that there was at least one rate shift event at a given branch, and v) the Bayes factor for the support of the hypothesis that there was at least one rate shift event at a given branch. These calculations are facilitated by simple formulae or differential equations that need to be solved only once per branch. Compared with stochastic mapping, our approach is much more computationally efficient and consequently the inferences can be made in a fraction of the time.

### Performing a preorder traversal to calculate posterior rate category probabilities

We use a dynamic programming approach to compute the posterior probabilities of the rate categories (Fig. 3), specifically the backward-forward algorithm (Rabiner, 1989) extended to accommodate a tree structure (Pearl, 1982). The motivation for applying the backward-forward algorithm is based on the fact that the birth-death-shift process can along a lineage be seen as a hidden Markov model where the diversification rate categories are the hidden states. The backward pass of the algorithm was explained previously, where we calculated the probabilities **D**_*M*_ (*t*). In the forward pass, we calculate the forward probabilities **F**_*M*_ (*t*). Specifically, *F*_*M,j*_(*t*) represents the probability that the rate category was *j* on branch *M* at time *t*, given the part of the phylogeny that is not descended from branch *M* at time *t*. At the root node *R*, there is no ancestral node, meaning that *F*_*R,j*_(*t*_*R*_) represents the prior probability of the rate category being *j*. We assume that the ancestral diversification rate is a random variable that is distributed according to the (discretized) base distribution. Therefore, we initialize *F*_*R,j*_(*t*) to 1*/K* at the root node (recall that each diversification rate category represents quantile intervals with equal probability mass). When initializing *F*_*R,j*_(*t*), we also condition on i) survival and ii) that there was a speciation event at the root, in the same manner as in the likelihood function (Eq. 4). Since the rate category variable is the only random variable (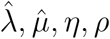 are fixed to their estimates), there are no other priors (see Figure S1). In order to solve for **F**_*M*_ (*t*) along the branches, we use the following set of differential equations

**Fig. 3:**
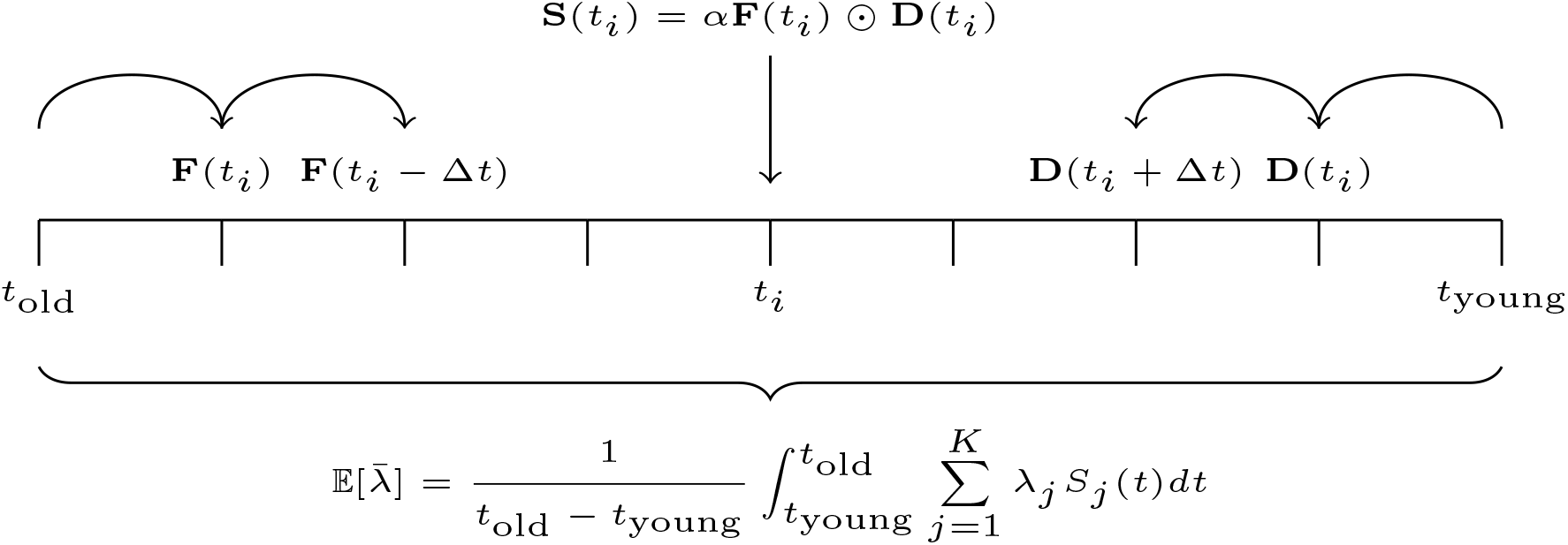
A simplified schematic of the backward-forward algorithm along a branch. First, we find the solution for **D**(*t*), by starting from *t*_young_ and going towards the past. Second, we find the solution for **F**(*t*), by starting from *t*_old_ and going towards the present. Third, we calculate the ancestral rate category probabilities **S**(*t*), which is **F**(*t*) element-wise times **D**(*t*), normalized by *α* such that **S**(*t*) sums to one for any particular time *t*. Fourth, we compute the posterior mean speciation rate, averaged across the time interval of the branch 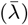. Note that the fixed time intervals Δ*t* are plotted for illustrative purposes. In the actual implementation, the number of time intervals, and the size of the time intervals can vary.

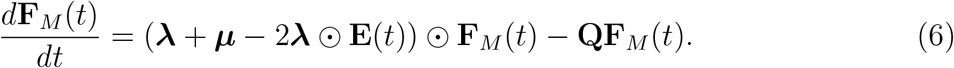

Note that *d***F***/dt* is equal to *d***D***/dt*, except the sign has changed on both the left and the right sides. The reason for the change in sign is that we are solving *d***F***/dt* in forward time (from ancient to recent), and therefore the time differential *dt* is negative.

Once we have solved **F**_*M*_ (*t*), we compute posterior probabilities of the rate categories (see the supplementary material for a discussion on the posterior). They are given as follows

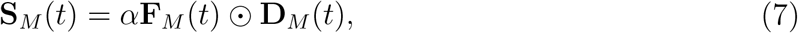

where *α* is a normalizing factor such that **S**_*M*_ (*t*) sums to one (Fig. 3, see Pearl 1982, 1988). In summary, the preorder traversal consists of two steps. First, we initialize **F**_*M*_ (*t*) := **S**_*P*_ (*t*) ⊘ **D**_*M*_ (*t*), where *P* is the parental branch, *t* is the divergence time, and ⊘ represents element-wise division. Second, we solve **F**(*t*) along branch *M*. We repeat these two steps for all descendant branches, in a preorder traversal of the tree (**Fig. 2d–f**). When the preorder traversal is completed, the backward-forward algorithm is also completed.

### Estimating the posterior mean branch-specific diversification rates

Estimating branch-specific diversification rates is a key result for any method that fits the birth-death-shift process to a phylogeny. We consider the diversification rate shift history and thus also the branch-specific diversification rates as random variables. For a given branch *M* and a time *t*, the moments of the speciation rate *λ*_*M*_ (*t*) are weighted averages of the speciation rate categories *λ*_*j*_, with the weights given by their posterior probabilities, which we can use to compute the posterior mean and variance

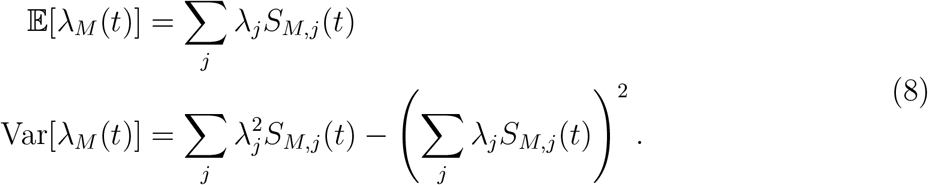

This can for example be used to compute tip rates, where the posterior mean speciation rates are computed at the present. To summarize over a branch as a whole, we use the symbol 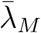, and get the expectation by integrating over the time span and dividing by the length of the branch:

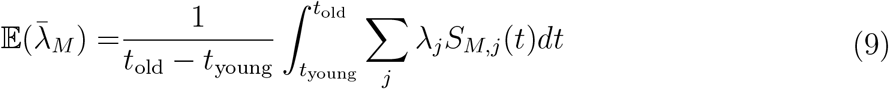

meaning that 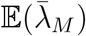 is the posterior mean speciation rate for branch *M*, and *t*_young_, *t*_old_ represent the youngest and oldest times of branch *M* (Fig. 3). If one is interested in the variance of 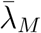, it is possible to approximate the integral by splitting the branch into *c* discrete bins represented by *λ*(*t*_1_), *λ*(*t*_2_), … etc. Then, we can apply the standard formula for the variance of a linear combination of correlated random variables

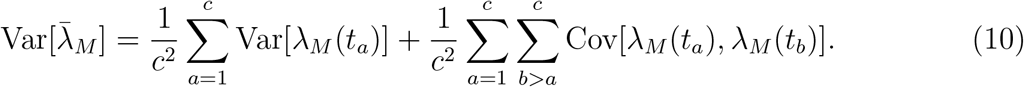

Since the time bins are not independent, it is necessary to calculate the covariance, which one can calculate as

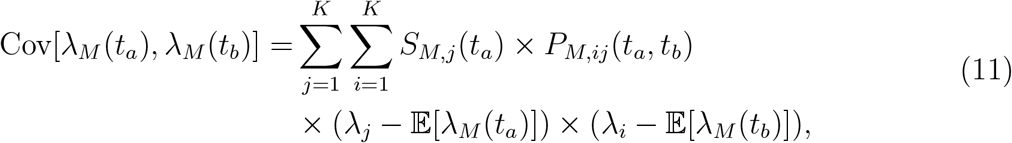

where *S*_*M,j*_(*t*_*a*_) is the marginal probability of the diversification rate category being *j* at time *t*_*a*_, and *P*_*M,ij*_(*t*_*a*_, *t*_*b*_) is the transition probability from category *j* at time *t*_*a*_ to category *i* at a younger time *t*_*b*_, normalized such that *P*_*M,ij*_(*t*_*a*_, *t*_*b*_) sums to one over the departure categories *j*. We note that computing the variance of the branch-specific speciation rates is significantly more computationally expensive than computing the mean. For more details on how to calculate the posterior variance of the speciation rate, see the supplementary material.

If one is interested in the posterior mean and variance of the branch-specific extinction rate, net-diversification rate, or relative extinction rate, one can substitute *λ*_*j*_ for *µ*_*j*_, *λ*_*j*_ − *µ*_*j*_, or *µ*_*j*_*/λ*_*j*_, respectively, and use the same formulae. In order to evaluate the integrals, we use Gauss-Legendre quadrature with *n* = 10 points. We want to emphasize that previous methods (Freyman and Höhna, 2019; Höhna et al., 2019) used Eq. 7 to simulate the branch-specific diversification rates (i.e., stochastic mappings), whereas *Pesto* uses Eq. 9 to compute the posterior mean deterministically. To present the results, we plot the posterior mean branch-specific diversification rates on the tree, however standard visualizations like scatter plots or histograms can also be used.

### Estimating the number of rate shifts

Another key property of the birth-death-shift process that we are interested in is the number of diversification rate shift events that occurred across a phylogeny. As for the branch-specific diversification rates, the number of diversification rate shifts for a particular branch is also a random variable, which we will summarize by computing the posterior mean. The idea is to compute the average number of diversification rate shifts in a small time interval Δ*t*, by computing the probability of a rate shift from rate category *j* to a different rate category *i*, and multiplying by the posterior probability that the process was in rate category *j* at the time. By summing the number of diversification shifts over several small time intervals Δ*t*_1_, Δ*t*_2_, … across the branch, we can calculate the total number of rate shifts across the time span of the branch. In the limit of Δ*t* → 0, the number of rate shifts converge to zero, however the change in the average number of rate shifts per time can still be expressed as a differential equation:

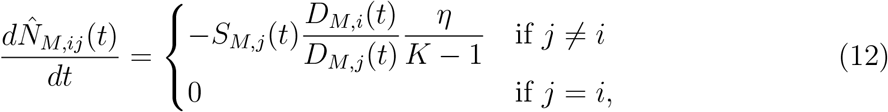

where 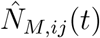 is the posterior mean number of accumulated rate shifts from a diversification rate category *j* to another diversification rate category *i* from the beginning of branch *M* until a younger time *t*. The initial value is 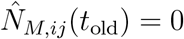, and the time steps *dt* are negative. For the posterior mean number of diversification rate shifts to be of any note (i.e., non-zero), two conditions must be met. First, the posterior probability that the process was in the departure category *j* at time *t* must be non-zero. Second, the probability density (of observing the tree, given the process was in the rate category) of the arrival category *i* must be equal or larger than that of the departure category *j* (i.e., *D*_*M,i*_(*t*) ⩾ *D*_*M,j*_(*t*)). For details on how the expression is derived, see the supplementary material. Since we can compute 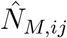 for all diversification rate shifts from any diversification rate category *j* to any other category *i*, we can build a rate shift matrix:

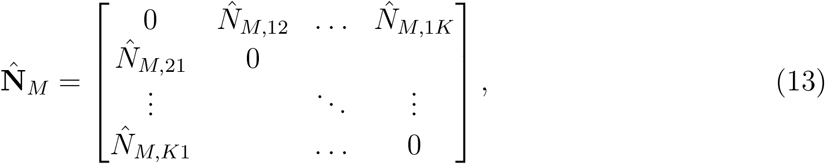

and group the elements 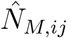 depending on which specific question we are interested in. For example, one can group 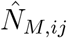 by whether it was the speciation rate or extinction rate that shifted. Another use case is to estimate the number of diversification rate shifts of a minimum shift size (e.g., *λ*_*i*_ − *λ*_*j*_ *>* 0.2). In the remainder of the manuscript, we will only focus on the total number of shifts, 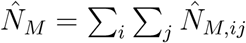, regardless of whether it was a shift in the speciation or extinction rate or both. If only the total number of diversification rate shifts is of interest, it is possible to simplify Eq. 12 to the following

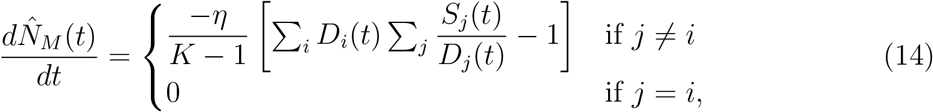

yielding the same result. We emphasize that 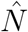 is not the probability that a diversification rate shift occurred. In a realization of the birth-death-shift process, the number of diversification rate shift events are integers, for example 0, 1 or 2. The posterior mean number of diversification rate shifts 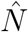 is a measure of the average or expected number of diversification rate shifts, which can take continuous values, for example 0.01 or 0.95. While this may seem counter-intuitive, it is not unlike how a six-sided dice roll has an expectation of 3.5.

### Identification of branches with strong support for a diversification rate shift

In the previous section we derived an approach to estimate the posterior mean branch-specific number of diversification rate shifts. Next, we focus on estimating whether a branch experienced at least one significant diversification rate shift. More specifically, we calculate the posterior probability of no diversification rate shifts in a small time interval Δ*t*. Over a longer time interval such as a branch, the joint probability of no diversification rate shifts can then be calculated as the product of posterior probabilities over successive small time intervals Δ*t*_1_, Δ*t*_2_, etc. For computational convenience we replace the product with a sum by using log-transformed probabilities and we express the change in the sum as a differential equation. This allows us to use standard differential equation solvers (Tsitouras, 2011), which give more precise solutions. Specifically, we calculate the posterior probability of no diversification rate shifts originating from rate category *j* on branch *M*, conditional on the process beginning in rate category *j*, represented by the quantity *X*_*M,j*_(*t*). The following differential equation describes how log *X*_*M,j*_(*t*) changes over time

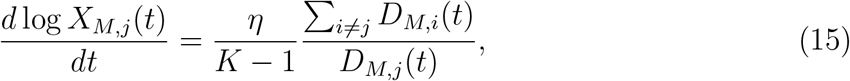

where the initial value is log *X*_*j*_(*t*_old_) = 0, and *dt <* 0. We provide details of the derivation in the supplementary material. Next, we take the weighted average across the diversification rate categories *j*, with the weights being the posterior probabilities for the ancestral rate categories (at the beginning of the branch), and take the complement. This gives us the posterior probability that there was at least one rate shift

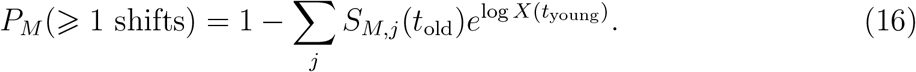

If we want a measure of support for the hypothesis of at least one diversification rate shift on a branch, we can calculate the Bayes factor (Shi and Rabosky, 2015)

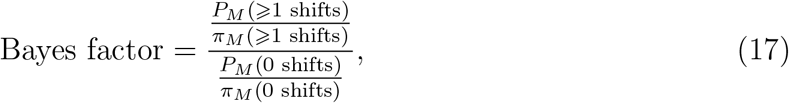

where we use 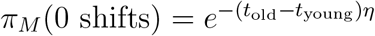 as an approximation for the prior probability of 0 diversification rate shifts —i.e., the probability of 0 events under a Poisson distribution with rate (*t*_old_ − *t*_young_)*η*.

In our exploration of how the model fitted to several test datasets, we noticed that very short branches, which have an extremely low prior probability of a diversification rate shift, can produce strongly supported Bayes factors even though the posterior mean number of diversification rates shifts is very low (e.g.,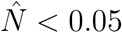). Therefore, when considering whether a branch showed strong support for a diversification rate shift event, we employed two criteria. First, we required the rate shift event to have a Bayes factor of more than 10 (Jeffreys, 1961; Kass and Raftery, 1995). Second, we required the branch to deviate substantially from an estimate of 0 rate shifts, thus we picked 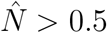 as a conservative criterion.

### Birth-death-shift simulations

To assess the performance of `Pesto`, we designed a procedure to simulate trees under the birth-death-shift process. We use a forward-simulation approach, in other words we begin at the root and move towards the tips. This approach generates “complete trees”, i.e., trees that also include extinct tips. Our simulator for complete trees rejects the tree if i) the number of lineages exceeds the maximum, or ii) if one or both subtrees descending from the root went extinct. In other words, we only accept trees where both the left and the right descendant branches of the root node have at least one surviving species (i.e., conditioning on survival of both daughter lineages of the root, see Eq. 4). Next, we prune the extinct lineages from the complete trees to form “reconstructed” trees. The information about diversification rate shifts on the surviving lineages is kept, while the records of diversification rate shifts on the extinct lineages is discarded. These final trees only have tips at the present, and appear as if the tree was reconstructed or inferred based on molecular or morphological characters of extant species.

### Simulation specifics

We simulated phylogenies under different settings to assess the performance of the `Pesto` inferences in different scenarios. Specifically, we used several models with three rate categories, ranging from low to high net-diversification rates (Table 1), with constant relative extinction (*µ/λ*). The initial diversification rate category was always the one with the lowest net-diversification rate, meaning that i) the first rate shift that could happen must be an upwards shift, and that we ii) expected upward shifts to be more common. We varied the range of rate variation (from tiny to large), meaning that the difference among the rate categories *λ*_*j*_ within each model was varying. Moreover, we simulated trees with a time interval ranging from 25 to 125 Ma, resulting in different distributions of tree size. For computational purposes, we rejected simulations with too many tips (*>* 50, 000), leading to a truncated distribution. We simulated 100 phylogenetic trees for each setting, meaning that there were in total 2000 simulated trees (5 time intervals and 4 levels of rate variation).

**Table 1:**
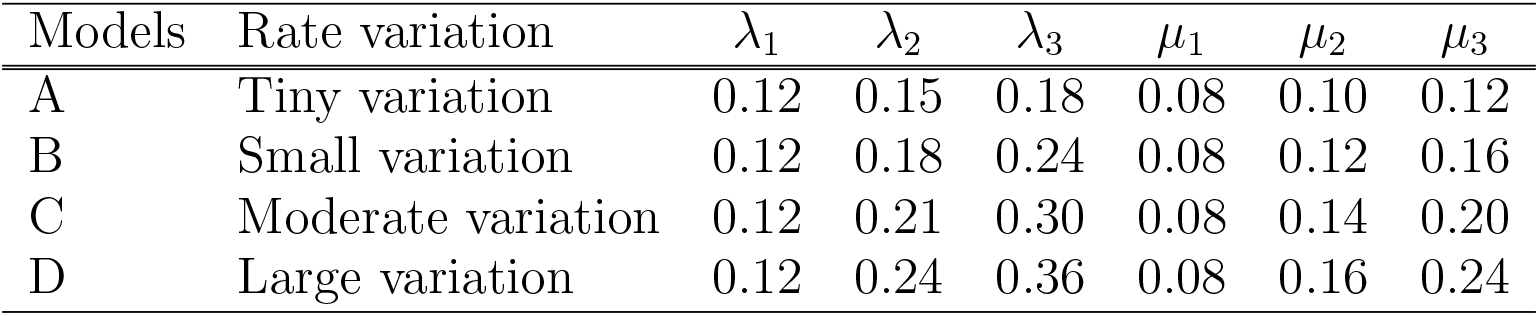
We set up a series of three-category models with net-diversification rates ***r*** = [0.04, 0.04 + *β*, 0.04 + 2*β*], and relative extinction rates of *µ/λ* = 2*/*3. *β* ∈ {0.01, 0.02, 0.03, 0.04} controls the range of allowed rate variation, from tiny to large. We simulated trees of varying height, including 25, 50, 75, 100 and 125 Ma between the most-recent common ancestor and present. All simulations begin with the first category (*λ*_1_, *µ*_1_), which results in up-shifts being common in the simulated trees. The speciation and extinction rates were chosen such that the variation in tree size was not too large (i.e. such that *n*_tips_ *>* 10^5^ was relatively rare). We selected a shift rate such that rate shift events are much less frequent than the branching events remaining in the reconstructed tree (*η* = (*λ*_1_ − *µ*_1_)*/*50 = 0.0008).

| Models | Rate variation | $\lambda_1$ | $\lambda_2$ | $\lambda_3$ | $\mu_1$ | $\mu_2$ | $\mu_3$ |
| --- | --- | --- | --- | --- | --- | --- | --- |
| A | Tiny variation | 0.12 | 0.15 | 0.18 | 0.08 | 0.10 | 0.12 |
| B | Small variation | 0.12 | 0.18 | 0.24 | 0.08 | 0.12 | 0.16 |
| C | Moderate variation | 0.12 | 0.21 | 0.30 | 0.08 | 0.14 | 0.20 |
| D | Large variation | 0.12 | 0.24 | 0.36 | 0.08 | 0.16 | 0.24 |

We tested the performance of `Pesto` under three different scenarios. First, we used the true diversification rate categories ***λ*** and ***µ*** and the true shift rate *η*. Second, we used the true diversification rate categories, but we assumed the shift rate to be unknown. Third, we assumed that both the diversification rate categories and the shift rate were were unknown. This allowed us to assess the estimates of the shift rate parameter, the inference of significant branch-specific diversification rate shifts, as well as the estimates of the branch-specific diversification rates, in various scenarios of parameter (un)certainty. To assess the inference of branch-specific diversification rate shift inferences, we used metrics that are standard in binary classification problems, including accuracy, false positive ratio, and false negative ratio (Powers, 2011).

### Implementation

We implemented our inference method in the *Julia* (Bezanson et al., 2017) module `Pesto`. `Pesto` is an acronym for *Phylogenetic Estimation of Shifts in the Tempo of Origination*. The source code and documentation is available on github in the form of vignettes with example code (*https://github.com/kopperud/Pesto.jl*). To solve the differential equations numerically, we use the Runge–Kutta algorithm of Tsitouras (2011) implemented in `DifferentialEquations.jl` (Rackauckas and Nie, 2017). We implemented the simulations in a different module, `BirthDeathSimulation.jl` (*https://github.com/kopperud/BirthDeathSimulation.jl*), where it is possible to export the tree as an extended Newick string with metadata for each node. We plotted the figures using `ggtree` (Yu et al., 2017), `Makie.jl` (Danisch and Krumbiegel, 2021) and Ti*k*Z.

## Results

We implemented an extremely efficient method for inferring branch-specific diversification rates and diversification rate shifts under the birth-death-shift process. `Pesto` uses the same underlying model, the birth-death-shift process, as the `LSBDS` implementation in `RevBayes` (Höhna et al., 2016, 2019). Note that the `LSBDS` implementation of the birth-death-shift model can estimate the parameters of the base distributions (and the shift rate) jointly using hierarchical models, whereas our approach in `Pesto` can not. Therefore, the results of a `Pesto` and `LSBDS` analysis may in practice be different. If the same model is chosen, however (i.e., the prior distributions and discretization is identical) the two methods will give exactly the same estimates. We show the equivalence in Fig. S3 and thus validated our implementation. The efficiency of our method enabled us to set up many tests for exploring various model choices and their impact on parameter estimates, which we explore in the following sections.

The primary use case of `Pesto` is to estimate branch-specific diversification rates and to infer branch-specific diversification rate shift events. As an example, we show the estimated number of accumulated diversification rate shifts and the average speciation rates —plotted with different colors and mapped on the branches of the phylogeny— for primates (Vos and Mooers, 2006, Fig. 4). Based on the primates phylogeny, our posterior mean estimate for the total number of diversification shifts is 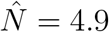, summed across all diversification rate shifts and across all branches in the phylogeny. However, only one branch experienced a significant diversification rate shift event. On the branch that led to the Old World Monkeys clade (Cercopithecidae), we estimated that the number of diversification rate shifts is far higher than what is typical on other branches (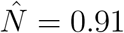 vs close to zero) and this branch had strong statistical support to have experienced at least one diversification rate shift event (with a Bayes factor of 173). This corresponds to the visual interpretation of one clear diversification rate shift (Fig. 4), whereas the additional diversification rate shifts are inferred with low frequency and low magnitude distributed over the remaining branches.

**Fig. 4:**
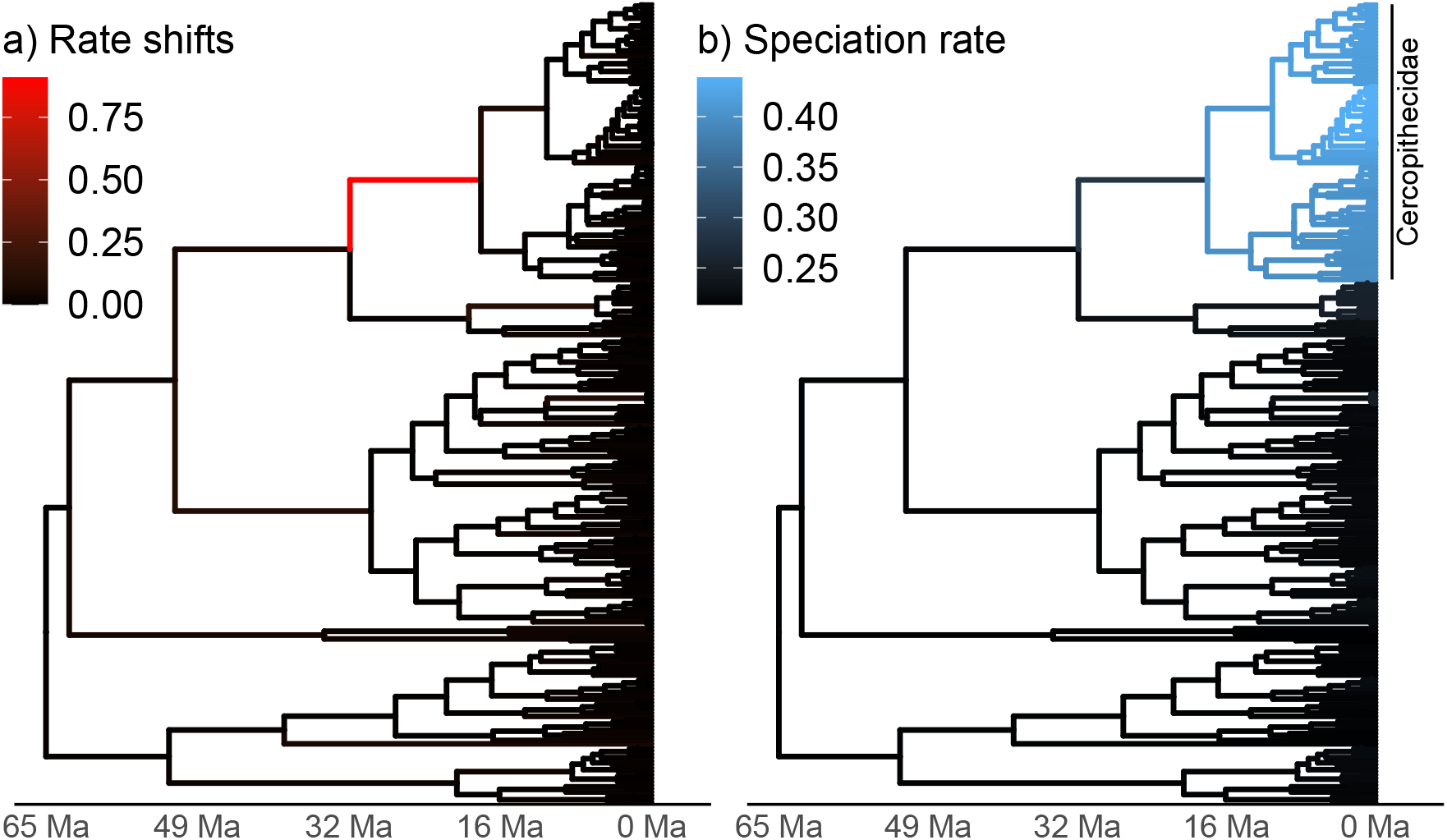
Posterior mean number of diversification rate shifts (a) and posterior mean branch-specific speciation rates (b) on the primate phylogeny (Vos and Mooers, 2006). We estimated the highest number of diversification rate shifts on the branch that led to the Old World Monkeys clade (Cercopithecidae). There are many branches with a low (but non-zero) posterior mean number of diversification rate shifts, which are not discernible using a simple color gradient. The estimate for the shift rate is *η* = 0.0032 number of diversification rate shift events per time per lineage. The posterior mean number of diversification rate shifts is 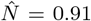 on the red branch, whereas the across the whole tree it is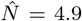. There are 233 species in the primate phylogeny, and we assumed an equal taxon sampling probability of *ρ* = 0.62 (i.e., we assumed that there were 376 species in total, Groves 2005). See Fig. 1 for the model design.

### Evaluation and Exploration of Pesto & Methods Choices

#### Impact of the diversification rate priors

When a rate shift occurs under the birth-death-shift process, the new diversification rates are determined based on a base distribution (Höhna et al., 2019). This base distribution could have many different shapes, for example, a log-normal distribution, a log-uniform distribution, an exponential distribution (e.g., as used in `BAMM`, Rabosky 2014), or a gamma distribution. So far, little is known about which base distribution is most realistic for diversification rates and what impact the choice of the base distribution has. The base distribution governs the rate heterogeneity allowed under the model. If the base distribution is too narrow or too wide, then the model may not be adequate to explain the data. Importantly, each potential base distribution has a different shape and therefore spreads the values differently.

For the primates example we see that the choice of base distribution has an impact on the estimated number of rate shifts (Fig. 5c). The posterior mean number of rate shifts were 4.9, 6.6, 5.7 and 6.6 for the log-normal, log-uniform, exponential and gamma base distributions, respectively. The pattern of one strongly supported branch was supported for the log-normal, log-uniform and gamma distributions (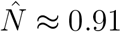 and Bayes factor in the range of 109–173). The exponential distribution also showed strong support for the Old World Monkeys branch (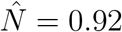 and a Bayes factor of 162), but additionally weaker support for a second branch (in the guenon clade, *Cercopithecus*). This second branch had a low number of estimated rate shifts 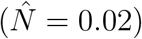 and weaker but still strong statistical support (with a Bayes factor of 11), but the overall magnitude of the rate shift was small. The pattern of many branches with few number of rate shifts 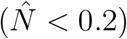 was consistent across all four choices of the base distribution. The branch-specific diversification rate estimates were quantitatively different across the four base distributions (at most by 0.05 speciation rate units), but all four choices were consistent in inferring much higher rates for the Old World Monkeys clade.

**Fig. 5:**
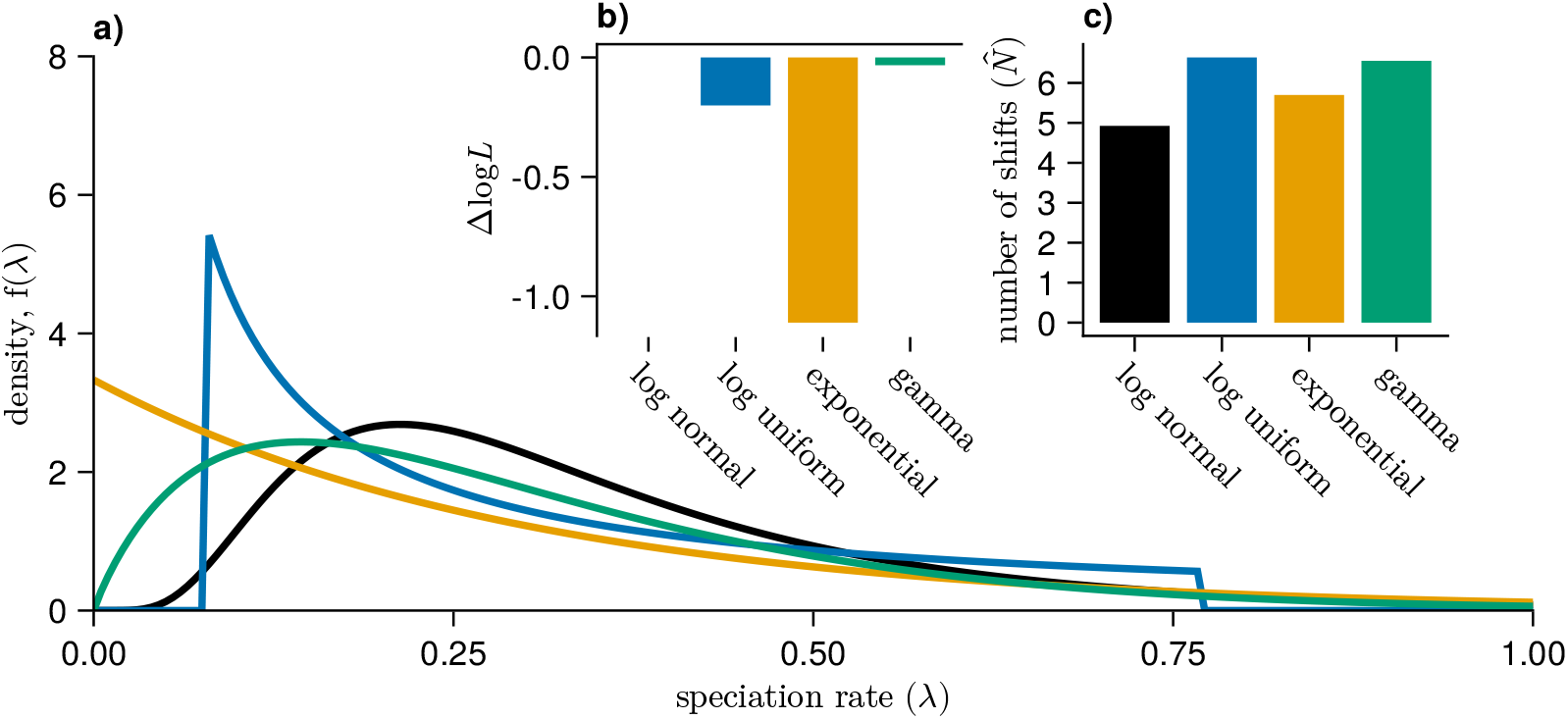
Impact of different base distributions on the branch-specific diversification rate analyses. Panel a) depicts the probability density functions of the log-normal 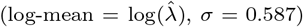, log-uniform (mean = 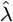, one order of magnitude range), exponential 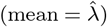 and gamma (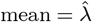, variance equal to the variance of the log-normal) distributions for the speciation rate. For all models, we drew 10 quantiles from the distributions, and we estimated *η* conditional on ***λ*** and ***µ*** using maximum likelihood. Panel b) shows the relative support (ΔlogL) for each prior distribution. Panel c) depicts the expected number of shifts across all branches in the primate phylogeny.

In principle it could be possible to choose base distributions based on model testing (Fig. 5b). For the remainder of this study we choose the log-normal distribution as the base distribution because it was the most supported distribution in this test case and has two main advantages. First, the standard deviation and thus the range of values can be specified independently of the mean (which is not possible for the exponential distribution). Second, the distribution is centered around an *a priori* specified value compared to assigning the highest density to small values, as is done for the log-uniform distribution (Fig. 5a). Overall, we expect the log-normal and gamma prior distributions to produce similar results as both distributions can resemble one another, although the gamma distribution is usually more skewed towards smaller values (Fig. 5a).

#### Impact of the number of rate categories

The base distributions of the birth-death-shift process are, in theory, full continuous distributions. Therefore, the number of possible rate categories is infinite. However, the approach implemented in `Pesto` is limited to a finite number of rate categories. Ideally, we would like to pick as many rate categories as possible, as the model will potentially fit better. In practice, we want to pick as few rate categories as possible, since more categories increase the computational time. In other words, we need to establish at what point adding more rate categories does not change the number of inferred diversification rate shifts, nor the branch-specific diversification rate estimates. We tested whether the number of allowed rate categories had a systematic impact on the estimates of the number of shifts and the branch-specific diversification rates for the primates tree (Fig. 6). When the state space is small, for example increasing the number of categories from 4 to 9, or from 9 to 16, there is some impact on the number of diversification rate shifts and the branch-specific diversification rates (Fig. 6). However, the estimates appear to stabilize when additional categories are used (*n* ⩾ 6). This indicates that the number of diversification rate shifts and branch-specific diversification rates are not systematically biased by the decision to use discrete rate categories.

**Fig. 6:**
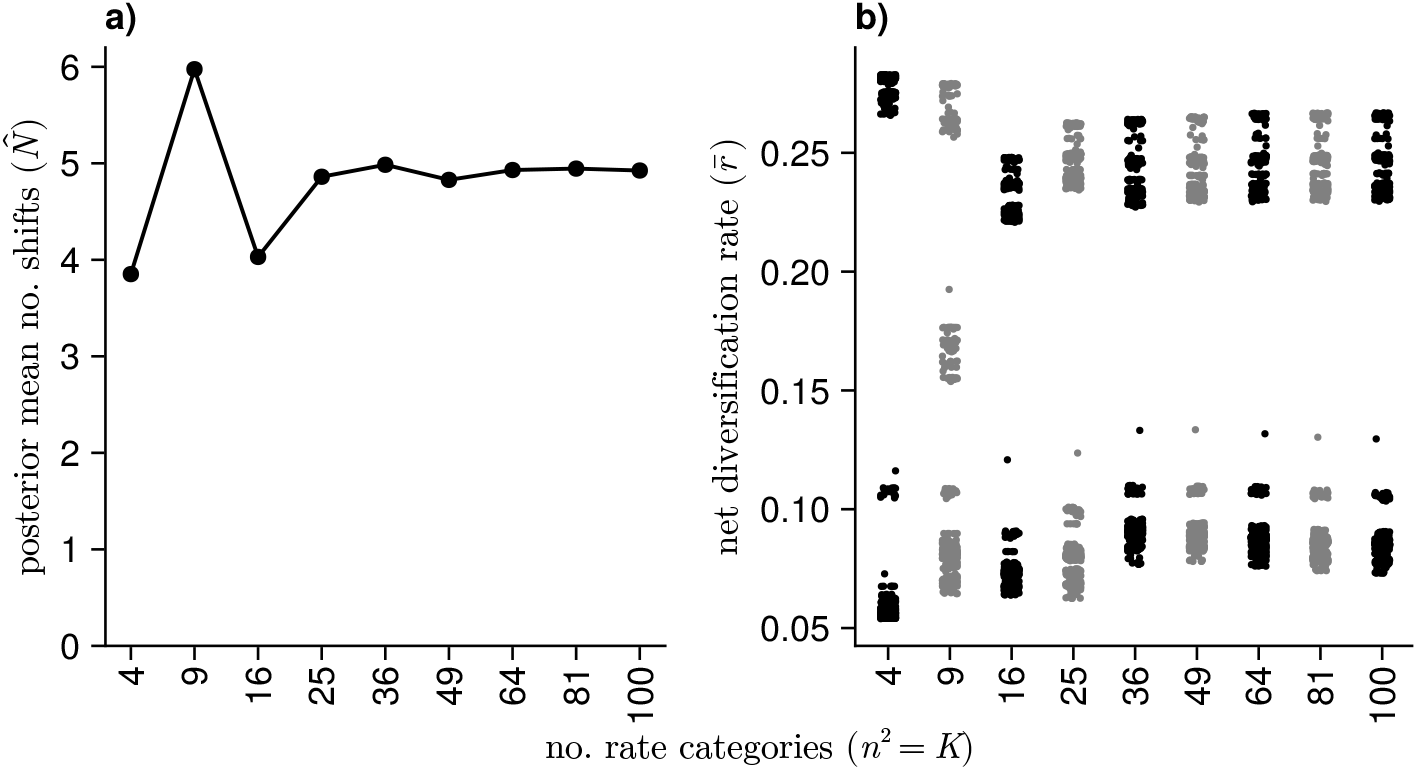
The impact of the number of rate categories on the number of rate shifts (a) and branch-specific rate estimates (b). All analyses are made on the primates phylogeny (Vos and Mooers, 2006). The bimodal distribution for average branch rates in (b) reflects the Old World Monkeys clade vs all of the other primates (see Fig. 4). The number of rate categories does not systematically bias the posterior mean number of rate shifts (a), nor the branch-specific rate estimates (b).

#### Impact of the shift rate parameter on the number of diversification rate shifts

Estimating the number of diversification rate shift events is arguably the most exciting result of branch-specific diversification rate methods (Alfaro et al., 2009; Rabosky, 2014). Previously, Moore et al. (2016) argued that estimates of the number of diversification rate shifts are not reliable and driven entirely by the choice of prior. That is, for a low rate of shifts, the methods infer few diversification rate shift events (or conversely, many events for a high shift rate). Our exploration of the impact of the shift rate parameter *η* on the posterior mean number of diversification rate shift events (Fig. 7) discovered several important aspects. First, the total number of shifts is sensitive to the shift rate *η* (see also Höhna et al., 2019). Second, the posterior mean number of diversification rate shifts is almost perfectly correlated with the prior number of shifts when the shift rate *η* is large (Fig. 7c). When *η* is small, however, the posterior number of diversification rate shifts is invariant to the rate prior (Fig. 7c). Thus, there is some but weak information in the phylogeny to estimate the number of diversification rate shifts.

**Fig. 7:**
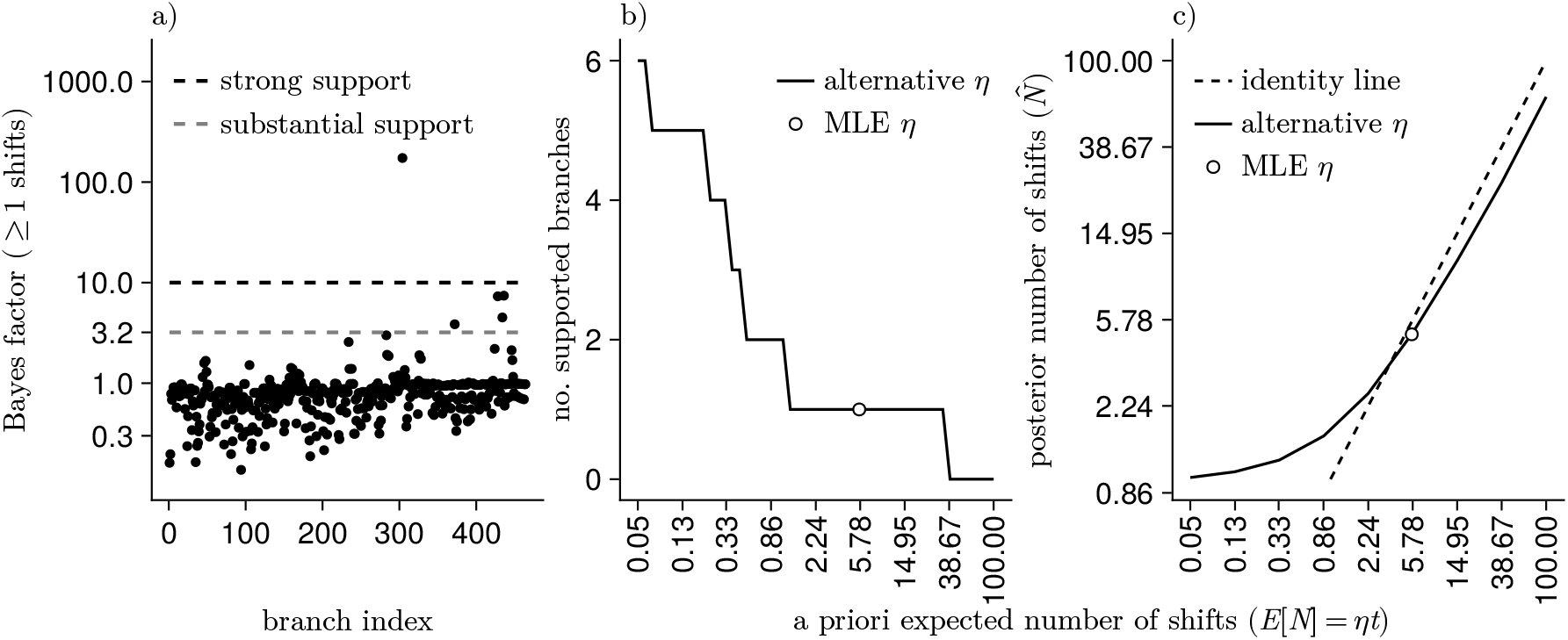
Branch-specific support for there being at least one shift, and the impact of the shift rate parameter (*η*) on the diversification rate shift inferences, for the primates phylogeny. The Bayes factors per branch (see Eq. 17) are computed using the maximum-likelihood estimate of the shift rate (*η* = 0.0032). The most supported branch, with a Bayes factor of 173, is the branch that led to the Old World Monkeys clade. If the Bayes factor is above 10, we consider the diversification rate shift to be strongly supported. Panel b) shows the impact of the shift rate (*η*) on the number of supported branches. With our maximum-likelihood estimate of the shift rate, there was one strongly supported branch. If the shift rate is much smaller, then there is more than one strongly supported branch. The posterior mean number of diversification rate shifts 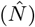 is highly correlated to the prior expectation (*E*[*N*] = *ηt*, where *t* is the tree length), except when the prior is small (c). Note that when the prior and posterior mean number of diversification rate shifts diverge (c) the Bayes factors become larger, which in turn results in statistical support for more branches (b).

A common approach to assess the support for a hypothesis, e.g., how many diversification rate shift events occurred along the phylogeny, is to use Bayes factors (Jeffreys, 1961; Kass and Raftery, 1995). Bayes factors are defined as the ratio of marginal likelihoods of the data between two hypotheses, e.g., if there was at least one diversification rate shift on the branch vs. no diversification rate shifts. The marginal likelihood is the integral over all parameter values, e.g., the branch-specific diversification rates, weighted by the prior of the parameters. Thus, Bayes factors have the advantage that they account for the prior on the number of diversification rate shift events. Nevertheless, Bayes factors are not independent of the prior on the shift rate (Fig. 7b). Since the marginal likelihood is often difficult to compute, as for the birth-death-shift process, one can compute instead the Bayes factor as the posterior ratio divided by the prior ratio (Eq. 17). We implemented and explored branch-specific Bayes factors for detecting diversification rate shifts (Fig. 7a,b). Indeed, we found that the Bayes factors are not independent of the shift rate parameter (*η*), which controls how many diversification rate shifts we expect *a priori*. Nevertheless, for the maximum likelihood estimate of the shift rate (*η* = 0.0032) we inferred one branch with a strong support for a diversification rate shift (Bayes factor = 173).

#### Run time

The run time of `Pesto` is the main advantage over similar methods that infer branch-specific diversification rates. We only perform two passes of the pruning algorithm to compute the likelihood function (backward pass) and find the posterior mean number of diversification rate shifts and branch-specific diversification rates (forward pass, Fig. 2). Furthermore, our approach has the tremendous advantage that the diversification rate category probabilities are computed precisely, without the common Monte Carlo uncertainty. Thus, convergence of branch-specific diversification rate estimates is guaranteed and convergence assessment is not necessary. Every analysis will yield exactly the same results, provided the same model and phylogeny are used.

In Fig. 8 we provide an overview of how long one can expect to run the program with various number of taxa in the phylogeny. The entire analysis takes about 1 second for a phylogeny with 100 taxa, whereas it takes less than 20 minutes for a phylogeny with 20,000 taxa. The backwards pass and the forwards pass (calculating **D**(*t*) and **F**(*t*)) scales approximately linearly with the number of tips, and quadratically with the number of rate categories. Computing the posterior mean number of diversification rate shifts 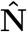 is considerably slower, and scales worse than linear (but better than quadratic) with the number of tips. Fig. 8 shows the run times using a single thread, however in the default settings we use multi-threading to calculate both the likelihood and 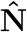 in parallel, meaning that one can expect the computation to be faster in practice. In our experience, finding the maximum-likelihood estimate for the shift rate parameter *η* is often the bottleneck of the inference procedure for small trees, requiring around 50-60 likelihood evaluations before a good estimate for *η* is found. We performed the time benchmarks using a desktop machine with an Intel Core i7-9700 CPU with a nominal clock rate of 3.0 GHz. Note that we did not want to compare the speed of `Pesto` to alternative implementations, e.g., `LSBDS`, as other implementations need to be carefully tailored to set up the length of the MCMC chain to achieve convergence. Nevertheless, `Pesto` runs easily 100 to 1000 times faster than `LSBDS`.

**Fig. 8:**
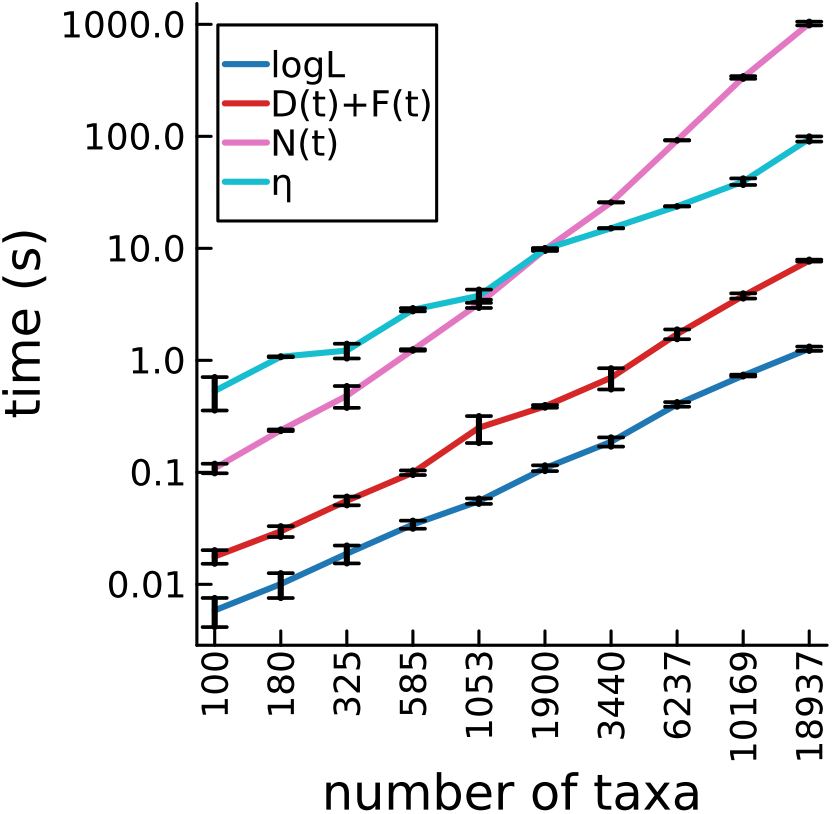
Time benchmarks of the inference procedure. The run time of *Pesto* is plotted, partitioned into the log-likelihood calculation (Eq. 4), the branch-rate calculation (postorder and preorder pass, **D**(*t*), Eq. 3 and **F**(*t*), Eq. 6), the maximum-likelihood estimation of the shift rate 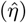, and the number of shifts calculation (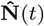, Eq. 12). We repeated these for a range of simulated phylogenies, all with *n*^2^ = 36 rate categories. All time measurements are repeated five times, and we present the median *±* the interquartile range. Time spent for installation, just-in-time compilation and garbage collection is not shown.

#### Simulation Study

In order to test the robustness of `Pesto`, we performed parameter estimation on phylogenies for which we knew the true model parameters, i.e., simulations. Note that we did not simulate under the exact same model as our inference settings, as we predefined different scenarios of rate variation (Table 1). Our prior settings for the inference on the rate variation is conservative, i.e., wider than realistic, which renders simulations computationally or practically infeasible. Furthermore, our simulations were not conditioned on specific outcomes, for example by conditioning on exactly one rate shift event, as has been done in some previous simulation studies (Rabosky, 2014). Thus, our simulations include realizations with zero, one or many rate shifts and explore the robustness of `Pesto` in a more agnostic but also more challenging scenario. Finally, the insights of our simulation study should extrapolate to other methods that rely on the same or similar underlying models and theory (see also Martínez-Gómez et al., 2024).

#### Robustness in estimates of the shift rate

We estimated the shift rate both when we considered the speciation and extinction rate categories as known and unknown. First, we will discuss the known case, where we set the speciation and extinction rate categories (***λ***, ***µ***) to their true values (see Table 1). The results indicate that it is almost impossible to accurately estimate the shift rate for small trees (*<* 100 taxa). For large trees (*>* 10, 000 taxa), the shift rate estimates converge to the true value. Fig. 9 broadly shows a bimodal distribution of estimates for the shift rate: one mode close to zero, and a second mode clearly larger than zero. Most of the trees with an estimated shift rate near zero either did not have any true shifts, or experienced shifts close to the present (e.g., on a terminal branch) during the simulation. Note that some shift rate estimates converged to our pre-defined boundary of the allowed parameter range (upper limit equal to one half times the maximum speciation rate). For these situations we noticed that the likelihood curve is flat and the hill-climbing algorithm can diverge.

**Fig. 9:**
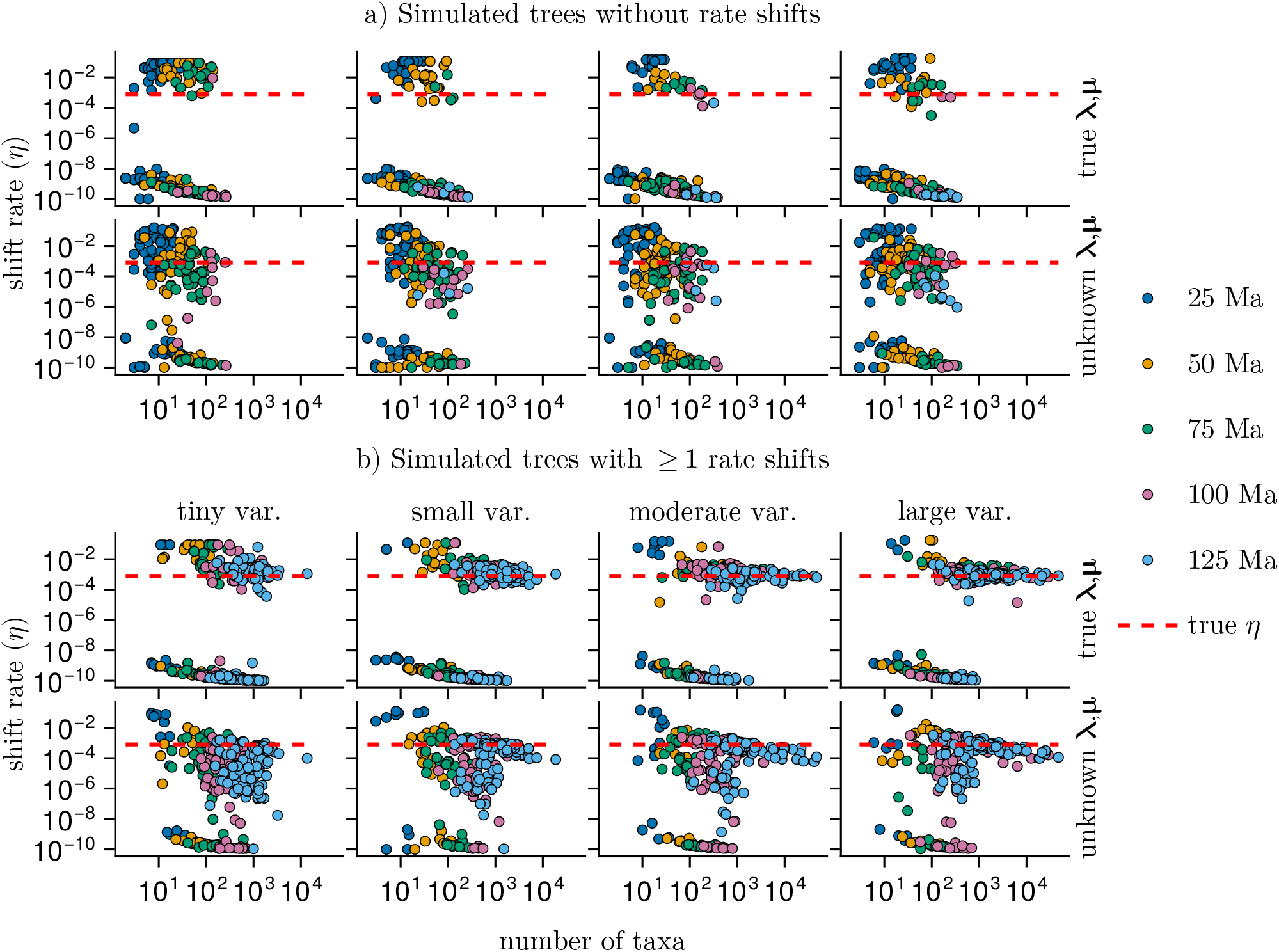
Maximum-likelihood estimates of the shift rate (*η*). Each point represents a phylogeny, simulated under the models in Table 1. The true shift rate is chosen such that diversification rate shift events are much less frequent than branching or extinction events (red line, *η* = 0.0008). The phylogenies are split by whether or not a rate shift event happened (panels a and b). We estimated the shift rate both with the true rate categories (i.e., the three-category models in Table 1) as well as using the two-step approach (i.e. first estimating 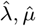 under the constant-rate birth-death process and secondly estimating the shift rate using maximum likelihood). The allowed rate variation ranges from tiny to large, and we simulated trees with a range of 25 to 125 Ma tree heights. Simulating for a longer time and increasing the rate variation has a similar effect in reducing the estimation error, since both will result in more species-rich trees.

Not surprising, when we consider the speciation and extinction rate categories (***λ, µ***) as unknown, it becomes more difficult to estimate the shift rate accurately (Fig. 9b, bottom row). Overall, the estimation error in *η* is larger if we also want to estimate the parameters of base distributions. For large trees, the shift rate estimates converge closer to the true values. However, estimates of the shift rate are biased even for very large trees (50, 000 taxa) — with underestimates of about one order of magnitude. This is likely due to the diversification rates being estimated using the constant-rate process 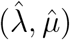, which are not representative when the true tree has undergone rate shifts.

#### Robustness of estimation in the number of diversification rate shifts

We also assessed how well `Pesto` can estimate the number of diversification rate shifts from simulated phylogenies. Altogether, 54.7% of the simulated trees experienced at least one diversification rate shift. Among these, shift histories with a single diversification rate shift were the most common (27.1%), and more numerous shifts less common. The mean number of diversification rate shifts (among the trees with at least one rate shift) was 8.3, while the median was 3. Before we discuss the quantitative results, we first present two hand-picked simulation-and-inference pairs that are useful in guiding the understanding of how the model behaves (Fig. 10). As a brief reminder, we considered a branch to have experienced a diversification rate shift if the Bayes factor was larger than 10 and the number of diversification rate shifts 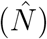 was larger than 0.5.

**Fig. 10:**
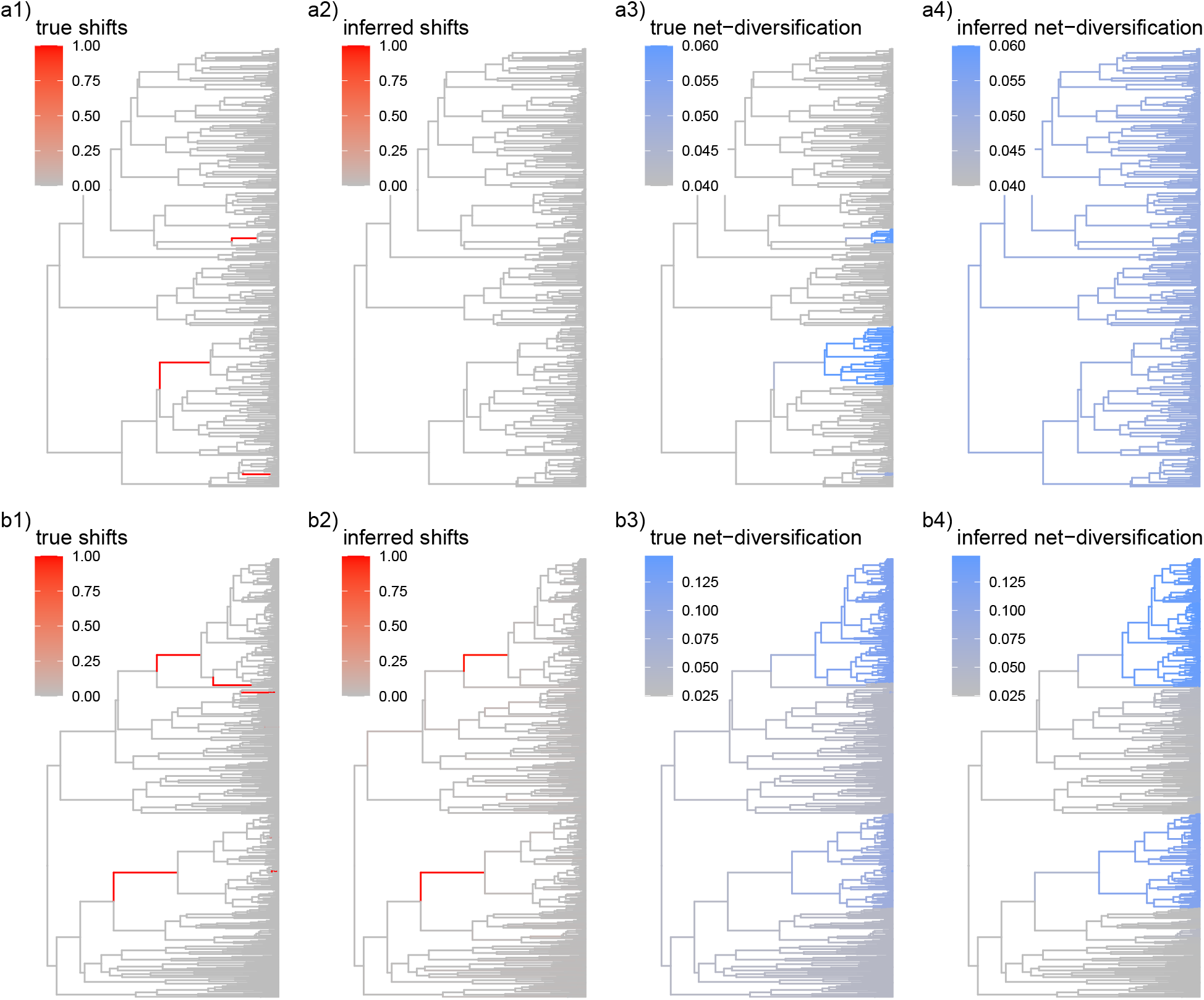
Two examples of simulated trees under the birth-death-shift model. First where *Pesto* did not recover the true shifts (a), and second where the *Pesto* analyses recovered the two oldest shifts (b). The first tree is simulated under tiny rate variation, and the second under large rate variation (models A and D in Table 1), for a period of 100 million years. We used the two-step approach in *Pesto* (with *n*^2^ = *K* = 36 categories) to estimate the rate shifts and the branch-specific diversification rates. The color scales are linked per row.

In Fig. 10, we show a tree simulated under a model that has tiny diversification rate variation, and a tree that has large diversification rate variation (models A and D, Table 1). These two trees are representative of the major trends across all 2,000 simulated trees. In the first tree, there were three diversification rate shifts, and `Pesto` incorrectly inferred zero shifts, i.e. no false positives and three false negative shifts. In the second tree, there were seven diversification rate shifts, and `Pesto` correctly inferred two major diversification rate shift events, i.e., no false positives and five false negative shifts. There are several reasons why `Pesto` might not be able to recover diversification rate shifts such as the ones in Fig. 10. As a general rule, we believe that the power to detect diversification rate shifts is high when the true diversification rate variation is large, and that the power is low when the true diversification rate variation is small. This is corroborated in Fig. 10 where we did not recover any diversification rate shifts in the tree with tiny diversification rate variation (*r*_1_ = 0.04 vs. *r*_3_ = 0.06), while we were able to detect some diversification rate shifts in the tree with large diversification rate variation (*r*_1_ = 0.04 vs. *r*_3_ = 0.12). For the diversification rate shifts that were not recovered in the second tree (despite there being a large amount of diversification rate variation), they i) occurred closer to the present, and some of the diversification rate shifts ii) led to a decrease in the net-diversification rate. Both of these scenarios make it unlikely for the descendant lineages to experience more than a few speciation events, and the resulting subclade is relatively poor in terms of the number of the species. When there are relatively few species in a particular clade, the information available for detecting a diversification rate shift is limited. In turn, this reduces the power of the method to detect diversification rate shifts.

Overall (Fig. 11), we found that the accuracy of diversification rate shift detection was very high (mean = 99.6%), the false positive ratio was tiny (mean = 0.004%), and the false negative ratio was high (mean = 94.4%). The inference of diversification rate shift events is more robust when the tree size is larger, and when the diversification rate variation is higher. We only saw substantial false positive shifts for small trees with fewer than 250 taxa. For all larger trees, the number of false positives were negligible. This means that `Pesto` is conservative by underestimating the number of diversification rate shifts. If `Pesto` infers a diversification rate shift, then there is strong evidence that this diversification rate shift corresponds to a true diversification rate shift because of the extremely low false positive ratio.

**Fig. 11:**
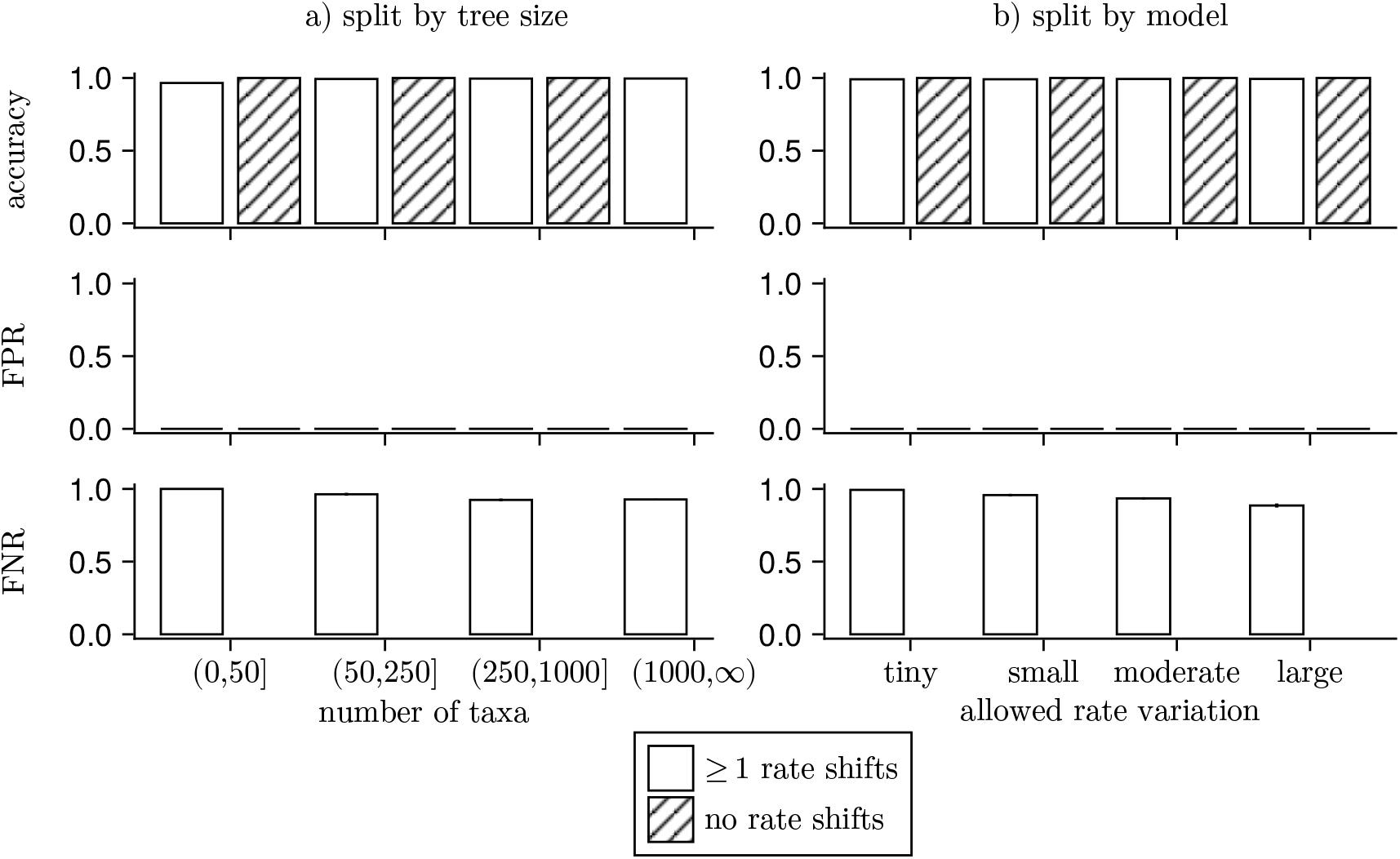
Classification of diversification rate shift inferences for 2000 simulated trees, under the two-step inference procedure, split by tree size (a), rate variation (b) and whether or not the true rate shift history contained at least one diversification rate shift. For each tree, we interpreted a diversification rate shift to be present if the Bayes factor was greater than 10 and 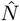 was greater than 0.5 per branch. Sorting each branch into true positives (TP), true negatives (TN), false positives (FP) and false negatives (FN), we calculated the accuracy ((TP+TN)/(TP+TN+FP+FN)), the false positive ratio (FPR = FP/(FP+TN)) and the false negative ratio (FNR = FN/(TP+FN)) for each tree. Each bar represents an average, and the error bar represents the standard error. We only calculated the false negative ratio for trees that had at least one diversification rate shift, since if TP+FN = 0, then the ratio is not defined. The false positive ratio is overwhelmingly small (mean = 0.004%), meaning that if we inferred a diversification rate shift on a branch, it was almost always present in the true tree. The false negative ratio is large, and often close to 1 (mean = 94.4%), meaning that we often inferred too few or no diversification rate shift even when there were shifts in the simulated tree. Different y-axis limits can be seen in Fig. S7.

#### Robustness of branch-specific diversification rate estimates

Next, we assessed the robustness of the branch-specific diversification rate estimates. First, we consider again the same two example trees (Fig. 10). For the tree simulated under tiny diversification rate variation, `Pesto` did not correctly infer the diversification rate shift, and thus the background net-diversification was overestimated, and the diversification rate was underestimated in the clades that shifted to a higher rate. For the second tree, `Pesto` recovered the two major shifts. Thus, `Pesto` correctly found a diversification rate difference in the two major clades that underwent a rate shift. However, the branch-specific diversification rates are slightly overestimated in the two high-rate clades, and slightly underestimated in the low-rate part of the tree.

To evaluate the estimation error in the full set of simulated trees, we calculated the proportional error in the branch-specific speciation rates (see Rabosky, 2014, eq. 15). For small trees (*<* 100 taxa), the average proportional error is roughly in the range of 0.5 to 2.0, meaning that the branch-specific diversification rates were underestimated by half, or overestimated by twice the true value (Fig. 12b, bottom row). When the true diversification rate category parameters (***λ, µ***) are used (Fig. 12, top and middle row), the proportional error converges to one when the number of taxa increases, indicating that the branch-specific diversification rate estimates are unbiased, and the precision of the method increases with larger trees. When we also estimate the diversification rate categories via the base distribution parameters, then the proportional error still converges, however with a small bias towards overestimating branch-specific diversification rates.

**Fig. 12:**
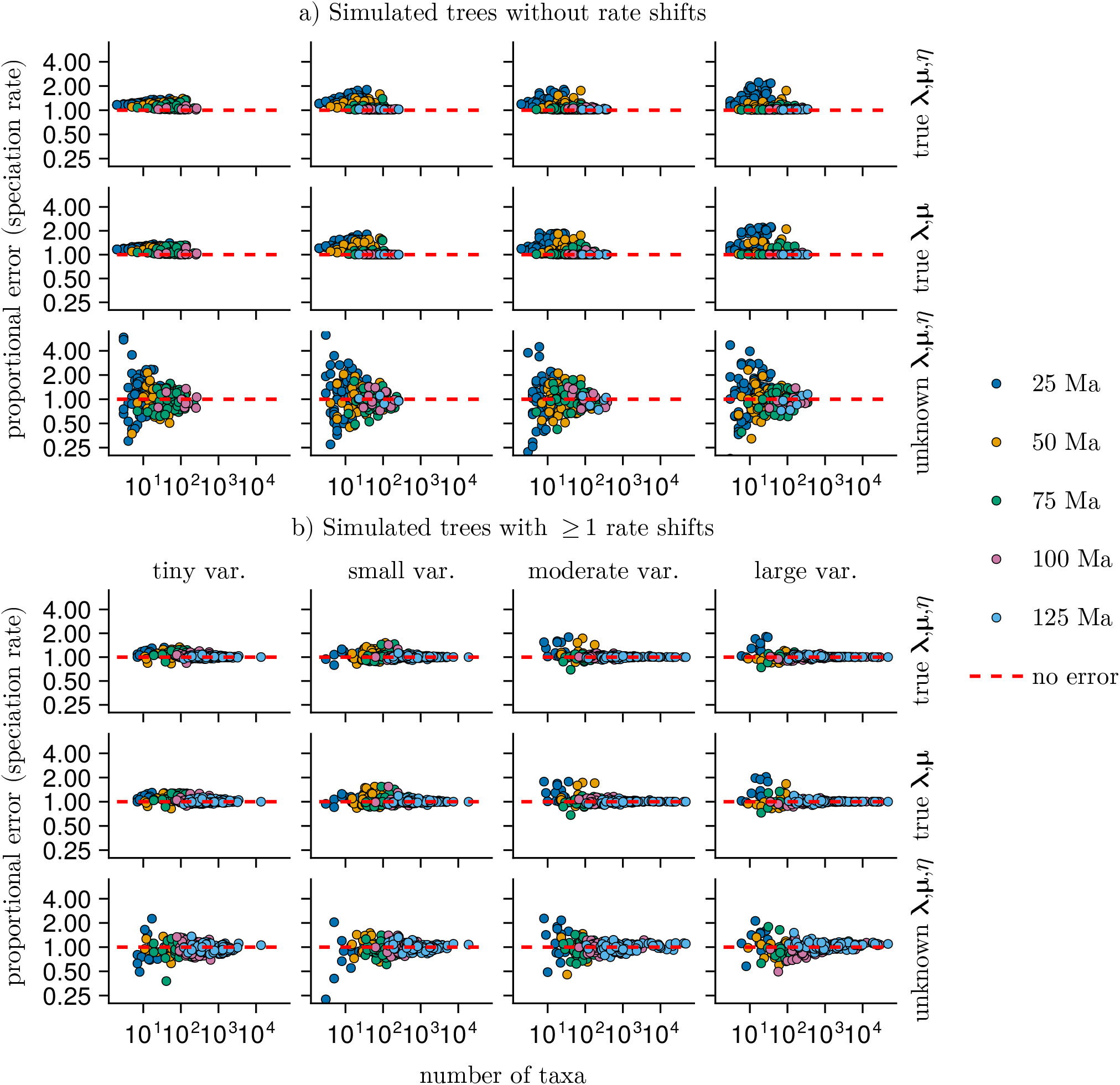
The estimation error in branch-specific speciation rates, as proportional errors, in simulated trees. Panel a) includes trees where no rate shifts happened, and panel b) includes trees where at least one rate shift happened. We calculated the proportional errors per branch, and averaged across the tree: 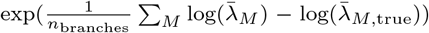, where 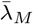 is the posterior mean speciation rate on branch *M* (Eq. 9). A value of one means that the branch-specific speciation rates are unbiased, on average across the tree. Larger than one means overestimation, and less than one means underestimation. See Figs. S7 and S8 for estimation errors in extinction and net-diversification rates.

## Discussion

In this study we developed a new approach called `Pesto` for estimating branch-specific diversification rates. The key feature of our approach is the computational speed; we can estimate branch-specific diversification rates on phylogenies with thousands of taxa without algorithmic stochasticity in minutes or hours. In this discussion, we will first discuss general aspects about our ability to infer branch-specific diversification rates and the number of diversification rate shifts that are not exclusive to `Pesto`. Then, we briefly contrast `Pesto` with other existing methods and indicate the advantages and disadvantages. Finally, we provide some general recommendation about how to use `Pesto` for inferring branch-specific diversification rates.

### Inferring diversification rate shifts

To explain how one can best interpret the branch-specific results obtained from `Pesto`, it can be useful to consider how Höhna et al. (2019) investigated the posterior distribution by simulating stochastic mappings, similar to how stochastic mappings of discrete character evolution can be simulated (Huelsenbeck et al., 2003). Our approach is conceptually the same. Consider a stochastic variable *Z* that represents the history of diversification rate shift events on the phylogeny. The posterior distribution of *Z* is determined by the model (i.e., the base distribution and its parameters, and the shift rate) and the observations (i.e., the reconstructed phylogeny). What we have done is to develop a fast algorithm that directly computes summaries of *Z*, for example the branch-specific posterior mean number of diversification rate shifts. While this approach is much faster than a stochastic mapping approach, it comes with the disadvantage that we disregard any information about the variation in *Z*, in other words how uncertain the branch-specific estimates are. If one wishes to know the extent of the uncertainty in the branch-specific estimates, we recommend to use stochastic mapping, for example as in `LSBDS` (Höhna et al., 2019).

### Sensitivity to the shift rate

Moore et al. (2016) argued based on the performance of `BAMM` that the posterior distribution of the number of diversification rate shift events is strongly related to the prior expectation, which in our case is controlled by the diversification shift rate η. Overall, we agree that there is strong sensitivity to the shift rate parameter, or in other words that the diversification rate shift events are only weakly identifiable. Despite this, our exploration of the primates phylogeny revealed that the posterior mean number of diversification rate shifts is at minimum about one, even if the shift rate is assumed to be arbitrarily small. Therefore, the hypothesis of there being no diversification rate shifts can confidently be excluded, and more generally this indicates that there is some (but weak) information in a phylogeny to detect branch-specific shifts in the diversification rate (Fig. 7c).

### Assessing significant diversification rate shifts events

Visually inspecting the branch-specific diversification rates can highlight the branches where a diversification rate shift event has occurred. Therefore, it might seem obvious to say how many branches experienced a diversification rate shift (Fig. 4). However, quantitatively assessing how many rate shifts occurred (and on which branches) has remained more challenging (Rabosky, 2014; Shi and Rabosky, 2015; May and Moore, 2016; Mitchell and Rabosky, 2017; Höhna et al., 2019). In principle, Bayes factors should be able to detect the branches for which the data and not only the prior increases the probability of a diversification rate shift. We note that the true prior distribution on the number of rate shifts is unknown, and we used a simplistic approximation for the prior. As with any statistical significance test, we recommend to assess both the hypothesis test (e.g., the Bayes factor) as well as the effect size (e.g., 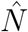, or the absolute change in net-diversification rate across the branch). In other words, if the statistical hypothesis test is significant, but the effect size is minimal, then we argue that the diversification rate shift is not biologically meaningful (Nakagawa and Cuthill, 2007). Therefore, we suggest to compute both the posterior mean number of diversification rate shifts 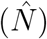 and the Bayes factor for there being at least one or more rate shift events. If the posterior mean number of diversification rate shifts on a branch 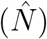 is substantially larger than 0, say 0.5, and the Bayes factor indicates strong support (say 10), then we are confident in that a diversification rate shift event actually occurred on the branch. These criteria are designed to be conservative, such that we get few false positives, and high accuracy, but not necessarily few false negatives (see also May and Moore, 2016).

In our simulation study, we observed that `Pesto` has high power to detect diversification rate shift events when there were sufficiently many descendant lineages (i.e., due to an increase in the net-diversification rate), and lower power in other cases (i.e., due to a decrease in net-diversification rate). Since roughly half of the branches in a phylogeny are terminal lineages with only one descendant, it is not surprising that `Pesto` and similar methods have low power to detect all diversification rate shift events. Finally, as expected, we find that larger diversification rate shifts are easier to detect.

### Comparison with other methods

The birth-death-shift model as implemented here and in `LSBDS` (Höhna et al., 2019) is conceptually similar to other birth-death models that allow for among-lineage rate variation, including `BAMM` (Rabosky, 2014), `ClaDS` (Maliet et al., 2019), `MTBD` (Barido-Sottani et al., 2020), `MiSSE` (Vasconcelos et al., 2022) or `RPANDA` (Mazet et al., 2023), and we provide a comparison in Table 2. For example, `ClaDS` models the diversification rate after a rate shift event as being correlated with the ancestral diversification rate, while we assume that the new diversification rate is drawn independently. Furthermore, `BAMM`, `MTBD`, `MiSSE` and our birth-death-shift model allow for diversification rates to shift along branches, unlike in `ClaDS` and `RPANDA`, where diversification rate shift events happen at branching events. `BAMM` is perhaps most similar to `Pesto` and `LSBDS`, although `BAMM` allows for the diversification rates to decay over time, while we assume the diversification rates to be constant within a diversification rate category. There are two main reasons for why `Pesto` is more computationally efficient than other methods. First, `Pesto` uses fixed values for the diversification rate categories 1). In other implementations, e.g., the `LSBDS` model in `RevBayes`, the values of the diversification rate categories can be estimated using hierarchical models. Second, unlike `LSBDS`, `BAMM`, `ClaDS` and `MTBD`, we do not make use of Monte Carlo algorithms for the inference. Instead, we use a dynamic programming algorithm to compute the marginal probabilities of the diversification rate categories across the tree, which only requires two tree traversals. This is in contrast to `MiSSE`, which uses the marginal ancestral state reconstruction algorithm of Yang et al. (1995) to obtain marginal probabilities at the nodes in the tree. The disadvantage of this marginal ancestral state reconstruction algorithm is that it requires a tree traversal for every node and every diversification rate category (Caetano et al., 2018), which is considerably more than the two in `Pesto`.

**Table 2:**
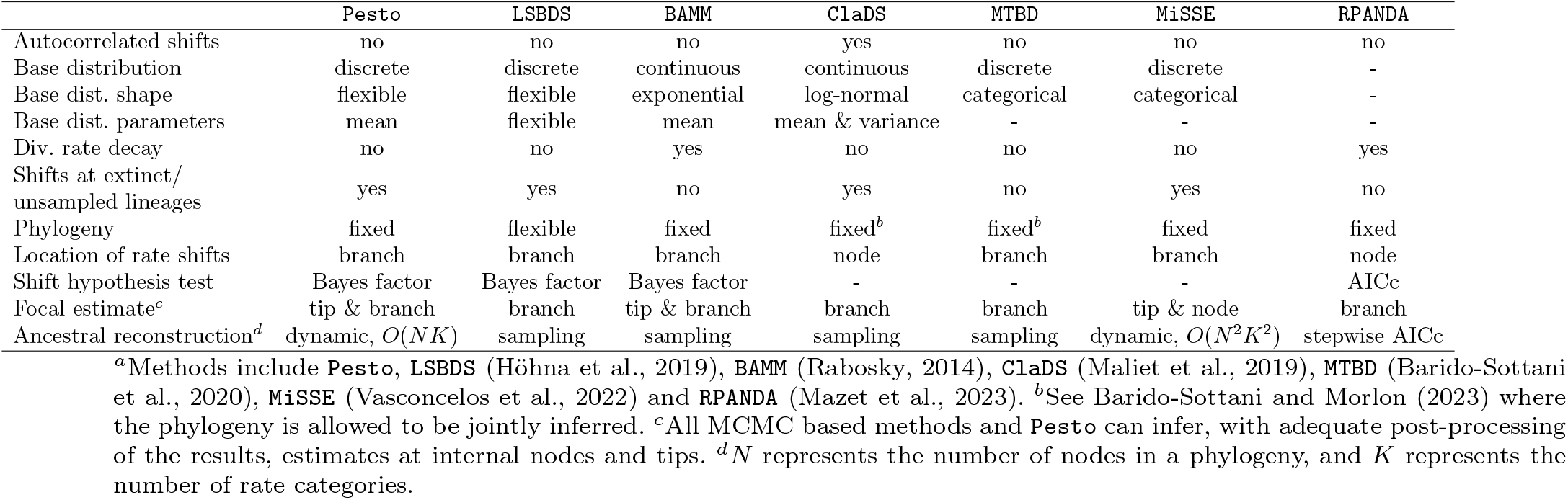
A comparison of methods^*a*^ for inferring branch-specific diversification rates and diversification rate shift events.

### Advantages and Disadvantages of Pesto

When designing how the parameters and the branch-specific metrics were to be estimated, our overall goal was for `Pesto` to be suitable to be run on exceptionally species-rich phylogenies. The main advantages of `Pesto` include i) its fast run time, ii) it is based on a generative stochastic process (unlike e.g. Chan and Moore 2005; Alfaro et al.2009), and iii) quantitative tests of the hypothesis of whether there was a diversification rate shift event. However, there are also some disadvantages with the estimation approaches that we used. Recall that we estimated the parameters of the base distributions (i.e., 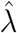 and 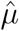) using the lineage-homogeneous, constant-rate birth-death process. While this gives a rough estimate of the overall scale of the birth and death rates, we expect that if a phylogeny has undergone diversification rate shifts, that the estimated rates obtained by fitting this simple model are biased. We suspect that the reason we see i) upward biased speciation rates and ii) downward biased shift rates, in particular for very large simulated trees, is partly due to the choice of using the lineage-homogeneous birth-death model for estimating the parameters of the base distributions. Furthermore, we investigated the posterior distribution of the rate shift histories conditionally on the parameters of the base distributions, and conditionally on the shift rate. We expect that i) joint estimation of the parameters and ii) joint inference of the parameters and the rate shift history would be more statistically robust (as in Höhna et al., 2019), however more research is needed for this to be computationally feasible for very large trees.

In keeping with most contemporaneous phylogenetic diversification rate studies (e.g. Egan et al., 2024; Quintero et al., 2024), we have assumed that the phylogeny is known without error. Even though Barido-Sottani and Morlon (2023) demonstrated that joint inference of phylogeny and a birth-death-shift model is possible, this type of joint inference is computationally challenging even for clades with few taxa, and is virtually impossible for species-rich clades. As a quicker but less robust alternative to the proper joint inference, however, one can independently apply `Pesto` to several samples from the posterior distribution of phylogenies, and interpret the variation in the results as estimation error that is due to uncertainty in the phylogeny.

### General Recommendations on how to use Pesto

Both simple and more detailed instructions on how to use `Pesto` are provided at https://kopperud.github.io/Pesto.jl/dev/. As when fitting any probabilistic model, we recommend to inspect and consider the parameter estimates and their units. The speciation rates, extinction rates, and shift rates are in units of “number of events per lineage per time”. If the phylogenetic tree is a species tree, then the time unit is typically one million years. In our experience, the shift rate should be much smaller than the mean speciation and extinction rates, by at least one or two orders of magnitude. It is biologically plausible that there are more speciation events than diversification rate shift events (note that not all speciation events are included in a reconstructed tree but all rate shift events would have to be included). If the estimated shift rate is too large, this might indicate that there was a numerical problem. For this reason, we put limits on the maximum allowed shift rate when finding the maximum likelihood value, however this hard boundary is not guaranteed to always avoid the issue.

In general, we recommend to use the log-normal distribution for the base distribution of diversification rates. Moreover, users of `Pesto` will have to decide how many rate classes to use. In principle, more rate classes are better than fewer classes as the underlying continuous distribution is better approximated. However, additional rate classes come with computational demands. For clades that are similar to the primates in terms of time span and species heterogeneity, we recommend to use at least six or more rate classes, and to use all pairwise combinations of extinction and speciation rates (i.e., at least *K* = *n*^2^ = 36 rate categories). If a phylogeny is particularly heterogeneous, we recommend to test whether six diversification rate classes is sufficient, and to use more if necessary.

This can for example be tested by fitting the model with seven or eight classes, and assessing whether the results remain similar or are substantially different.

## Conclusion

We have shown that we are able to estimate posterior mean branch-specific diversification rates and number of diversification rate shift events more efficiently than before (one computation vs. thousands or millions of simulations). The birth-death-shift model can be fitted to small trees (*<* 100 taxa) in about one second. Larger trees (*>* 10, 000 taxa) take slightly more time, on the order of a few minutes on a standard desktop computer. Compared with the established estimation technique of using Markov-chain Monte Carlo and sampling stochastic mapping of the diversification rate shift history, which can take hours, days or even weeks for large trees, our new method and implementation is several orders of magnitude more efficient. This improvement in run time is not only more convenient, but enables empiricists to broaden their taxonomic resolution. We expect that through `Pesto`, investigations of diversification rate variation across the tree of life will become more practically feasible and accessible.

## Reproducibility

The analyses and figures presented here can be reproduced by running the scripts in the accompanying github repository (*https://github.com/kopperud/pesto_ms_analyses*). The simulated trees and results are not committed to the repository due to file size restrictions, however they can be downloaded from datadryad DOI: 10.5061/dryad.mgqnk995b

## Acknowledgements

We thank Alessio Capobianco and John T. Clarke for discussions. This work was supported by the Deutsche Forschungsgemeinschaft (DFG) Emmy Noether-Program (Award HO 6201/1-1 to SH) and by the European Union (ERC, MacDrive, GA 101043187). Views and opinions expressed are however those of the authors only and do not necessarily reflect those of the European Union or the European Research Council Executive Agency. Neither the European Union nor the granting authority can be held responsible for them.

## References

Alfaro, M. E., F. Santini, C. Brock, H. Alamillo, A. Dornburg, D. L. Rabosky, G. Carnevale, and L. J. Harmon. 2009. Nine exceptional radiations plus high turnover explain species diversity in jawed vertebrates. Proceedings of the National Academy of Sciences 106:13410–13414.

Allen, A. P. and J. F. Gillooly. 2006. Assessing latitudinal gradients in speciation rates and biodiversity at the global scale. Ecology letters 9:947–954.

Barido-Sottani, J. and H. Morlon. 2023. The clads rate-heterogeneous birth–death prior for full phylogenetic inference in beast2. Systematic Biology 72:1180–1187.

Barido-Sottani, J., T. G. Vaughan, and T. Stadler. 2020. A Multitype Birth–Death Model for Bayesian Inference of Lineage-Specific Birth and Death Rates. Systematic Biology 69:973–986.

Beaulieu, J. M. and B.C. O’Meara. 2016. Detecting hidden diversification shifts in models of trait-dependent speciation and extinction. Systematic Biology 65:583–601.

Bezanson, J., A. Edelman, S. Karpinski, and V. B. Shah. 2017. Julia: A fresh approach to numerical computing. SIAM Review 59:65–98.

Budd, G. E. and R. P. Mann. 2024. A covariant model of molecular and diversification patterns through time and the history of large clades. bioRxiv Pages 2024–02.

Caetano, D. S., B. C. O’Meara, and J. M. Beaulieu. 2018. Hidden state models improve state-dependent diversification approaches, including biogeographical models. Evolution 72:2308–2324.

Chan, K. M. A. and B. R. Moore. 2005. SYMMETREE: whole-tree analysis of differential diversification rates. Bioinformatics 21:1709–1710.

Condamine, F. L., J. Rolland, S. Höhna, F. A. H. Sperling, and I. Sanmartín. 2018. Testing the role of the Red Queen and Court Jester as drivers of the macroevolution of Apollo butterflies. Systematic Biology 67:940–964.

Danisch, S. and J. Krumbiegel. 2021. Makie.jl: Flexible high-performance data visualization for Julia. Journal of Open Source Software 6:3349.

Egan, J. P., A. M. Simons, M. S. Alavi-Yeganeh, M. P. Hammer, P. Tongnunui, D. Arcila, R. Betancur-R, and D. D. Bloom. 2024. Phylogenomics, lineage diversification rates, and the evolution of diadromy in clupeiformes (anchovies, herrings, sardines, and relatives). Systematic Biology 73:683–703.

FitzJohn, R. G. 2010. Quantitative traits and diversification. Systematic Biology 59:619–633.

FitzJohn, R. G. 2012. Diversitree: comparative phylogenetic nalyses of diversification in R. Methods in Ecology and Evolution 3:1084–1092.

FitzJohn, R. G., W. P. Maddison, and S. P. Otto. 2009. Estimating trait-dependent speciation and extinction rates from incompletely resolved phylogenies. Systematic Biology 58:595–611.

Freyman, W. A. and S. Höhna. 2018. Cladogenetic and anagenetic models of chromosome number evolution: a Bayesian model averaging approach. Systematic Biology 67:1995–215.

Freyman, W. A. and S. Höhna. 2019. Stochastic character mapping of state-dependent diversification reveals the tempo of evolutionary decline in self-compatible Onagraceae lineages. Systematic Biology 68:505–519.

Goldberg, E. E. and B. Igić. 2008. On phylogenetic tests of irreversible evolution. Evolution 62:2727–2741.

Groves, C. P. 2005. Order Primates. in Mammal species of the world: a taxonomic and geographic reference (D. Wilson and D. Reeder, eds.) 3rd ed. Johns Hopkins University Press.

Hodges, S. A. and M. L. Arnold. 1995. Spurring plant diversification: are floral nectar spurs a key innovation? Proceedings of the Royal Society of London. Series B: Biological Sciences 262:343–348.

Höhna, S., W. A. Freyman, Z. Nolen, J. P. Huelsenbeck, M. R. May, and B. R. Moore. 2019. A Bayesian Approach for Estimating Branch-Specific Speciation and Extinction Rates. bioRxiv.

Höhna, S., M. J. Landis, T. A. Heath, B. Boussau, N. Lartillot, B. R. Moore, J. P. Huelsenbeck, and F. Ronquist. 2016. RevBayes: Bayesian Phylogenetic Inference Using Graphical Models and an Interactive Model-Specification Language. Systematic Biology 65:726–736.

Höhna, S., T. Stadler, F. Ronquist, and T. Britton. 2011. Inferring speciation and extinction rates under different species sampling schemes. Molecular Biology and Evolution 28:2577–2589.

Hua, X. and J. J. Wiens. 2013. How does climate influence speciation? The American Naturalist 182:1–12.

Huelsenbeck, J. P., R. Nielsen, and J. P. Bollback. 2003. Stochastic mapping of morphological characters. Systematic biology 52:131–158.

Jeffreys, H. 1961. Theory of Probability. 3rd ed. Oxford University Press.

Jetz, W., G. H. Thomas, J. B. Joy, K. Hartmann, and A. Ø. Mooers. 2012. The global diversity of birds in space and time. Nature 491:444–448.

Kass, R. E. and A. E. Raftery. 1995. Bayes factors. Journal of the american statistical association 90:773–795.

Maddison, W. P., P. E. Midford, and S. P. Otto. 2007. Estimating a binary character’s effect on speciation and extinction. Systematic Biology 56:701.

Maliet, O., F. Hartig, and H. Morlon. 2019. A model with many small shifts for estimating species-specific diversification rates. Nature Ecology & Evolution 3:1086–1092.

Martínez-Gómez, J., M. J. Song, C. M. Tribble, B. T. Kopperud, W. A. Freyman, S. Höhna, C. D. Specht, and C. J. Rothfels. 2024. Commonly used bayesian diversification methods lead to biologically meaningful differences in branch-specific rates on empirical phylogenies. Evolution Letters 8:189–199.

May, M. R. and B. R. Moore. 2016. How Well Can We Detect Lineage-Specific Diversification-Rate Shifts? A Simulation Study of Sequential AIC Methods. Systematic Biology 65:1076–1084.

May, M. R. and C. J. Rothfels. 2023. Diversification models conflate likelihood and prior, and cannot be compared using conventional model-comparison tools. Systematic Biology 72:713–722.

Mazet, N., H. Morlon, P.-h. Fabre, and F. L. Condamine. 2023. Estimating clade-specific diversification rates and palaeodiversity dynamics from reconstructed phylogenies. Methods in Ecology and Evolution 14:2575–2591.

Mitchell, J. S. and D. L. Rabosky. 2017. Bayesian model selection with BAMM: effects of the model prior on the inferred number of diversification shifts. Methods in Ecology and Evolution 8:37–46.

Moore, B. R., S. Höhna, M. R. May, B. Rannala, and J. P. Huelsenbeck. 2016. Critically evaluating the theory and performance of Bayesian analysis of macroevolutionary mixtures. Proceedings of the National Academy of Sciences 113:9569–9574.

Morlon, H. 2014. Phylogenetic approaches for studying diversification. Ecology letters 17:508–525.

Nakagawa, S. and I. C. Cuthill. 2007. Effect size, confidence interval and statistical significance: a practical guide for biologists. Biological reviews 82:591–605.

Nielsen, R. 2002. Mapping mutations on phylogenies. Systematic biology 51:729–739.

Pagel, M. 1994. Detecting correlated evolution on phylogenies: a general method for the comparative analysis of discrete characters. Proceedings of the Royal Society of London. Series B: Biological Sciences 255:37–45.

Palazzesi, L., O. Hidalgo, V. D. Barreda, F. Forest, and S. Höhna. 2022. The rise of grasslands is linked to atmospheric CO2 decline in the late Paleogene. Nature Communications 13:293.

Pawitan, Y. 2001. In all likelihood: statistical modelling and inference using likelihood. Oxford University Press.

Pearl, J. 1982. Reverend Bayes on Inference Engines: A Distributed Hierarchical Approach. Pages 133–136 in Proceedings of the National Conference on Artificial Intelligence AAAI, Pittsburgh, PA.

Pearl, J. 1988. Probabilistic Reasoning in Intelligent Systems: Networks of Plausible Inference. 1st ed. Morgan Kaufmann Publishers, San Francisco, CA.

Powers, D. M. 2011. Evaluation: From precision, recall and f-measure to roc, informedness, markedness and correlation. International Journal of Machine Learning Technology 2:37–63.

Quintero, I., N. Lartillot, and H. Morlon. 2024. Imbalanced speciation pulses sustain the radiation of mammals. Science 384:1007–1012.

Rabiner, L. R. 1989. A tutorial on hidden markov models and selected applications in speech recognition. Proceedings of the IEEE 77:257–286.

Rabosky, D. L. 2014. Automatic detection of key innovations, rate shifts, and diversity-dependence on phylogenetic trees. PLoS One 9:e89543.

Rabosky, D. L., J. Chang, P. F. Cowman, L. Sallan, M. Friedman, K. Kaschner, C. Garilao, T. J. Near, M. Coll, M. E. Alfaro, et al. 2018. An inverse latitudinal gradient in speciation rate for marine fishes. Nature 559:392–395.

Rabosky, D. L., M. Grundler, C. Anderson, P. Title, J. J. Shi, J. W. Brown, H. Huang, and J. G. Larson. 2014. BAMM tools: an R package for the analysis of evolutionary dynamics on phylogenetic trees. Methods in Ecology and Evolution 5:701–707.

Rackauckas, C. and Q. Nie. 2017. DifferentialEquations.jl – a performant and feature-rich ecosystem for solving differential equations in Julia. Journal of open research software 5.

Raup, D. M. and J. J. Sepkoski. 1982. Mass extinctions in the marine fossil record. Science 215:1501–1503.

Ricklefs, R. E. 2007. Estimating diversification rates from phylogenetic information. Trends in ecology & evolution 22:601–610.

Sanderson, M. J. and M. J. Donoghue. 1994. Shifts in Diversification Rate with the Origin of Angiosperms. Science 264:1590–1593.

Sanderson, M. J. and M. J. Donoghue. 1996. Reconstructing shifts in diversification rates on phylogenetic trees. Trends in Ecology & Evolution 11:15–20.

Sepkoski, J. J. 1998. Rates of speciation in the fossil record. Philosophical Transactions of the Royal Society of London. Series B: Biological Sciences 353:315–326.

Shi, J. J. and D. L. Rabosky. 2015. Speciation dynamics during the global radiation of extant bats. Evolution 69:1528–1545.

Silvestro, D., G. Zizka, and K. Schulte. 2014. Disentangling the effects of key innovations on the diversification of Bromelioideae (Bromeliaceae). Evolution 68:163–175.

Tsitouras, C. 2011. Runge–Kutta pairs of order 5(4) satisfying only the first column simplifying assumption. Computers & Mathematics with Applications 62:770–775.

Vasconcelos, T., B. C. O’Meara, and J. M. Beaulieu. 2022. A flexible method for estimating tip diversification rates across a range of speciation and extinction scenarios. Evolution 76:1420–1433.

Vos, R. and A. Mooers. 2006. A new dated supertree of the primates. in Inferring large phylogenies: the big tree problem (R Vos, Phd thesis). Simon Fraser University, Burnaby, British Columbia.

Weir, J. T. and D. Schluter. 2007. The latitudinal gradient in recent speciation and extinction rates of birds and mammals. Science 315:1574–1576.

Yang, Z., S. Kumar, and M. Nei. 1995. A new method of inference of ancestral nucleotide and amino acid sequences. Genetics 141:1641–1650.

Yu, G., D. K. Smith, H. Zhu, Y. Guan, and T. T.-Y. Lam. 2017. ggtree: an R package for visualization and annotation of phylogenetic trees with their covariates and other associated data. Methods in Ecology and Evolution 8:28–36.

