## Supplementary materials for "Phylogenetic Estimation of branch-specific Shifts in the Tempo of Origination"

#### Supplementary Appendix

[1, 2]Bjørn T. Kopperud [1, 2]Sebastian Höhna
[1] *GeoBio-Center LMU, Ludwig-Maximilians-Universität München, 80333 Munich, Germany*
[2] *Department of Earth and Environmental Sciences, Paleontology & Geobiology, Ludwig-Maximilians-Universität München, 80333 Munich, Germany*

### S1 Statistical inference of ancestral diversification rates

Statistical inference of ancestral diversification rates under the birth-death-shift process poses some interesting questions. We note that several previous approaches that used an underlying stochastic process used a Bayesian statistical inference framework (Rabosky, 2014; Barido-Sottani et al., 2020; Höhna et al., 2019; Martínez-Gómez et al., 2024), or at least claimed to be doing so.

The simplest possible Bayesian inference problem can be depicted like so

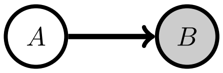

where the state of the random variable $A$ influences the state of the random variable $B$. Using Bayes theorem we can pose the question, "what is the probability of $A$, given that we know $B$?"

$$\overbrace{\Pr(A|B)}^{\text{Posterior}} = \frac{\overbrace{\Pr(B|A)}^{\text{Likelihood}} \times \overbrace{\Pr(A)}^{\text{Prior}}}{\underbrace{\Pr(B)}_{\text{Marginal Likelihood}}}. \tag{S1}$$

Here, the prior probability reflects the strength of belief in $A$ before observing $B$, and the likelihood function tells us the probability of observing $B$ given $A$. When more complex hierarchical models are considered, for example $B \leftarrow A \rightarrow C$, they are referred to more generally as *Bayesian belief networks* (Pearl, 1988). Bayesian networks that represent the splitting behaviour of a phylogeny are well established in macroevolution (e.g. Felsenstein, 1981; Pagel, 1994; Yang et al., 1995; Lewis, 2001; Höhna et al., 2014). There is also a generalized version of Bayes theorem for trees (Pearl, 1982), which we can use to ask questions such as "what is the probability of an internal node $A$ given my observations?" Despite this, we argue that Bayesian inference is not well understood in the context of birth-death-shift models.

If we are interested in Bayesian estimates of ancestral diversification rates under the birth-death-shift process, we must represent the process as a Bayesian network. Conceptually, we would like to know the probability of a hypothesis $Z$, representing a specific diversification rate history, given that we know the phylogeny $\Psi$. In the next two sections, we discuss two problems that arise when we attempt to represent the birth-death-shift process as a Bayesian network. In the first approach, we are not able to separate the diversification rate history $Z$ from the phylogeny $\Psi$. In the second approach, the "evidence" provided by the phylogeny is not encoded in the same way as for standard Bayesian networks. These two problems imply that the probabilistic representation of the birth-death-shift process is incompatible with established theory on Bayesian networks (Pearl, 1988; Russell and Norvig, 2016). Nevertheless, we provide a probabilistic method for inferring ancestral diversification rates that is inspired by Bayesian inference of ancestral discrete characters (Yang et al., 1995; Nielsen, 2002). Due to the incompatibilities, we describe our approach as being "pseudo-Bayesian", highlighting that our analysis is motivated by Bayesian inference.

#### S1.1 Problem 1: Ancestral diversification rates and phylogeny are not separable

Finding Bayesian estimators of branch-specific diversification rates on a phylogeny requires us to define what the hypothesis is and what the observation is. This is where the challenge for statistical inference of branch-specific diversification rate estimation arises. The hypotheses that we are most interested in, concerns what the ancestral diversification rates were, including the possibility that different clades were diversifying at a different pace (Höhna et al., 2019). If we simulate under a birth-death-shift process, then a realization is a specific phylogenetic tree $\Psi$, including the history $Z$ (Fig. S1). The phylogeny and the diversification rate history are linked, and it is not possible to separate the one from the other. With empirical phylogenies, however, the history of the rate shift events on the phylogeny are not recorded. This means that the process is only partially observed (hence the partially shaded node in Fig. S1). Thus, we are trying to infer the unobserved variable $Z$, and not a variable on which the data depends. This is incompatible with established theory on Bayesian inference, because there is no separation between the prior of $Z$ and the likelihood of $Z$ (see also May and Rothfels, 2023).

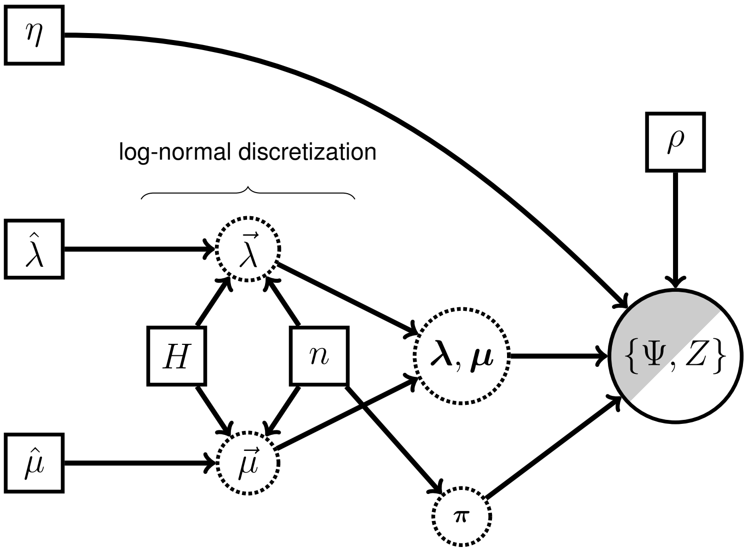

**Figure S1:** Bayesian network of the birth-death-shift model as used in `Pesto`. The network contains a node that is partially observed. This node is a stochastic variable representing the outcome of the birth-death-shift process. The node is partially shaded, meaning that we observe the reconstructed phylogeny $\Psi$, but we do not observe the diversification rate shift history $Z$. The $\hat{\lambda}$ and $\hat{\mu}$ parameters control the scale of the speciation and extinction rates, the number $H$ controls the spread of the log-normal distribution, $n$ represents the number of rate class discretizations, $\eta$ is the diversification shift rate, $\rho$ is the taxon sampling fraction, and $\boldsymbol{\pi}$ are the prior probabilities of the diversification rate categories at the root node. In `Pesto`, we estimate $\hat{\lambda}, \hat{\mu}$ using the lineage-homogeneous birth-death process, and we estimate $\eta$ using the birth-death-shift process, while we use a priori fixed values for $H, n, \rho$.

#### S1.2 Problem 2: Evidence is not encoded as evidence nodes

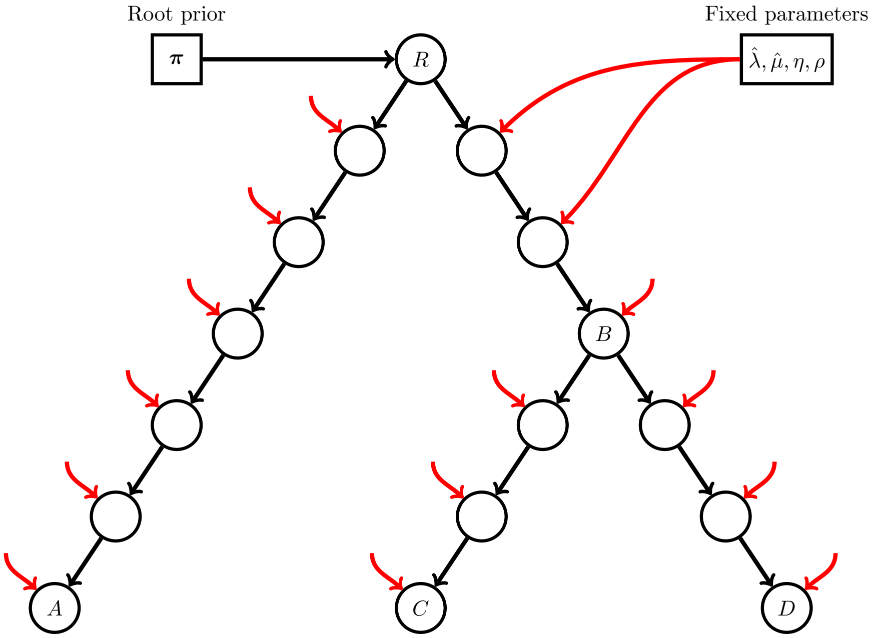

**Figure S2:** A graphical model representation of the distribution of the rate categories (circle nodes representing stochastic variables) along a three species phylogeny. The square nodes are fixed (they are not stochastic). The realization of the tree parameter $\Psi$ is represented by the branching structure of the stochastic nodes, as well as the divergence times. What we are interested in calculating, is the probability of the diversification rate category some node $B$, conditional on i) the prior probability at the root, ii) the observed phylogeny, and iii) the fixed model parameters.

Suppose we consider a more specific hypothesis, by asking "what is the diversification rate on a particular branch and at a particular time?" In order to apply Bayesian statistical inference on this problem, we need to be able to represent the relationships of the hypothesized ancestral diversification rate, with respect to the hypothesis at other branches in the tree, or at the same branch but at a different time. Specifically, we use capital letters $R, A, B, \dots$ to represent the hypothesized diversification rate category at different nodes in the phylogeny (i.e., that of the root node, and descendants of the root node). The symbol $Z$ then represents the set of all the nodes $\{R, A, B, \dots\}$. A graphical representation of the Bayesian network is shown in Fig. S2, where the diversification rate of a descendant $B$ depends on the diversification rate of its ancestor $R$.

The difference between the Bayesian network in Fig. S2 and that of a standard Bayesian network lies in how empirical evidence is encoded in the network. Typically, this is encoded as a specific evidence node (depicted as a stochastic node with gray shading, Höhna et al. 2014), where the node is initialized to for example $[1, 0, \dots]$ (as in BiSSE, Maddison et al. 2007). However, in the birth-death-shift model, the "evidence" is not encoded by initializing stochastic nodes in this way. Instead, it is encoded in the topology and branch lengths of the phylogeny. This encoding of empirical evidence is different to that of established theory on Bayesian graphical networks (Pearl, 1988). Consequently, it is not possible to pose the question "what were the ancestral diversification rate categories, given that I know the phylogeny?" as a Bayesian inference problem, in the same manner as for standard problems.

#### S1.3 Joint probability of ancestral diversification rates

As we have shown, the Bayesian network of the birth-death-shift process is incompatible with established theory on Bayesian inference. Nevertheless, we attempt to represent the hypotheses about the ancestral diversification rates $Z \in \{R, A, B, \dots\}$ in a probabilistic manner. For the root node $R$ in the phylogeny, the probability depends on the prior probability $\boldsymbol{\pi}$. It also depends on whether or not the root node is a speciation event

$$\Pr(R = r) = \begin{cases} \pi_r \Delta t \lambda_r & \text{if } R \text{ is a speciation event} \\ \pi_r (1 - \Delta t \lambda_r) & \text{if } R \text{ is a not speciation event,} \end{cases} \tag{S2}$$

where, $\Delta t \lambda_r$ is the probability of a speciation event occurring in category $r$ in a small time interval $\Delta t$, and $(1 - \Delta t \lambda_r)$ is the probability of there not being a speciation event in the interval. Since $\Delta t$ is an arbitrarily small constant, the established practice is to simplify $\Delta t \lambda_r$ to $\lambda_r$, and to omit $1 - \Delta t \lambda_r$ entirely (Maddison et al., 2007). Suppose node $R$ has a descendant node $A$, then the probability of $A$ conditionally on $R$ also depends on whether or not $A$ is a speciation event

$$\Pr(A = a | R = r) = \begin{cases} D_{a|r}(t_{\text{old}}) \lambda_a & \text{if } A \text{ is a speciation event} \\ D_{a|r}(t_{\text{old}}) & \text{if } A \text{ is a not speciation event,} \end{cases} \tag{S3}$$

where $D_{a|r}(t_{\text{old}})$ represents the $r$th entry in the solution to the $dD/dt$ differential equation, when it is initialized to $D_a(t_{\text{young}}) = 1$ and $D_{i \neq a}(t_{\text{young}}) = 0$. Note that Eq. (S3) does not sum to one over $A$. This is because not all possible outcomes are considered. For example, the process could have either gone extinct, or ended with two or more lineages (in which case node $A$ does not exist).

Eqs. (S2) and (S3) allow us to write the joint probability of the diversification rate categories $Z \in \{R, A, B, \dots\}$. If we use the shorthand notation $\Pr(A = a | R = r) = P(a, r)$, then the joint probability of the belief network in Fig. S2 is

$$\Pr(z) = \Pr(r, a, b, c, d) = \alpha \Pr(r) \Pr(a|r) \Pr(b|r) \Pr(c|b) \Pr(d|b), \tag{S4}$$

where $\alpha$ is a normalizing factor such that the joint probability sums to one. If we are interested in the probability distribution of a node, regardless of the other nodes in the phylogeny, we would analyze its marginal distribution. There are several methods one could use for calculating the marginal probabilities. The most common approach is to sample the distribution by Markov chain Monte Carlo simulation (Rabosky, 2014; Barido-Sottani et al., 2020) or stochastic mapping (Höhna et al., 2019). However, it is not necessary to use MCMC simulation methods to characterize the marginal distribution. Perhaps the simplest method is the enumeration method, in which all possible configurations of the Bayesian network are considered, and their probabilities are summed over (Russell and Norvig, 2016, p. 523). If we want to calculate the marginal probability of node $B$ using the enumeration method, we would compute

$$\Pr(B=b|\Psi) = \alpha \sum_r \sum_a \sum_c \sum_d \Pr(r)\Pr(a|r)\Pr(b|r)\Pr(c|b)\Pr(d|b), \tag{S5}$$

i.e., by marginalizing out the other nodes $R, A, C$ and $D$. If we separate and move the summation signs as far to the right as possible, as well as decompose $\Pr(b|r)$, it becomes more clear which part of the equation corresponds to the "likelihood"

$$\begin{aligned} \Pr(B=b|\Psi) &= \alpha \sum_r \Pr(r)\left(\sum_a \Pr(a|r)\right)\Pr(b|r)\left(\sum_c \Pr(c|b)\right)\left(\sum_d \Pr(d|b)\right) \\ \Pr(B=b|\Psi) &= \alpha \sum_r \Pr(r)\left(\sum_a \Pr(a|r)\right) D_{b|r}(t_R)\,\underbrace{\lambda_b \left(\sum_c \Pr(c|b)\right)\left(\sum_d \Pr(d|b)\right)}_{\text{likelihood}}, \end{aligned} \tag{S6}$$

meaning the probability of observing the tree descended from node $B$, conditional on that $B = b$. Observe that we introduce a summation sign for each node that we marginalize over. Therefore, the number of operations scales exponentially with the number of nodes. While this is not an issue for small Bayesian networks, it quickly becomes a problem for larger networks. Fortunately, more computationally efficient algorithms exist. Pearl (1982) demonstrated how marginal probabilities could be calculated efficiently using a dynamic programming algorithm (see also Gallager, 1962). By efficient we mean that the amount of operations scales linearly with the amount of nodes in the tree. While we call this algorithm the backward-forward algorithm, it is also known as a belief propagation algorithm, a message passing algorithm, or the sum-product algorithm (Teo et al., 2025). The following product rule describes the marginal probability of the diversification rate category at node $B$

$$\Pr(B=b|\Psi) = \alpha \times \Pr(\Psi^{\text{Downstream}}|B=b) \times \Pr(B=b|\Psi^{\text{Upstream}}) \tag{S7}$$

where the phylogeny $\Psi$ has been split into two parts, $\Psi^{\text{Downstream}}$ (the part of the tree that is descended from node $B$) and $\Psi^{\text{Upstream}}$ (the part of the tree that is not descended from node $B$). This product rule is fundamentally based on Bayes theorem, see Pearl (1982) for the original presentation, and (Pearl, 1988, pp. 162–174) for a more in-depth explanation. This is the same equation as in the main text, it is only the notation that is different

$$S_j(t) = \alpha D_j(t) F_j(t). \tag{S8}$$

We emphasize that Eqs. (S5) to (S8) give identical results for the marginal probabilities.

#### S1.4 Inference motivated by Bayesian thinking

Let us now take a step back to assess what type of statistical inference we used to estimate the ancestral diversification rate categories. We are, in fact, more interested in inferring these unobserved diversification rate categories than we are interested in any of the other parameters of the birth-death-shift process. The inference of ancestral (or branch-specific) diversification rate categories could be considered to be a similar problem to that of ancestral character estimation (e.g. Yang et al., 1995). The main differences are i) we assumed (perhaps unjustifiably) that we could characterize the Bayesian belief network of $Z$ conditionally on $\Psi$, and that ii) the evidence is not encoded as specific evidence nodes in the network. We specify a prior probability at the root node of the phylogeny, in accordance with our prior beliefs. The marginal probability of the diversification rate category at some node $B$ is determined by i) the prior on the root node, ii) the branching structure of the phylogeny, and iii) the fixed parameters $\boldsymbol{\lambda}, \boldsymbol{\mu}, \eta, \rho$. Since the marginal distribution depends on the prior and on the observation, we intuitively think of it as being a posterior distribution. There is one challenge, in that the Bayesian network for the birth-death-shift process is not compatible with standard theory on Bayesian inference (there is no data if the phylogeny is represented by the

Bayesian network structure). Therefore, the joint probability (Eq. (S4)) may not be a rigorous probabilistic representation of the birth-death-shift model. Hence, we suggest the terms "pseudo-Bayesian inference", and "pseudo-posterior probability," indicating that our approach is motivated by Bayesian inference. Overall, we suggest that Bayesian motivated analysis is intrinsically consistent, but more research might shed more light on this particular statistical problem.

### S2 Notation

We have tried to use notation that is consistent with the literature history, albeit with a few exceptions. Scalars are written in plain typeface, and vectors and matrices are in bold. Vectors are matrices with one column. Transposed vectors are matrices with one row.

| Symbol | Description |
|---|---|
| $K$ | The total number of rate categories in the model, indexed by $i, j \in \{1, 2, 3, \ldots K\}$. |
| $M$ | The branch index variable, i.e. $M \in \{1, 2, 3, \ldots 2(n_{\text{tips}} - 1)\}$. |
| $\odot$ | Element-wise multiplication. |
| $\oslash$ | Element-wise division. |
| $\boldsymbol{\lambda}$ | A vector with elements $\lambda_1, \lambda_2 \ldots, \lambda_j$. |
| $\boldsymbol{\mu}$ | A vector with elements $\mu_1, \mu_2 \ldots, \mu_j$. |
| $\eta$ | The shift rate from any category $j$ to any other category $i$. |
| $\frac{\eta}{K-1}$ | The shift rate from any category $j$ a different category $i$. |
| $\mathbf{Q}$ | The matrix representation of the shift rates, with $-\eta$ on the diagonal elements, and $\eta/(K-1)$ on the offdiagonal elements. |
| $\mathbf{E}(t)$ | A vector function with elements $E_j(t)$. Evaluated at age $t$ in the past, $E_j(t)$ gives the probability of the lineage not being in the sampled taxa at the present ($t = 0$). If there is complete taxon sampling, then $E_j(t)$ is the probability of the lineage going extinct before the present. |
| $\mathbf{D}(t)$ | A vector function with elements $D_j(t)$. The elements $D_j(t)$ represent the probability densities of observing the subtending clade at the present given that it is in category $j$ at age $t$. |
| $\mathbf{F}(t)$ | A vector function with elements $F_j(t)$. $F_j(t)$ is similar to $D_j(t)$, as in it is a probability density of observing the subtending clade. It is, however, given conditional on the ancestral values of $D_j(t)$. |
| $\mathbf{S}(t)$ | A vector function with elements $S_j(t)$ that represent the marginal probability of being in category $j$ at time $t$. The elements of $\mathbf{S}(t)$ sum to one. |
| $\mathbf{A}(t)$ | A matrix function which when evaluated at time $t$ yields a square matrix of size $(K, K)$. $\mathbf{A}(t)$ determines how the branch probability densities change with time. |
| $\mathbf{P}(t, t + \Delta t)$ | A square matrix of size $(K, K)$. $P_{ij}$ is the $i$th row and $j$th column of $\mathbf{P}$, and represents the probability of going from category $j$ to category $i$ in a short time interval from an older time $t$ to a younger time $t + \Delta t$, conditionally on that the process began in category $j$ at time $t$. |
| $\hat{\mathbf{N}}(t)$ | A square matrix of size $(K, K)$. $\hat{\mathbf{N}}(t)$ is the estimator for the number of rate shifts from the beginning of the branch until time $t$. Zero on the diagonal entries, and with the number of rate shifts $\hat{N}_{ij}(t)$ on the offdiagonal. |

**Table S1:** A summary and descriptions of the symbols used in the methods section.

### S3 Extended methods

#### S3.1 Number of rate shifts

The idea is to calculate how many rate shifts there were along a branch, in the time span of a small time interval. The time interval must be sufficiently short, such that the assumption that only event can happen in the time interval is reasonable. We begin by considering the rate shift probabilities. The transition probability between time $t$ and time $t + 0$ is the identity matrix, i.e. the probability of changing category is zero, and the probability of remaining in the same category is one:

$$\mathbf{P}(t, t+0) = \mathbf{I}. \tag{S9}$$

When using the matrix representation of the transition probabilities, we will follow the notation of Felsenstein (2004, page 205), in that the element $P_{ij}$ is the $i$th row and $j$th column of $\mathbf{P}$, and represents the transition probability of going from category $j$ to category $i$ in the time interval. $P_{ij}$ in a small time interval ($\Delta t < 0$, going from old to young) is approximately (Höhna et al., 2019)

$$P_{ij}(t, t+\Delta t) \approx F_{ij}(t, t+\Delta t) D_i(t+\Delta t), \tag{S10}$$

where $F_{ij}$ has been initialized as 1 in category $j$, and as 0 in all other categories $i \neq j$. Hence, $F_{ij}(t, t+\Delta t)$ is

$$F_{ij}(t, t+\Delta t) = \begin{cases} 1 - \Delta t[-(\lambda_j + \mu_j + \eta) + 2\lambda_j E_j(t)], & \text{if } j = i \\ 0 - \Delta t \frac{\eta}{K-1}, & \text{if } j \neq i \end{cases} \tag{S11}$$

Alternatively, $F_{ij}(t+\Delta t)$ can be expressed using matrix notation as $F_{ij}(t+\Delta t) = (\mathbf{e}_j - \Delta t \mathbf{A}(t)\mathbf{e}_j)_i$, where the vector $\mathbf{e}_j$ is a one-hot vector, i.e. it is 1 at index $j$, and 0 elsewhere. The matrix $\mathbf{A}(t)$ is (Louca and Pennell, 2020)

$$\mathbf{A}(t) = \text{diag}(-\boldsymbol{\lambda} - \boldsymbol{\mu} + 2\boldsymbol{\lambda} \odot \mathbf{E}(t)) + \mathbf{Q}. \tag{S12}$$

The diag(...) function constructs a diagonal matrix from a vector argument. The elements of $\mathbf{A}(t)$ are

$$\mathbf{A}(t) = \begin{bmatrix} -(\lambda_1 + \mu_1 + \eta) + 2\lambda_1 E_1(t) & \dots & \frac{\eta}{K-1} \\ \vdots & \ddots & \vdots \\ \frac{\eta}{K-1} & \dots & -(\lambda_K + \mu_K + \eta) + 2\lambda_K E_K(t) \end{bmatrix}. \tag{S13}$$

It follows that the matrix form of $P_{ij}(t, t+\Delta t)$ is

$$\mathbf{P}(t, t+\Delta t) \approx (\mathbf{I} - \Delta t \mathbf{A}(t)) \odot (\mathbf{D}(t+\Delta t)\mathbf{1}^\top), \tag{S14}$$

where $\mathbf{1}^\top$ is a row-vector of ones. For clarity, we populate $\mathbf{P}(t, t+\Delta t)$ with an example where we have $K = 3$ categories:

$$\mathbf{P}(t, t+\Delta t) = \begin{pmatrix} P_{11}(t, t+\Delta t) & P_{12}(t, t+\Delta t) & P_{13}(t, t+\Delta t) \\ P_{21}(t, t+\Delta t) & P_{22}(t, t+\Delta t) & P_{23}(t, t+\Delta t) \\ P_{31}(t, t+\Delta t) & P_{32}(t, t+\Delta t) & P_{33}(t, t+\Delta t) \end{pmatrix}, \tag{S15}$$

or approximately

$$\approx \begin{pmatrix} \begin{matrix}(1 - \Delta t[-(\lambda_1 + \mu_1 + \eta) + 2\lambda_1 E_1(t)]) \\ D_1(t+\Delta t)\end{matrix} & -\Delta t \frac{\eta}{K-1} D_1(t+\Delta t) & -\Delta t \frac{\eta}{K-1} D_1(t+\Delta t) \\ -\Delta t \frac{\eta}{K-1} D_2(t+\Delta t) & \begin{matrix}(1 - \Delta t[-(\lambda_2 + \mu_2 + \eta) + 2\lambda_2 E_2(t)]) \\ D_2(t+\Delta t)\end{matrix} & -\Delta t \frac{\eta}{K-1} D_2(t+\Delta t) \\ -\Delta t \frac{\eta}{K-1} D_3(t+\Delta t) & -\Delta t \frac{\eta}{K-1} D_3(t+\Delta t) & \begin{matrix}(1 - \Delta t[-(\lambda_3 + \mu_3 + \eta) + 2\lambda_3 E_3(t)]) \\ D_3(t+\Delta t)\end{matrix} \end{pmatrix}. \tag{S16}$$

Since the columns of Eq. (S14) don't sum to one, they do not represent a valid probability distribution. To normalize them, we divide each element by the corresponding column sum:

$$\mathbf{P}(t, t+\Delta t) = \Big[(\mathbf{I} - \Delta t \mathbf{A}(t)) \odot (\mathbf{D}(t+\Delta t)\mathbf{1}^\top)\Big] \oslash \mathbf{C}, \tag{S17}$$

where $\mathbf{C} = \mathbf{1}\mathbf{1}^\top[(\mathbf{I} - \Delta t \mathbf{A}(t)) \odot (\mathbf{D}(t+\Delta t)\mathbf{1}^\top)]$. The elements in $\mathbf{P}(t, t+\Delta t)$ are as follows:

$$P_{ij}(t, t+\Delta t) = \begin{cases} \dfrac{D_i(t+\Delta t)(1 - \Delta t[-(\lambda_i + \mu_i + \eta) + 2\lambda_i E_i(t)])}{D_j(t+\Delta t)(1 - \Delta t[-(\lambda_j + \mu_j + \eta) + 2\lambda_j E_j(t)]) - \Delta t \frac{\eta}{K-1} \sum_{i \neq j} D_i(t+\Delta t)} & \text{if } j = i \\[2ex] \dfrac{-\Delta t \frac{\eta}{K-1} D_i(t+\Delta t)}{D_j(t+\Delta t)(1 - \Delta t[-(\lambda_j + \mu_j + \eta) + 2\lambda_j E_j(t)]) - \Delta t \frac{\eta}{K-1} \sum_{i \neq j} D_i(t+\Delta t)} & \text{if } j \neq i. \end{cases} \tag{S18}$$

Apart from the normalization, we are not interested in the diagonal elements, since they do not represent rate shifts. If we weigh by the probability of beginning in ancestral category $j$ at time $t$, we can get the number of shifts in an interval from time $t$ to $t + \Delta t$:

$$\text{Shifts}_{ij}(t, t{+}\Delta t) = S_j(t) \frac{-\Delta t \frac{\eta}{K-1} D_i(t + \Delta t)}{D_j(t + \Delta t)(1 - \Delta t[-(\lambda_j + \mu_j + \eta) + 2\lambda_j E_j(t)]) - \Delta t \frac{\eta}{K-1} \sum_{i \neq j} D_i(t + \Delta t)}, \text{if } j \neq i. \tag{S19}$$

Let $\hat{N}_{ij}(t)$ be the estimator for the accumulated number of rate shifts from the beginning of the branch until time $t$. Suppose we know $\hat{N}_{ij}(t)$, or $\hat{N}_{ij}(t_0) = 0$. Then, we can calculate the number of shifts after some time $\Delta t$:

$$\hat{N}_{ij}(t{+}\Delta t) = \hat{N}_{ij}(t){+}S_j(t) \frac{-\Delta t \frac{\eta}{K-1} D_i(t + \Delta t)}{D_j(t + \Delta t)(1 - \Delta t[-(\lambda_j + \mu_j + \eta) + 2\lambda_j E_j(t)]) - \Delta t \frac{\eta}{K-1} \sum_{i \neq j} D_i(t + \Delta t)}, \text{if } j \neq i. \tag{S20}$$

Subtract $\hat{N}_{ij}(t)$ and divide by $\Delta t$:

$$\frac{\hat{N}_{ij}(t + \Delta t) - \hat{N}_{ij}(t)}{\Delta t} = \frac{-\eta}{K-1} S_j(t) \frac{D_i(t + \Delta t)}{D_j(t)(1 - \Delta t[-(\lambda_j + \mu_j + \eta) + 2\lambda_j E_j(t)]) - \Delta t \frac{\eta}{K-1} \sum_{i \neq j} D_i(t + \Delta t)}, \text{if } j \neq i. \tag{S21}$$

If we take the limit as $\Delta t \to 0$, then the expression simplifies (and we set the derivative to 0 if $j = i$ since we don't need these):

$$\lim_{\Delta t \to 0} \frac{\hat{N}_{ij}(t + \Delta t) - \hat{N}_{ij}(t)}{\Delta t} = \frac{d\hat{N}_{ij}(t)}{dt} = \begin{cases} -S_j(t) \dfrac{D_i(t)}{D_j(t)} \dfrac{\eta}{K-1} & \text{if } j \neq i \\ 0 & \text{if } j = i. \end{cases} \tag{S22}$$

In matrix form, this is

$$\frac{d\hat{\mathbf{N}}(t)}{dt} = \frac{-\eta}{K-1} \odot (\mathbf{1}\mathbf{1}^\top - \mathbf{I}) \odot \mathbf{D}(t)\mathbf{S}(t)^\top \oslash \mathbf{1}\mathbf{D}(t)^\top. \tag{S23}$$

We can solve Eq. (S23) using the same numerical integration techniques as before, over the time interval of the branch, with initial condition $\hat{\mathbf{N}}(t_0) = \mathbf{0}$. The number of shifts among all pairs $(i, j)$ for all $j \neq i$, in the time direction of old to young, are on the off-diagonals. This represents $K(K-1)$ non-zero differential equations, and so if the state space $K$ is large, then computing the number of rate shifts $\hat{\mathbf{N}}$ can be slow. If we sum over all entries in $\hat{\mathbf{N}}$, we obtain an estimate of the total number of shifts in the time interval from $t_0$ until $t$:

$$\hat{N}(t) = \mathbf{1}^\top \hat{\mathbf{N}}(t) \mathbf{1}, \tag{S24}$$

and we can compare with the results from simulated mappings of rate category histories, as implemented in RevBayes (Fig. S3). The posterior number of rate shifts $\hat{N}$ can also be summed across all of the branches, in order to get an overall idea of how many shifts there were in the entire phylogeny.

#### S3.2 Probability of a shift

We want to calculate the probability that there was at least one shift on a branch:

$$P_{\geq 1 \text{ shifts}} = \ldots, \tag{S25}$$

or equivalently, the probability that there were zero shifts

$$P_{0 \text{ shifts}} = 1 - P_{\geq 1 \text{ shifts}}. \tag{S26}$$

One way to do so would be to begin by calculating the probability that there were no rate shifts in rate category $j$. Remember that the probabilities in Eq. (S18) are conditional on that the current category is $j$, i.e. they sum to one over the arrival categories $i$. Conditional on that the process began in the departure category $j$, we can calculate the probability that at least no shifts with departure category $j$ occurred as follows

$$
\begin{aligned}
P_{0\text{ shifts},j}(t, t+\Delta t) &= P_{jj}(t, t+\Delta t) \\
P_{0\text{ shifts},j}(t, t+\Delta t) &= \frac{D_i(t+\Delta t)(1-\Delta t[-(\lambda_i+\mu_i+\eta)+2\lambda_i E_i(t)])}{D_j(t+\Delta t)(1-\Delta t[-(\lambda_j+\mu_j+\eta)+2\lambda_j E_j(t)]) - \Delta t \frac{\eta}{K-1}\sum_{i\neq j} D_i(t+\Delta t)}.
\end{aligned} \tag{S27}
$$

Next we want to calculate the probability that there were no rate shifts away from category $j$ along the entire branch, meaning we need to take the product of several probabilities. We can write it recursively as follows, by introducing a dummy variable $X_j(t)$ that represents the probability that there were no rate shifts departing from category $j$ until time $t$. We know the probability $X_j(t)$ at the beginning of a time span, it is simply set to one. Then, the probability of no shifts after some small time interval $\Delta t$ has elapsed is simply the previous probability times the transition probability

$$
X_j(t+\Delta t) = Xj(t)P_{jj}(t, t+\Delta t), \tag{S28}
$$

representing the progression along the branch. Because sums are typically more numerically stable than products, we will instead use the log transformed probability

$$
\begin{aligned}
\log X_j(t+\Delta t) &= \log X_j(t) + \log P_{jj}(t, t+\Delta t) \\
\log X_j(t+\Delta t) - \log X_j(t) &= \log P_{jj}(t, t+\Delta t),
\end{aligned} \tag{S29}
$$

and next we populate $\log P_{jj}(t, t+\Delta t)$ using Eq. (S18). Because there will be a lot of symbols, we use a substitute $c_j$ representing $-(\lambda_j+\mu_j+\eta)+2\lambda_j E_j(t)$ and shorten the equations a bit. Then, we have

$$
\log X_j(t+\Delta t) - \log X_j(t) = \log\left[\frac{D_j(t+\Delta t)(1-\Delta t c_j)}{D_j(t+\Delta t)(1-\Delta t c_j) - \Delta t\frac{\eta}{K-1}\sum_{i\neq j} D_i(t+\Delta t)}\right] \tag{S30}
$$

and we can divide by $\Delta t$ on both sides and take the limit

$$
\begin{aligned}
\frac{\log X_j(t+\Delta t) - \log X_j(t)}{\Delta t} &= \frac{1}{\Delta t}\log\left[\frac{D_j(t+\Delta t)(1-\Delta t c_j)}{D_j(t+\Delta t)(1-\Delta t c_j) - \Delta t\frac{\eta}{K-1}\sum_{i\neq j} D_i(t+\Delta t)}\right] \\
\lim_{\Delta t\to 0}\frac{\log X_j(t+\Delta t) - \log X_j(t)}{\Delta t} &= \lim_{\Delta t\to 0}\frac{1}{\Delta t}\log\left[\frac{D_j(t+\Delta t)(1-\Delta t c_j)}{D_j(t+\Delta t)(1-\Delta t c_j) - \Delta t\frac{\eta}{K-1}\sum_{i\neq j} D_i(t+\Delta t)}\right].
\end{aligned} \tag{S31}
$$

Note that the limits of $1/\Delta t$ and the log term both converge to zero. In this case we can use L'Hôpital's rule, differentiating the numerator and the denominator with respect to $\Delta t$. The left term vanishes, and it can be shown that the derivative for the log-term is

$$
\frac{d}{d\Delta t}\log\left[\frac{D_j(t+\Delta t)(1-\Delta t c_j)}{D_j(t+\Delta t)(1-\Delta t c_j) - \Delta t\frac{\eta}{K-1}\sum_{i\neq j} D_i(t+\Delta t)}\right] = \frac{\frac{\eta}{K-1}\sum_{i\neq j} D_i(t+\Delta t)}{D_j(t+\Delta t) + \alpha\Delta t + \beta\Delta t^2}, \tag{S32}
$$

where $\alpha$ and $\beta$ is represent some terms that are tedious to write. When evaluating the limit of $\Delta t \to 0$, then $\alpha$ and $\beta$ vanish, and we get the following differential equation

$$
\frac{d\log X_j(t)}{dt} = \frac{\eta}{K-1}\frac{\sum_{i\neq j} D_i(t)}{D_j(t)}, \tag{S33}
$$

which we can solve for numerically using the initial condition $\log X_j(t_{\text{old}}) = 1$. Note that $dt$ is a negative number, since we are integrating from the past to the present. Recall that all this is conditional on that the process began in rate category $j$. In order to get the posterior probability that there were no rate shifts in any rate category, we need to take the weighted average as

$$
P_{0\text{ shifts}} = \sum_j S_j(t_{\text{old}})e^{\log X_j(t_{\text{young}})}, \tag{S34}
$$

where $S_j(t_{\text{old}})$ is the posterior probability that the process was in rate category $j$ at the beginning of the branch, and $X_j(t_{\text{young}})$ is the probability that there were no rate shifts departing from rate category $j$ in the time interval $\{t_{\text{old}}, t_{\text{young}}\}$ (i.e. the branch time). It follows that the posterior probability that there was at least one rate shift is

$$P_{\geq 1 \text{ shifts}} = 1 - P_{0 \text{ shifts}}. \tag{S35}$$

The probability obtained by this equation is validated by comparing with an alternative simulation approach in Fig. S3c.

#### Numerical solutions

We use the algorithm by Tsitouras (2011), a fourth-order Runge-Kutta method, implemented in the Julia module `DifferentialEquations` (Rackauckas and Nie, 2017) in order to numerically solve the systems of differential equations. Conditional on some tolerance values (we used abstol = $10^{-3}$, reltol = $10^{-6}$), the algorithm adaptively selects a set of time points $t_i$ to solve the system of differential equations. It will fit the trajectory of the probability densities at times $t_i \in [t_0, t_{\text{end}}]$, where $t_0$ is the beginning, and $t_{\text{end}}$ is the end. We store the exact solutions at the time points $t_i$. To get values at intermediate time points (between $t_i$ and $t_{i+1}$), we fit and evaluate an interpolating function (a fourth-order polynomial) to get an approximate result. First, we solve $d\mathbf{E}/dt$ across the time span of the phylogeny, without regard to the phylogeny. In a second step, we iterate in a postorder traversal of the tree, and solve $d\mathbf{D}/dt$ for each branch, storing the resulting $\mathbf{D}(t)$, evaluating the polynomial representation of $\mathbf{E}(t)$ where necessary. In a third step, we iterate over the tree in a preorder traversal. We solve $d\mathbf{F}/dt$ for each branch, using the polynomial representations of $\mathbf{E}(t)$ and $\mathbf{D}(t)$ where necessary, and store the result $\mathbf{F}(t)$. At the end of each branch, we normalize by a scaling factor:

$$\begin{aligned}\mathbf{F}_M(t_{\text{youngest}}) &:= \frac{\mathbf{F}_M(t_{\text{youngest}})}{\theta_{M,F}},\\ \mathbf{D}_M(t_{\text{oldest}}) &:= \frac{\mathbf{D}_M(t_{\text{oldest}})}{\theta_{M,D}}.\end{aligned} \tag{S36}$$

We follow the implementation of `diversitree` (FitzJohn, 2012) and use a scaling factor such that the normalized probabilities sum to unity: $\theta_{M,D} = \mathbf{1}^\top \mathbf{D}_M(t_{\text{oldest}})$. Since the proportionality of the probabilities are preserved, we can use the normalized values in the initial conditions for the next step in the tree iteration. This avoids numerical problems where the products of $\mathbf{D}$ and $\mathbf{F}$ across many branches become too tiny, and floating point operations become inaccurate. When we compute the log-likelihood, we add the scaling factors back in ($\theta_{M,D}$).

### S4 RevBayes equivalence

The birth-death-shift model implemented in `Pesto` is fundamentally the same model as the one used in `RevBayes` (Höhna et al., 2019), with a few technical caveats. In `Pesto`, the branch-specific rate estimates and the shift inferences are calculated conditionally on the diversification rate categories (i.e. $\boldsymbol{\lambda}, \boldsymbol{\mu}$) and the shift rate ($\eta$). If the diversification rate categories and the shift rate are estimated jointly using a hierarchical model, as in `RevBayes`, then the two approaches are not identical. If all rate categories and parameters are kept equal, however, then the `RevBayes` and `Pesto` implementations are theoretically equivalent, and should give the same result. In Fig. S3, we calculated a) the posterior mean branch-specific speciation rates, b) the posterior mean number of rate shift events, and c) the posterior probability that there was at least one rate shift — all per branch, using both `Pesto` and `RevBayes`. We used the same model in both scenarios, with the rate categories being fixed to $\boldsymbol{\lambda} = [0.1, 0.2, 0.4, 0.1]$, $\boldsymbol{\mu} = [0.05, 0.15, 0.10, 0.2]$, and the shift rate fixed to $\eta = 0.00248$. Given sufficiently many stochastic character maps (we used 25000), the branch-specific estimates should correspond exactly. In a qualitative sense, our comparison shows that the branch estimates are corresponding. There are two parts of the tree with different rates, the Old World Monkeys clade vs the rest. There is also one branch where there was strong support for a rate shift ($N > 0.8$).

In a quantitative sense, however, there are some differences between the `RevBayes` and the `Pesto` analysis results, as there are a few deviations from the one-to-one line. We attribute this to numerical differences in solving the ordinary differential equations, and the stochastic character maps, which are both approximate methods. Both `Pesto` and `RevBayes` use a Runge-Kutta algorithm with adaptive step size. `Pesto` uses the `Tsit5` method (Tsitouras, 2011) implemented in the `DifferentialEquations.jl` Julia library (Rackauckas and Nie, 2017), while `RevBayes` uses the Dormand–Prince method (Dormand and Prince, 1980) implemented in the `boost` C++ library. Both of these algorithms are fundamentally similar, and give similar solutions. `RevBayes` uses a fixed step size on top of this, to simulate stochastic character maps forward in time (with standard settings `nTimeSlices = 500`, i.e. on average 500 time slices on a lineage spanning from the root to the tip of the phylogeny), and saves the result only on these specific time points. `Pesto` also saves the intermediate solutions (between the beginning and the end of the branch), but not nearly as many, and the number of saved states are determined dynamically instead of a priori. If the probability densities are relatively flat along a branch, then `Pesto` saves only a handful of time points, e.g. 3 or 4. If the probability densities vary drastically along a branch, i.e. the derivative is different to zero, then `Pesto` will save several more, e.g. 20-40. In practice, this means that `RevBayes` uses far more steps (i.e. evaluations of the difference equation) than `Pesto`, and so it is slower. `Pesto` also saves the solution for every branch, instead of discarding it. There is a trade-off in that `Pesto` saves more information, and the drawback is that `Pesto` requires more memory. Furthermore, `RevBayes` has to repeat the stochastic character maps thousands of times before the branch rate estimates converge, whereas `Pesto` only needs to calculate the branch rates once.

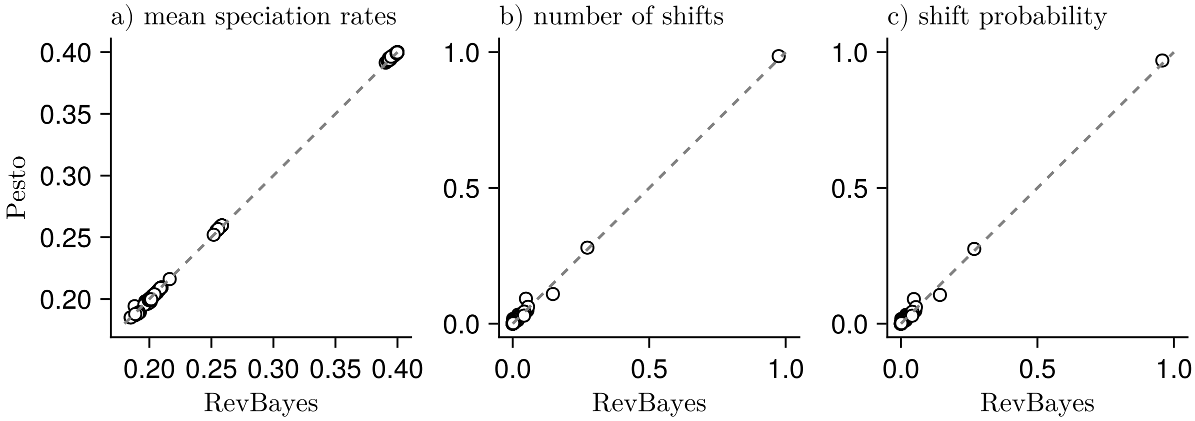

**Figure S3:** Comparisons of branch-specific calculations between `Pesto` and `RevBayes`. We calculated a) the posterior mean speciation rates, b) the posterior mean number of rate shift events, and c) the posterior probability that there was at least one rate shift. Each point represents one branch in the primates tree. The rate categories were set to $\boldsymbol{\lambda} = [0.1, 0.2, 0.4, 0.1]$, $\boldsymbol{\mu} = [0.05, 0.15, 0.10, 0.2]$, and the shift was was set to $\eta = 0.00248$ (i.e. the maximum-likelihood shift rate, conditional on $\boldsymbol{\lambda}, \boldsymbol{\mu}$. We conditioned on survival in both cases. In `RevBayes`, we used the setting `nTimeSlices = 500`, and drew 25000 independent stochastic character maps. The value on the x-axis is the mean across the 25000 stochastic character maps. The dashed line represents the one-to-one line.

### S5 Variance of branch-specific speciation rates

Whereas in the previous section we plotted branch-specific speciation rates that were averaged across the branch length, it is also possible to compute branch-specific diversification rates that are evaluated at a specific point $t$ in time. Since the `RevBayes` implementation does not by default print speciation rates that are specific to a particular time, we re-implemented the stochastic mapping in a `Julia` script. Suppose we are given an initial probability vector $\mathbf{S}(t)$, where $t$ is the oldest time of the branch, and we use some small step size $\Delta t$. Note that we are going from the past to the present, and therefore $\Delta t$ is a negative number. The simulation algorithm can be described as follows

1. Draw a rate category $j$ according to the probability vector $\mathbf{S}(t)$.

2. Initialize $F_j(t) = 1$ and set $F(t)$ to 0 for all other categories.

3. Find a solution to $\mathbf{F}(t + \Delta t)$ using numerical integration.

4. Update the probability vector $\mathbf{S}(t+\Delta t) := \alpha \mathbf{F}(t+\Delta t) \odot \mathbf{D}(t+\Delta t)$, where $\alpha$ is a normalizing constant such that $\mathbf{S}(t+\Delta t)$ sums to one.

5. Set $t := t + \Delta t$.

6. Repeat steps 1–5 until the youngest time point of the branch is reached.

This will give us a vector of the diversification rate categories, for example $[1, 1, 3, 3, 3, \ldots]$, which we can use to calculate the mean and variance of the speciation rate for each time bin. Suppose that $\lambda(t)$ is a discrete random variable with possible values $\lambda_1, \lambda_2, \ldots$ and probabilities $S_1(t), S_2(t), \ldots$, then we can calculate the theoretical expectation and variance for a particular time $t$

$$\begin{aligned}
\mathbb{E}[\lambda(t)] &= \sum_j \lambda_i S_j(t) \\
\mathrm{Var}[\lambda(t)] &= \mathbb{E}[\lambda(t)^2] - \mathbb{E}[\lambda(t)]^2 \\
\mathrm{Var}[\lambda(t)] &= \sum_j \lambda_i^2 S_j(t) - \left( \sum_j \lambda_j S_j(t) \right)^2 .
\end{aligned} \tag{S37}$$

The simulated and theoretical approaches to compute the mean and variance is shown in Fig. S4, and the estimates are very similar, up to some simulation error (we used 100 time bins and 5000 replicates in the simulated approach). Note that unlike `RevBayes`, this specific simulation does not include estimation error that accumulates from older branches in the tree, as we fixed $\mathbf{S}(t)$ to be equal for this branch for both the simulated and theoretical approaches. If the whole tree were used to simulate the diversification rate shift history, then we expect that more time bins and/or more replicates would be needed to achieve errors of similar scale.

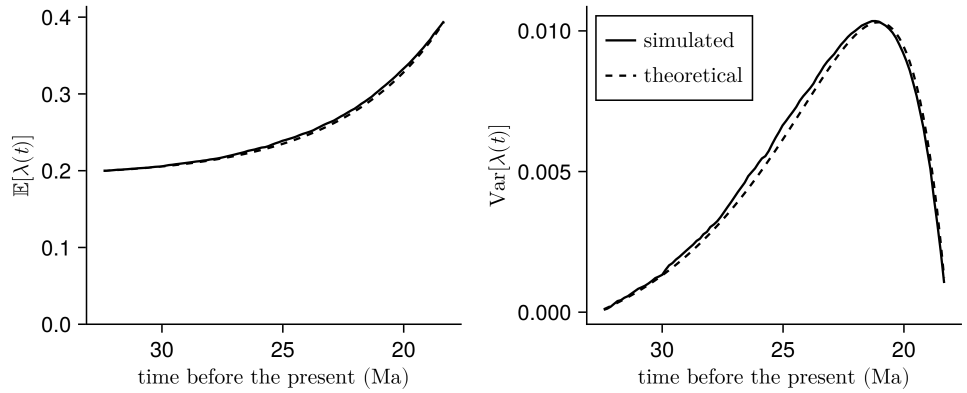

**Figure S4:** Estimates of the posterior mean speciation rate $\lambda(t)$ for the branch in the primates phylogeny that showed the strongest support for a branch-specific diversification rate shift (the branch leading up to the Old World Monkeys clade). The speciation rate categories are $\boldsymbol{\lambda} = [0.1, 0.2, 0.4, 0.1]$, the extinction rate categories are $\boldsymbol{\mu} = [0.05, 0.15, 0.10, 0.2]$, and the diversification shift rate is $\eta = 0.00248$. At the beginning of the branch, the second rate category is the most probable (0.994), whereas the third category is the most probable at the end of the branch (0.968).

Suppose one is interested in calculating the variance of the branch-wide average speciation rate, in addition to the mean (as in the previous section). Conceptually, if $\lambda(t)$ is the discrete random variable for the speciation rate at time $t$, we can imagine the branch-average $\bar{\lambda}$ as a weighted sum of several of time bins, where $t_1, t_2, \ldots$ correspond to the time of the bins

$$\bar{\lambda} = \frac{1}{c}\lambda(t_1) + \frac{1}{c}\lambda(t_2) + \ldots, \tag{S38}$$

where $c$ is the number of time bins. In other words, we have created a new random variable that is a weighted sum of other random variables. Computing the expectation of $\bar{\lambda}$ is straight forward

$$\mathbb{E}[\bar{\lambda}] = \frac{1}{c}\sum_{a=1}^{c} \lambda(t_a). \tag{S39}$$

Computing the variance is more tricky, however, as we also need to take into the account the covariance among the time bins

$$\text{Var}[\bar{\lambda}] = \frac{1}{c^2}\sum_{a=1}^{c} \text{Var}[\lambda(t_a)] + \frac{1}{c^2}\sum_{a=1}^{c}\sum_{b>a}^{c} \text{Cov}[\lambda(t_a), \lambda(t_b)], \tag{S40}$$

where $t_a$ is some time point and $t_b$ is time point younger than $t_a$. The variance $\text{Var}[\lambda(t)]$ we had earlier, but this expression also contains the covariance across different time bins. Assuming that the covariance is approximately zero is not a reasonable assumption, as the variance then converges to zero with increasing number of time bins. Consequently, to theoretically calculate the across-branch variance in speciation rates, we must somehow calculate the covariance across time bins. One way of calculating the covariance is

$$\begin{aligned} \text{Cov}[\lambda(t_a), \lambda(t_b)] = &\sum_{j=1}^{K}\sum_{i=1}^{K} S_j(t_a) \times P_{ij}(t_a, t_b) \\ &\times (\lambda_j - \mathbb{E}[\lambda(t_a)]) \times (\lambda_i - \mathbb{E}[\lambda(t_b)]), \end{aligned} \tag{S41}$$

where $S_j(t_a)$ represents the marginal probability of the rate category being $j$ at time $t_a$, and $P_{ij}(t_a, t_b)$ is the transition probability from category $j$ at time $t_a$ to the category $j$ at a younger time $t_b$.

Specifically, $P_{ij}(t_a, t_b)$ is calculated by setting $\mathbf{F}(t_a)$ to a one-hot vector, and numerically integrating with $d\mathbf{F}/dt$ to a younger time $t_b$, and then element-wise multiplying with $\mathbf{D}(t_b)$, all repeated for all possible one-hot vectors of $\mathbf{F}(t_a)$, and assembling them into a matrix $\mathbf{P}$. Finally, we normalize the matrix $\mathbf{P}$ such that the columns sum to one. Note that we can not use Eq. (S18) to calculate $P_{ij}(t_a, t_b)$, as Eq. (S18) is only a good approximation for very short time intervals $t$ to $t + \Delta t$. All taken together, the expression for the variance becomes

$$\begin{aligned} \text{Var}[\bar{\lambda}] = \frac{1}{c^2}\sum_{a=1}^{c} \text{Var}[\lambda(t_a)] + \frac{1}{c^2}\sum_{a=1}^{c}\sum_{b>a}^{c}\sum_{j=1}^{K}\sum_{i=1}^{K} &S_j(t_a) \times P(Z(t_b) = i | Z(t_a) = j) \\ &\times (\lambda_j - \mathbb{E}[\lambda(t_a)]) \times (\lambda_i - \mathbb{E}[\lambda(t_b)]). \end{aligned} \tag{S42}$$

When comparing $\text{Var}[\bar{\lambda}]$ between the simulated, stochastic mapping approach and the theoretical approach (Fig. S5), we can see that the theoretical approach converges when more time bins are used, and 20 or 30 time bins are perhaps sufficient for a reasonable approximation. One caveat of the example in Fig. S5 is, however, that it was made with a toy model that only has four diversification rate categories. In a more typical setting, where we want more robust estimates of the branch-specific diversification rates, we would like to use more diversification rate categories, for example $n^2 = K = 36$ or $n^2 = K = 100$. The equation to calculate $\text{Var}[\bar{\lambda}]$ contains four nested sums, and we expect that for a larger model it will become cumbersome to calculate the variance in this way. Therefore, if the across-branch variance in diversification rates is of interest, we recommend to use the stochastic mapping approach as used in Höhna et al. (2019).

### S6 The likelihood surface

The likelihood surface, when plotted as a function of $\hat{\lambda}$ and $\hat{\mu}$, has several ridges with local optima (Fig. S6). This type of surface can be difficult to traverse using numerical methods. Optimization algorithms can find a local maximum instead of the global maximum likelihood parameter values. Similarly, Markov chain Monte Carlo samplers can have a difficult time switching between the ridges. This means that it is difficult to sample the posterior distribution, and in practice it can lead to problems with convergence. In `Pesto` we used a two-step approach to find estimates for $\hat{\lambda}$, $\hat{\mu}$ and $\eta$. Based on our exploration of the likelihood surface for the primates tree, this will not lead to the global maximum likelihood parameter estimates, but instead something that is reasonably close (Fig. S6, white dot), within a few log-likelihood units of the global maximum.

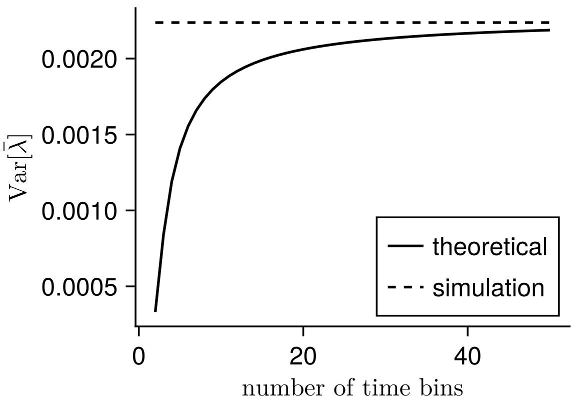

**Figure S5:** The posterior variance of the across-branch mean speciation rate, using i) a simulated, stochastic mapping approach, and ii) a theoretical approach that uses a time-bin approximation. The units of $\mathrm{Var}[\bar{\lambda}]$ are "number of speciation events per lineage per Ma" squared. When sufficiently many number of time bins are used, then the variance converges.

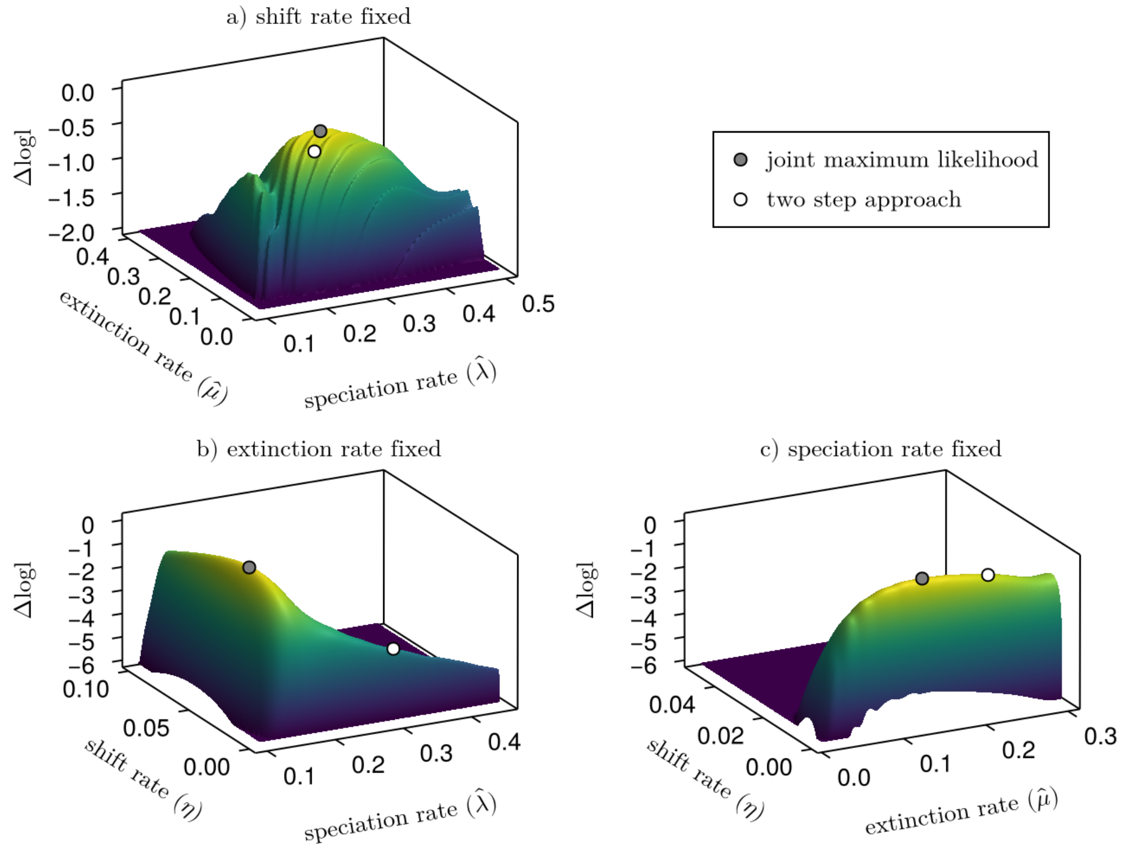

**Figure S6:** The likelihood surface of the birth-death-shift model as a function of the three parameters $\hat{\lambda}$, $\hat{\mu}$ and $\eta$. From $\hat{\lambda}$, $\hat{\mu}$, we take 10 quantiles from the distributions LogNormal($\log(\hat{\lambda})$, sd = 0.587) and LogNormal($\log(\hat{\mu})$, sd = 0.587), and use all pairwise comparisons to form $\boldsymbol{\lambda}$ and $\boldsymbol{\mu}$. The log likelihoods are offset such that the maximum is zero. The data is the primates phylogeny. The three panels depict grid searches where we held one parameter constant in each panel. In panel a) we set $\eta = 1/\sum_i b_i$, where $b_i$ is the length of branch $i$. In b) we set $\hat{\mu} = 0.22$, and in c) we set $\hat{\lambda} = 0.30$ (the maximum-likelihood parameter values under the constant-rate birth-death process). Notice that the parameter values inferred using the two-step approach is within a few log-likelihood units of the joint maximum likelihood approach, however they are not identical.

### S7 Estimation error in branch-specific diversification rates

In the main text, we showed the proportional error in the inference of branch-specific speciation rates. Here, we show the equivalent in branch-specific extinction rates Fig. S7. The figure shows that most proportional errors are between 0.33 and 3.0, meaning that branch-specific extinction rate estimates are underestimated by a third, or overestimated by a factor of three. However, as the tree size increases, the proportional errors converge to approximately zero, meaning that the method is approximately unbiased. We also calculated branch-specific estimation errors for net-diversification rates. Since branch-specific net-diversification rates are often negative, however, it made less sense to use the geometric mean as a summary method per tree. For this reason, we plot the arithmetic mean of the branch-specific estimation errors in Fig. S8.

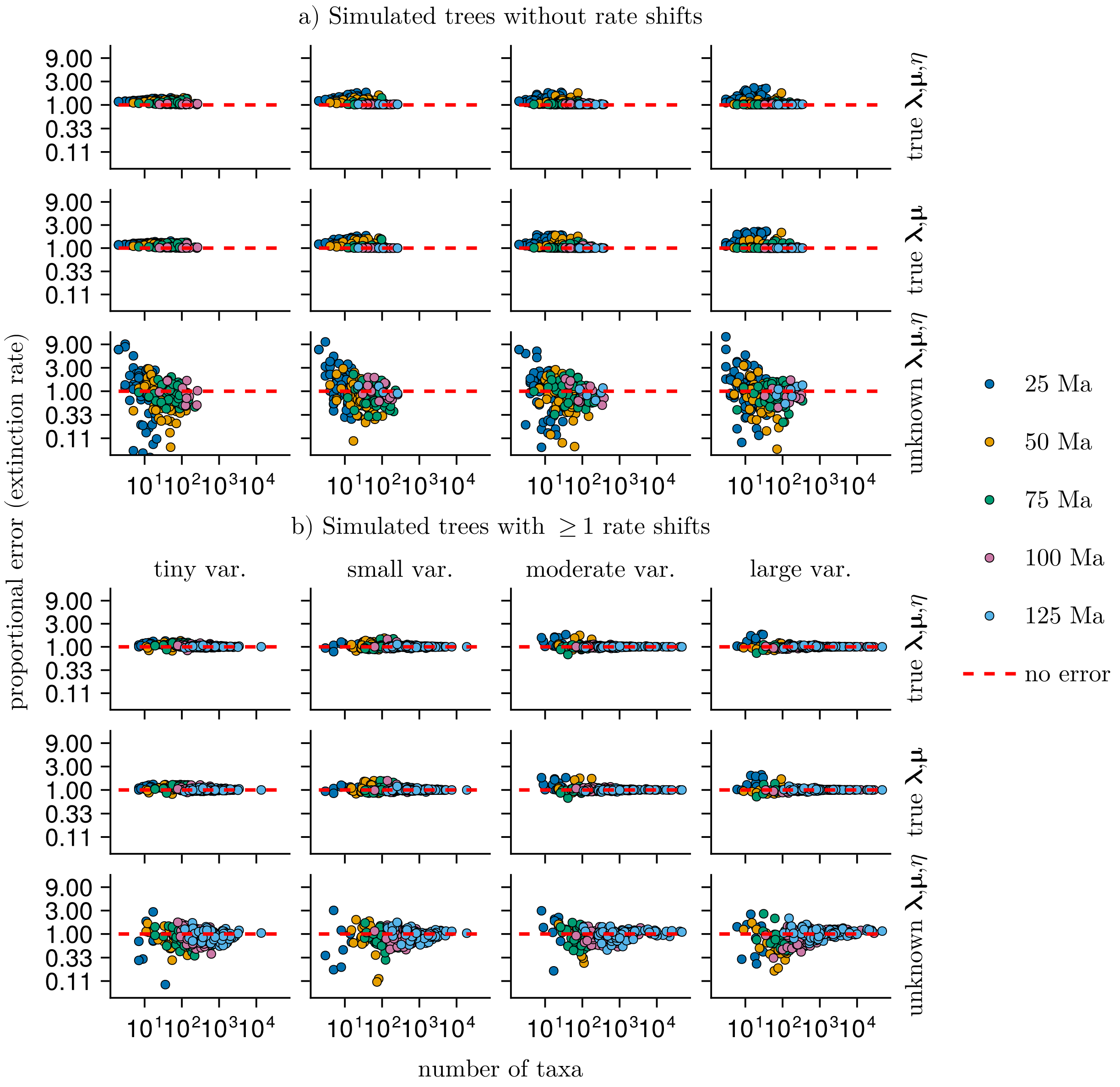

**Figure S7:** The estimation error in branch-specific extinction rates, as proportional errors. For each tree simulated tree, we calculated the proportional errors per branch, and averaged across the tree: $\exp(\frac{1}{n_{\text{branches}}} \sum_M \log(\bar{\mu}_M) - \log(\bar{\mu}_{M,\text{true}}))$, where $\bar{\mu}_M$ is the estimate for the average extinction rate on branch $M$. A value of one means that the branch-specific extinction rates are unbiased, on average across the tree. Larger than one means overestimation, and less than one means underestimation. The proportional errors for extinction rates are slightly larger than those for speciation rates.

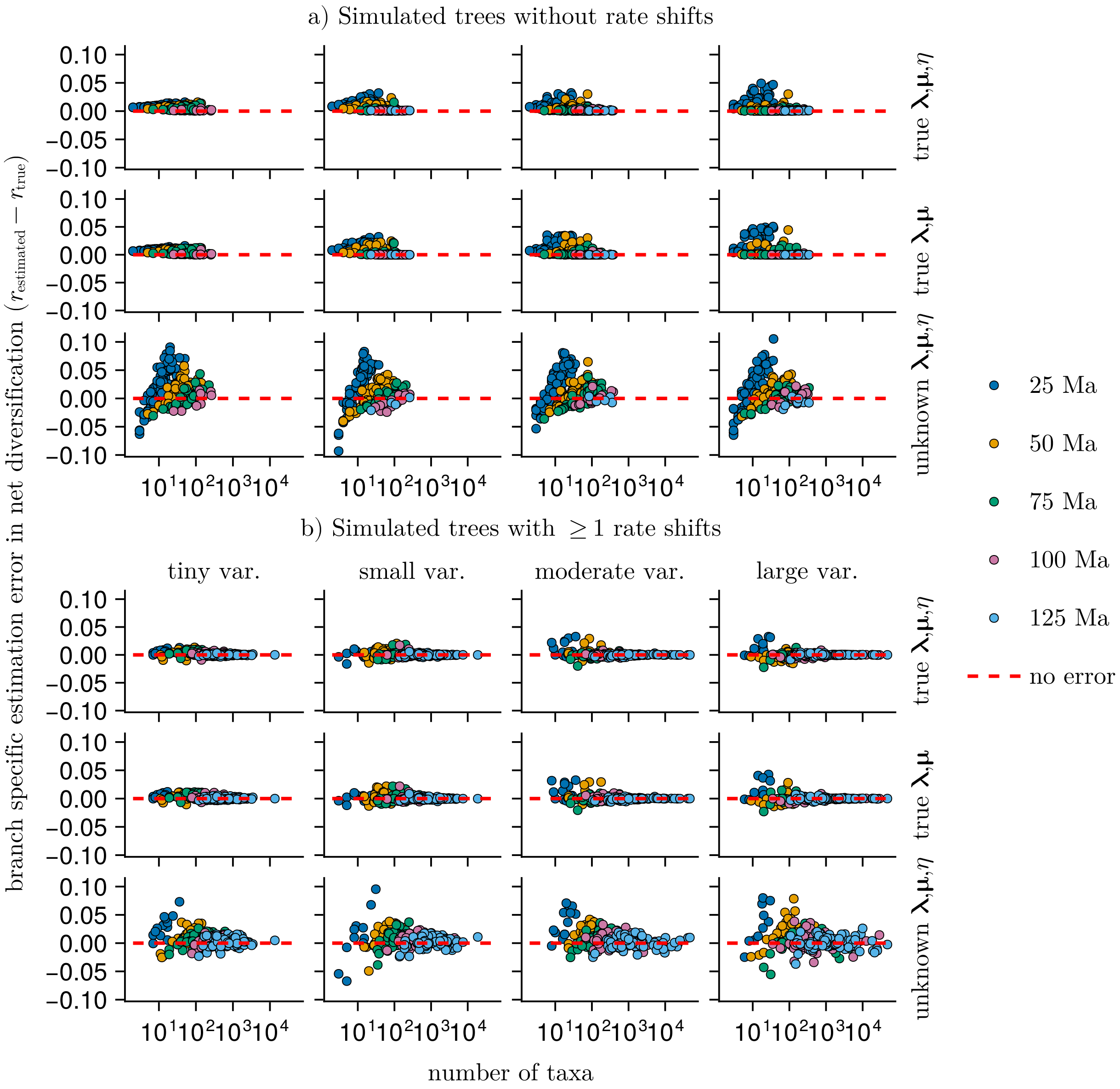

**Figure S8:** The estimation error in branch-specific net-diversification rates. For each tree simulated tree, we calculated the estimation errors per branch, and averaged across the tree: $\frac{1}{n_{\text{branches}}} \sum_M (\bar{r}_M - \bar{r}_{M,\text{true}})$, where $\bar{r}_M$ is the estimate for the average net-diversification rate on branch $M$. Note that this plot shows arithmetic means, unlike geometric means in the other figures of estimation error. A value of zero means that the branch-specific extinction rates are unbiased, on average across the tree. Larger than zero means overestimation, and less than zero means underestimation. Seven data points with an error of about $-0.61$ are omitted from the figure due to the choice of y-axis limits.

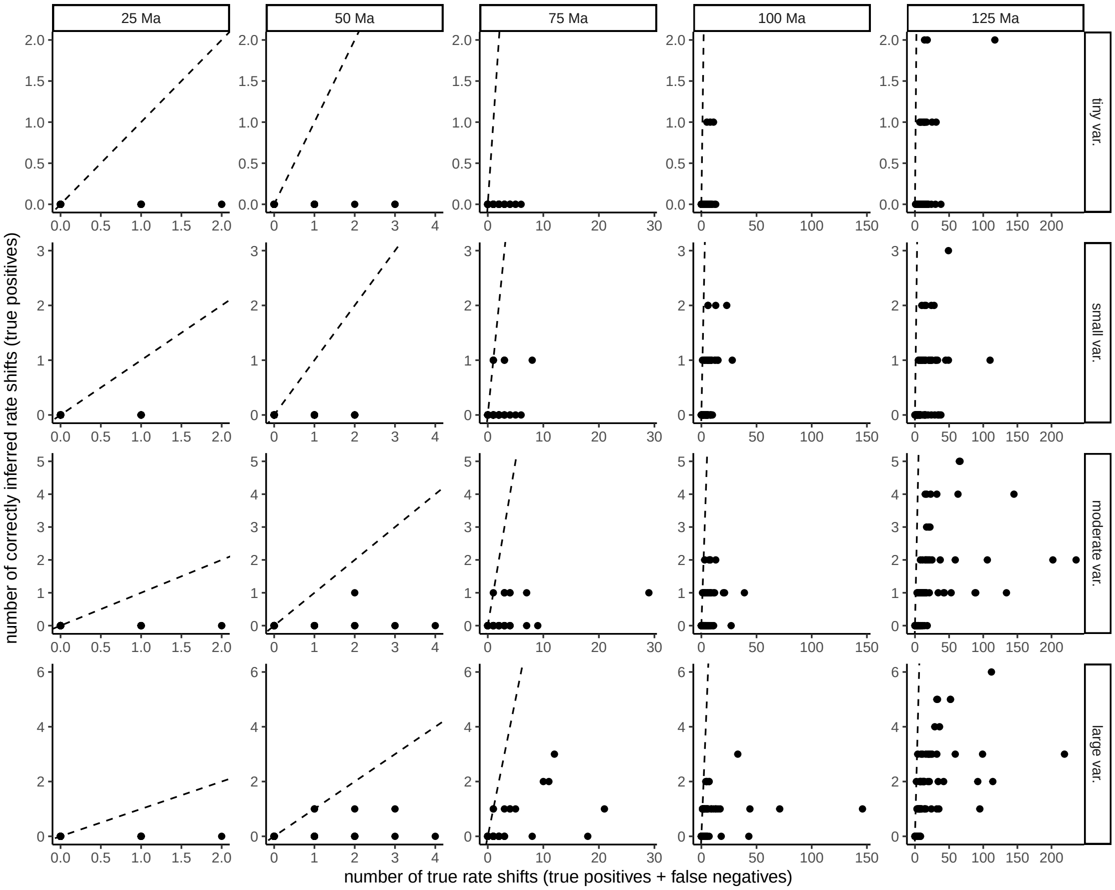

**Figure S9:** The number of correctly inferred rate shifts (i.e. true positives) versus the number of rate shifts that happened (true positives + false negatives) in the simulated trees. Each point represents one phylogeny. The criteria for that a shift was inferred was that the Bayes factor had to be larger than 10, and $\hat{N}$ had to be larger than 0.5. The panels are split by simulation time (from 25 to 125 Ma) and rate variation in the true model (i.e. models A to D in Table 1). The dashed lines represent the one-to-one relationship.

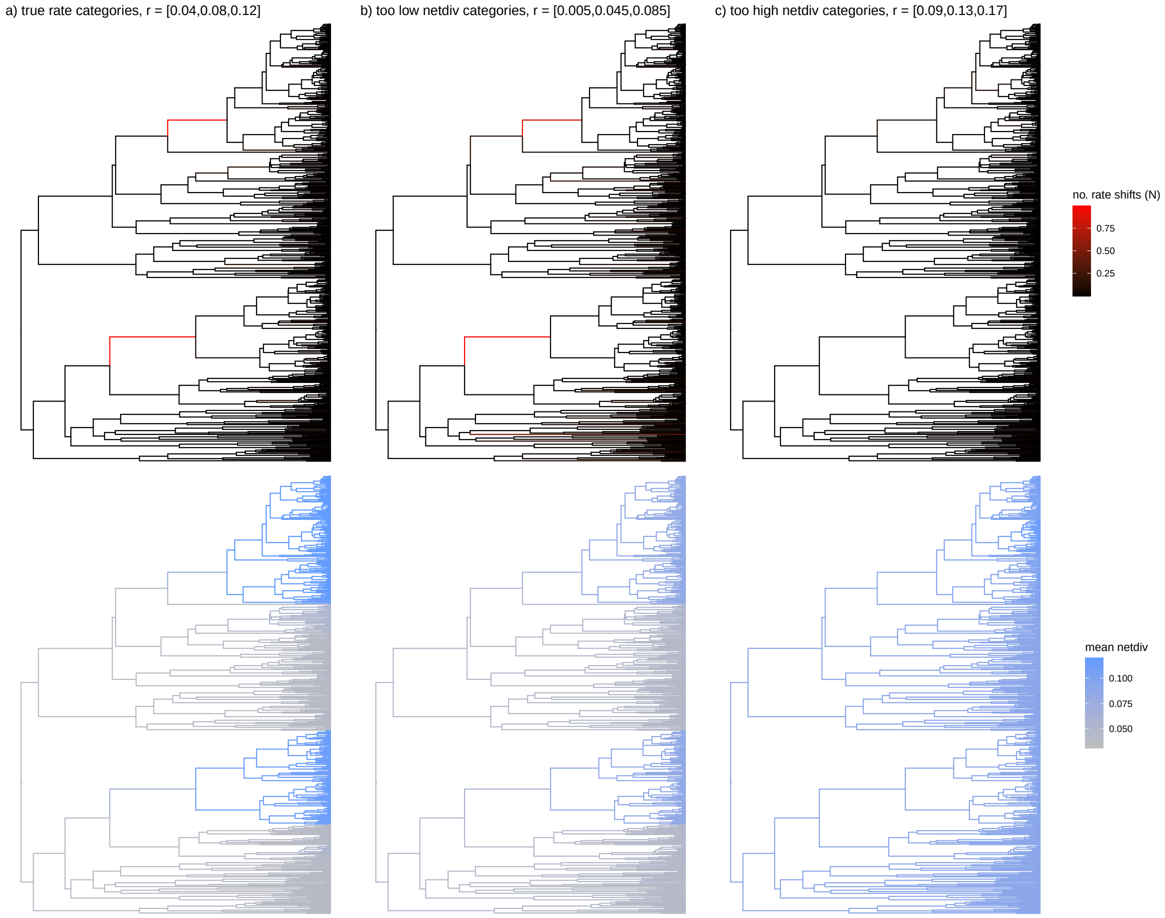

**Figure S10:** A brief example of what the result might be if the rate categories are mis-specified. To emulate this, we set the net-diversification rates to a) their true values, b) too low and c) too high. The relative extinction (i.e. $\mu/\lambda = 2/3$) is equal across all three models and across the rate categories. The phylogeny is the same that was used in the main text in Fig. 9b, simulated from the model with large rate variation. In all three models we estimated the shift rate using maximum likelihood. In a) and b), the two rate shifts that led to the radiation of species-rich clades are recovered. However, in b) the net-diversification rates in the two in-groups are underestimated, and there is one false positive. In c) the shift rate estimate is about zero, no shifts are recovered, and the net-diversification rates are overestimated in what would be the ”outgroup”.

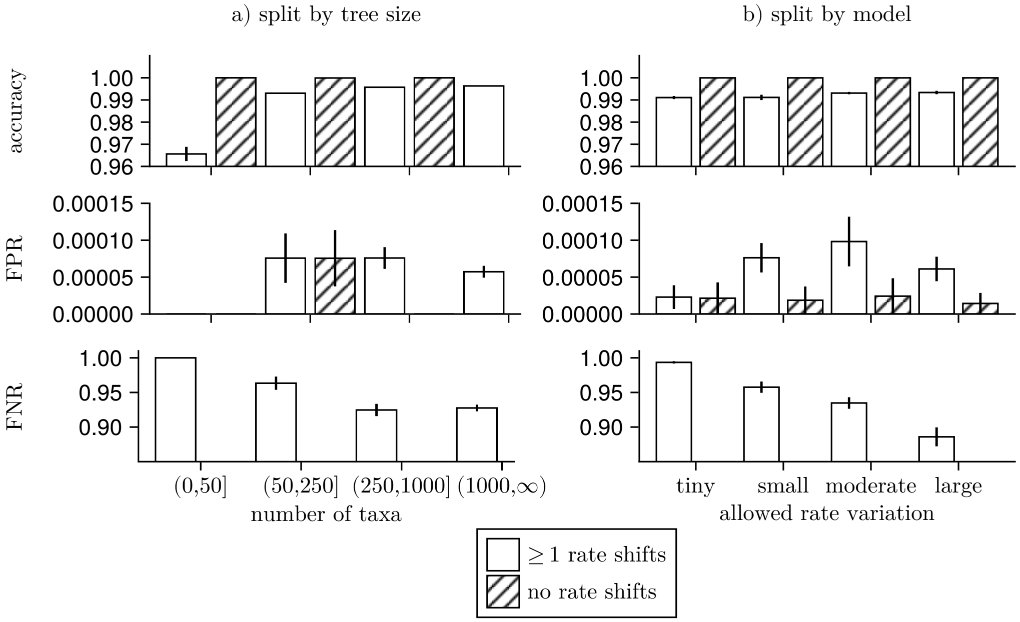

**Figure S11:** Equivalent to Fig. 10 in the main manuscript, however with the y-axis limits chosen on a much smaller interval than {0,1}, better show-casing the variation within each panel.

### References

Barido-Sottani, J., Vaughan, T. G., and Stadler, T. (2020). A Multitype Birth–Death Model for Bayesian Inference of Lineage-Specific Birth and Death Rates. *Systematic Biology*, 69(5):973–986.

Dormand, J. R. and Prince, P. J. (1980). A family of embedded Runge-Kutta formulae. *Journal of computational and applied mathematics*, 6(1):19–26.

Felsenstein, J. (1981). Evolutionary trees from dna sequences: a maximum likelihood approach. *Journal of molecular evolution*, 17:368–376.

Felsenstein, J. (2004). *Inferring phylogenies*. Sinauer Associates, Sunderland, MA.

FitzJohn, R. G. (2012). Diversitree: comparative phylogenetic nalyses of diversification in R. *Methods in Ecology and Evolution*, 3(6):1084–1092.

Gallager, R. (1962). Low-density parity-check codes. *IRE Transactions on information theory*, 8(1):21–28.

Höhna, S., Freyman, W. A., Nolen, Z., Huelsenbeck, J. P., May, M. R., and Moore, B. R. (2019). A Bayesian Approach for Estimating Branch-Specific Speciation and Extinction Rates. *bioRxiv*.

Höhna, S., Heath, T. A., Boussau, B., Landis, M. J., Ronquist, F., and Huelsenbeck, J. P. (2014). Probabilistic graphical model representation in phylogenetics. *Systematic biology*, 63(5):753–771.

Lewis, P. O. (2001). A likelihood approach to estimating phylogeny from discrete morphological character data. *Systematic biology*, 50(6):913–925.

Louca, S. and Pennell, M. W. (2020). A general and efficient algorithm for the likelihood of diversification and discrete-trait evolutionary models. *Systematic Biology*, 69(3):545–556.

Maddison, W. P., Midford, P. E., and Otto, S. P. (2007). Estimating a binary character's effect on speciation and extinction. *Systematic Biology*, 56(5):701.

Martínez-Gómez, J., Song, M. J., Tribble, C. M., Kopperud, B. T., Freyman, W. A., Höhna, S., Specht, C. D., and Rothfels, C. J. (2024). Commonly used bayesian diversification methods lead to biologically meaningful differences in branch-specific rates on empirical phylogenies. *Evolution Letters*, 8(2):189–199.

May, M. R. and Rothfels, C. J. (2023). Diversification models conflate likelihood and prior, and cannot be compared using conventional model-comparison tools. *Systematic Biology*, 72(3):713–722.

Nielsen, R. (2002). Mapping mutations on phylogenies. *Systematic biology*, 51(5):729–739.

Pagel, M. (1994). Detecting correlated evolution on phylogenies: a general method for the comparative analysis of discrete characters. *Proceedings of the Royal Society of London. Series B: Biological Sciences*, 255(1342):37–45.

Pearl, J. (1982). Reverend Bayes on Inference Engines: A Distributed Hierarchical Approach. In *Proceedings of the National Conference on Artificial Intelligence*, pages 133–136, Pittsburgh, PA. AAAI.

Pearl, J. (1988). *Probabilistic Reasoning in Intelligent Systems: Networks of Plausible Inference*. Morgan Kaufmann Publishers, San Francisco, CA, 1st edition.

Rabosky, D. L. (2014). Automatic detection of key innovations, rate shifts, and diversity-dependence on phylogenetic trees. *PLoS One*, 9(2):e89543.

Rackauckas, C. and Nie, Q. (2017). DifferentialEquations.jl – a performant and feature-rich ecosystem for solving differential equations in Julia. *Journal of open research software*, 5(1).

Russell, S. J. and Norvig, P. (2016). *Artificial intelligence: a modern approach*. Pearson.

Teo, B., Bastide, P., and Ané, C. (2025). Leveraging graphical model techniques to study evolution on phylogenetic networks. *Philosophical Transactions B*, 380(1919):20230310.

Tsitouras, C. (2011). Runge–Kutta pairs of order 5(4) satisfying only the first column simplifying assumption. *Computers & Mathematics with Applications*, 62(2):770–775.

Yang, Z., Kumar, S., and Nei, M. (1995). A new method of inference of ancestral nucleotide and amino acid sequences. *Genetics*, 141(4):1641–1650.
